# Regulatory stochasticity drives opposing phenotypic outcomes in cell-fate decision networks

**DOI:** 10.64898/2026.08.24.746658

**Authors:** Kishore Hari, Abhay Gupta, Lakshmi Malvadi Shivakumar, Prakash Kulkarni, Ravi Salgia, Mohit Kumar Jolly, Herbert Levine

## Abstract

Models of gene regulatory networks (GRNs) underlying cell-fate decision systems treat regulatory parameters as fixed quantities, although transcription-factor efficacy varies with molecular context. We asked how temporal fluctuations in interaction strength, rather than molecular abundance, alter phenotype occupancy in multistable regulatory networks. We applied three stochastic update rules to fold-change parameters—independent fluctuations anchored to their initial values, a bounded additive random walk, and a bounded multiplicative random walk—across RACIPE ensembles of mutually inhibitory circuits, and tested analogous perturbations in Boolean models of epithelial–mesenchymal plasticity and gonadal-fate determination. Anchored fluctuations largely preserved deterministic occupancies of network states. Additive fluctuations increased occupancy of all-high co-expression states, particularly in multistable regimes and when high expression was readily accessible; this effect extended to hybrid team-expression states in both biological networks. Multiplicative fluctuations biased inhibitory fold changes toward stronger repression and favored single-high states in a topology-dependent manner. Fixed-parameter controls sampled from the same noise-induced distributions did not fully reproduce the additive effect or the multiplicative effect in several circuits. Our analysis traced the additional additive bias to an asymmetry between unconditional derepression and conditional repression. Thus, temporally evolving regulatory parameters can redistribute phenotype occupancy in ways not predicted from static parameter ensembles, with outcomes jointly determined by fluctuation dynamics and network topology.

## 1 Introduction

GRNs are biological regulatory circuits that constitute a complex collection of molecular regulators including transcription factors (TFs), signaling molecules, and DNA segments that interact to regulate gene expression. They determine the spatiotemporal patterns of gene expression involved in determining cellular development, identity, and environmental responses. They also play a central role in determining the organism’s body plan, which is central to evolutionary developmental biology [1, 2, 3]. GRNs are conventionally modeled as deterministic systems with fixed interaction parameters although it is clear that the effective strength of any regulatory interaction is subject to continuous fluctuation in vivo. Nonetheless, an overwhelming fraction (≈90%) of TFs are intrinsically disordered proteins (IDPs) [4] with significant disordered regions [5, 6]. These disordered regions mediate conformationally heterogeneous, post-translational modification (PTM)-sensitive contacts with DNA and co-factors, causing regulatory affinities to vary dynamically rather than assume a fixed value . Therefore, understanding IDP conformational noise is paramount for understanding cell-fate specification. Although the individual conformers in an IDP ensemble are in fast exchange, certain conformations are stabilized by specific PTMs, especially phosphorylation, that exist with half-lives that range from minutes to hours, contributing to “conformational noise” [7, 8]. Non-specific TF binding to genomic decoy sites further modulates the effective regulatory signal in an affinity- and abundance-dependent manner [9, 10]. Enhancer dynamics and transcriptional bursting add a third layer, coupling stochastic TF occupancy at cis-regulatory elements directly to fluctuations in target gene activation [11, 12]. The availability of TF binding sites can also vary over the course of transcription, due to noise in chromatin folding states [13]. Collectively, these mechanisms argue that the parameters governing regulatory interactions are not static; instead, they are stochastic quantities. Consequently, the impact of such regulatory noise—temporal stochasticity in these parameters—on the dynamics of multistable GRN motifs underlying cell-fate decisions remains poorly understood.

This question is especially relevant in the context of cancer, where genetic instability, aberrant chromatin remodeling, and rewired signaling cascades collectively amplify both intrinsic gene expression noise and extrinsic cell-to-cell variability [14, 15, 16]. Non-genetic heterogeneity — stable phenotypic variation within a clonal population arising from gene expression noise and network multi-stability — is now recognized as a major contributor to drug tolerance and tumor progression [16, 17, 18]. In this setting, the parameters of regulatory interactions that govern the dynamics of non-genetic heterogeneity cannot be considered fixed: PTM landscapes, co-factor availability, and chromatin accessibility to TFs all fluctuate, effectively introducing stochasticity into the fold-change parameters that govern network behavior.

Perhaps a natural starting point to studying how regulatory noise propagates through GRNs is the toggle switch — a simple mutually repressive circuit between two master regulators that is among the most commonly occurring and extensively characterized motifs for binary cell-fate decisions [19]. The toggle switch exhibits bistability under a broad range of kinetic parameters, and noise-induced switching between its two attractors has been studied in detail in both synthetic and natural contexts [20]. Many biological decisions, however, are not binary. For example, T helper cell differentiation, haematopoietic lineage commitment, and epithelial–mesenchymal–hybrid state transitions all involve three or more stable phenotypes. The addition of self-activation to the toggle switch (TSSA, toggle switch with self-activation) stabilizes the intermediate phenotype — corresponding to hybrid phenotypes in EMT and progenitor states in differentiation cascades — where both nodes are co-expressed simultaneously [21]. Three nodes connected to each other via a toggle switch, forming the toggle triad (TT). TT enables three distinct cell states, each defined by one node being highly expressed and the other two repressed [22]. Importantly although GRNs underlying cell-fate decision systems often involve a larger number of TFs than accounted for in these motifs, TFs corresponding to a phenotype often form teams such that nodes belonging to the same team activate each other. These teams mutually inhibit each other, thereby forming higher order forms of the motifs listed above. These higher order motifs have similar qualitative behavior as their small motif counterparts such as a two team network promoting bistability with a small fraction of intermediate states, similar to TSSA.

The effect of stochasticity on toggle switch dynamics has been studied extensively, though almost exclusively through noise applied to molecular copy numbers rather than interaction parameters. Early work established that intrinsic noise — arising from the discreteness of protein and mRNA molecules — can drive spontaneous switching between the attractors of a multistable circuit, with switching rates that decrease exponentially with copy number and increase with noise magnitude [20, 23]. Analytical treatments using the Chemical Langevin Equation and mean first-passage time theory have characterized how the depth and asymmetry of the potential wells governing each attractor determine residence times and transition rates [24, 25, 26]. A key finding across these studies is that noise type — not only noise magnitude — shapes attractor occupancy: additive and multiplicative noise applied to the same deterministic system produce different effective potential landscapes and therefore different stationary probability distributions [27]. Extending beyond Gaussian noise, Lévy noise models have shown that heavy-tailed fluctuations can induce coherent switching at substantially lower intensities than Gaussian noise requires, by enabling large jumps that entirely overcome the potential barrier [28]. At the network level, toggle switch dynamics has been studied under coupled and multicellular settings, where intracellular multiplicative noise can induce synchronized switching across cell populations via quorum-sensing pathways [23]. More recently, the sRACIPE framework demonstrated that expression-level (state) noise merges bistable clusters at high intensity, while static parametric variation across the model ensemble instead broadens clusters without destabilizing them [29]. In both cases, however, the kinetic parameters themselves remain fixed once sampled; neither treats the parameter as a time-evolving stochastic process, which is the gap that our regulatory noise framework addresses. Stochastic analysis of the toggle triad and related tristable motifs has received comparatively little attention, with existing studies focused primarily on deterministic bifurcation structure and attractor geometry [22, 30] rather than noise-driven state transitions. Whether the richer transition topology of the toggle triad — permitting direct switching between all three attractor pairs — confers qualitatively different susceptibility to regulatory noise compared to the toggle switch remains an open question.

In this manuscript, we study how regulatory noise – that is, the fluctuations of the parameters governing the interactions in GRNs – systematically shapes the dynamics of these motifs. We focus on the fold-change parameter [31], which describes the change in the production rate of a target node due to an upstream transcription factor. We couple stochastic fold-change perturbations with the Random Circuit Perturbation (RACIPE) framework [31]. RACIPE generates large ensembles of ODE models with randomly sampled kinetic parameters, enabling topology-level inferences that are robust to parameter uncertainty. By superimposing continuous stochastic noise on the fold-change parameters of these ensembles, we successfully isolated the effect of regulatory noise from parameter-set-specific idiosyncrasies and characterized how mean residence times and switching frequencies depend on noise magnitude, noise mode, and attractor stability class. Our results reveal that regulatory noise is sufficient to induce state transitions that are inaccessible in the deterministic limit, and that bistable and tristable parameter regimes differ qualitatively in their susceptibility to such noise. Therefore, our results demonstrate that the type of regulatory noise, not just its magnitude, is a key determinant of phenotypic plasticity in cell-fate circuits.

## 2 Results

### 2.1 Simulating regulatory noise in GRN motifs underlying cell-fate decision

We begin by analyzing the binary and tertiary cell-fate decision motifs — the toggle switch (TS) and the toggle triad (TT) and their extensions that include self-activation edges (TSSA and TTSA). A toggle switch consists of two mutually inhibiting transcription factors (Figure 1A(i-ii)), whereas a toggle triad consists of three mutually inhibiting transcription factors (Figure 1A(iii-iv)). TS represents the core motif underlying binary cell-fate decision networks. The phenotypic landscape emergent from TS is dominated by two states with mutually exclusive expression patterns - one of the nodes has low expression while the other is high - (0, 1) or (1, 0). Under appropriate parametric conditions, the toggle switch also exhibits bistability, enabling transitions between (1, 0) and (0, 1) states [19]. The addition of self-activation (TSSA) stabilizes the (1, 1) state in which both nodes are co-expressed. This co-expression state, often referred to as a “hybrid” state, has been associated with stemness in development [32] and with high plasticity and pro-metastatic features in cancer [33, 34, 35].

**Figure 1:**
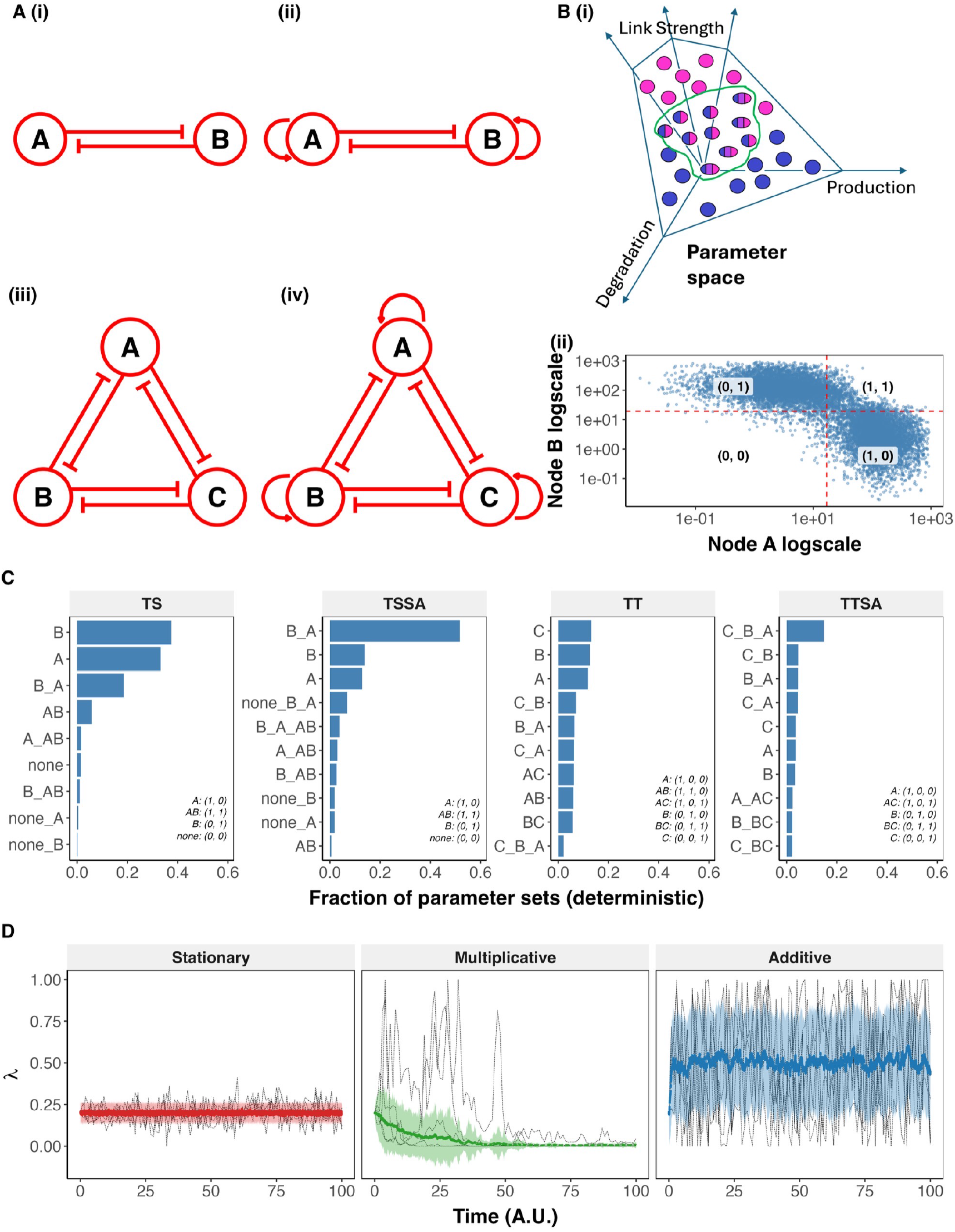
RACIPE network topologies and the stochastic simulation framework.. **(A)** Network topology diagrams for the four base circuits used throughout this study: (i) TS (toggle switch), (ii) TSSA (toggle switch with self-activation), (iii) TT (toggle triad), (iv) TTSA (toggle triad with self-activation). Red circles denote nodes; bar-headed edges denote inhibitory interactions; curved arrows denote self-activation. **(B)** (i) Schematic of the RACIPE parameter-sampling framework: production, degradation, and interaction-strength parameters are randomly sampled within a bounded parameter space, generating an ensemble of models with varying dynamical behavior. (ii) Steady-state expression of node A vs. node B (log scale) across the RACIPE parameter ensemble for the TS network. Dashed red lines mark each gene’s discretization threshold (mean of its expression in log scale across all steady states); quadrants are labeled with the corresponding discrete state, (A, B). **(C)** Frequency of the 10 most frequent discrete attractors found by RACIPE for each network (TS, TSSA, TT, TTSA), expressed as the fraction of parameter sets that map onto these attractors. States are represented in short-hand form, with the key given inside the plot. Compound labels (e.g. “A_B”) denote multistable parameter sets. Because TT and TTSA support many more distinct multistable combinations than TS/TSSA, the top-10 bars shown capture only part of the full ensemble: 100%, 99.2%, 77.5%, and 46.2% of all parameter sets for TS, TSSA, TT, and TTSA respectively (out of 9, 12, 97, and 119 total distinct attractor combinations found). **(D)** Simulated trajectories of the fold-change parameter *λ*(*t*) under the three stochastic noise-update rules used in this study (Stationary, Multiplicative, Additive), shown for an inhibitory-edge-range parameter (*λ* ∈ [0, 1], *λ*_0_ = 0.2). Thin black dashed lines show 5 representative individual trajectories (sampled once per time unit for legibility); the colored line and shaded ribbon show the mean *±* SD across 100 simulated trajectories.

While binary cell-fate decision systems dominate biology owing to their stability and robustness to internal and external perturbations [36, 37], some instances of tertiary cell-fate systems — such as T-cell differentiation — employ a toggle triad architecture. A toggle triad consists of three mutually inhibiting transcription factors (Figure 1A(iii)) and permits three differentiated, single-high states where one of the three nodes has a high expression while the other two have low expression ((0, 0, 1), (1, 0, 0), and (0, 1, 0)) as well as three hybrid, double-high states with two out of three nodes having high expression (e.g. (1, 1, 0)). Parameter sweeps of the toggle triad result predominantly in monostable and bistable configurations, with a relatively smaller fraction exhibiting tristability. Single network architectures that allow the co-existence of four or more mutually exclusive states are not currently known [38].

We model these networks using a Hill-equation kinetics-based ODE system (Equation 1, 2, Section 4.1).

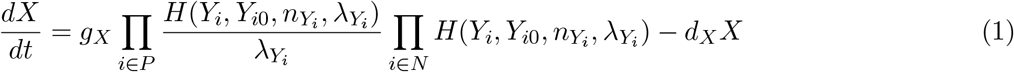

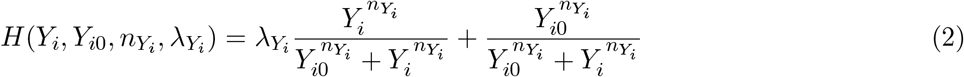

Briefly, these equations combine transcription and translation into a single production term and a first-order degradation term representing mRNA and protein turnover. The production term consists of a rate constant (*g*_*X*_) representing polymerization of RNA and protein, together with a combination of shifted Hill functions that formalize transcriptional regulation by input nodes. Input nodes change the production rate via the fold-change parameter *λ*: for inhibitory inputs, *λ <* 1, so that the presence of the regulator reduces the production rate; for activating inputs, *λ >* 1. The *λ* of inhibitory links is bounded between 0.01 (strongest inhibition) and 1 (no inhibition) and that of the activating links is bounded between 100.0 (strongest activation) and 1.0 (no activation).

We first simulated these networks deterministically using RACIPE [31], a tool that allows us to explore the different configurations that the emergent phenotypic landscape can take across a large parameter space. RACIPE generates a large ensemble of ODE models by sampling multiple parameter sets (Figure 1B) and simulates each ODE model from multiple initial conditions until steady states are reached, resulting in an ensemble of steady states. We discretized expression of each node in each steady state into high (1) or low (0) based on the node’s log-average expression across the ensemble of steady states (Figure S1B). Threshold values for each network are given in Table S1. The models generated by RACIPE can be classified as monostable or bistable based on the number of discrete steady states they support, and further subdivided by the specific attractor configuration (see Table S2). For the toggle switch, the dominant attractors were (0, 1), (1, 0), and the bistable (0, 1)–(1, 0), as expected (Figure 1C, i). Although a small fraction of models do permit (1, 1) as a steady state, the corresponding fold-change parameters are always close to 1, rendering inhibitions ineffective [39]. TSSA enhances bistability in the parameter space, with the other dominant attractors being (0, 1) and (1, 0). Additionally, TSSA also supports tristability with (0, 1) and (1, 0) and either (0, 0) or (1, 1). The (0, 0) state was accessed by a very small number of initial conditions, while (1, 1) was more accessible (Figure 1C, ii). TT and TTSA support single-high (one of the nodes has a high expression - (0, 0, 1) etc.) and double-high (two of the nodes have high expression - (0, 1, 1) etc.) nearly equally. TTSA enhances multistability over TT, often exhibiting a tristable attractor with three single-high states (Figure 1C, iii, iv). We explored three noise modes, described by the equations below:

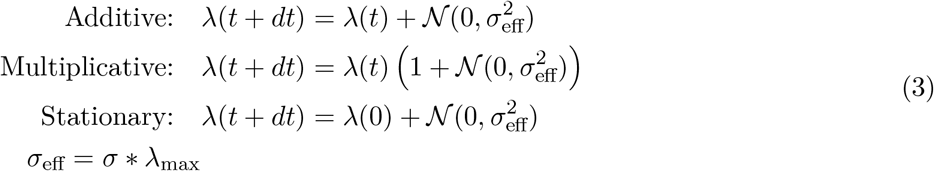

Networks were simulated at eight levels of *σ* : 0.0, 0.001, 0.005, 0.01, 0.05, 0.1, 0.5 and 1.0, scaled by the upper bound of *λ* (Eq. 3, *λ*_*max*_ = 1.0 for inhibitory edges and *λ*_*max*_ = 100.0 for excitatory edges). The *σ* values were chosen to ensure a larger coverage of the low to moderate noise region (*σ* ≤ 0.1), as that region likely represents the underlying biology better than unrealistically high noises that span 50 or 100% of the parameter range (*σ* ∈ [0.5, 1.0]). The higher *σ* values serve a more theoretical purpose in helping us understand the effect of stochastic simulations. The three noise modes produce qualitatively distinct trajectories of *λ* over time. Stationary noise models normally distributed fluctuations around the initial value of *λ* (Figure 1C, i, S1B, i). Additive noise models a constrained random walk of *λ* bounded by the corresponding limits for activating and inhibiting *λ* values. The resultant stationary distribution has a mean of 0.5 for inhibitory links and 50 for activatory links (Figure 1C, ii, S1B, ii). Multiplicative noise scales *σ* by the current *λ* value, such that lower noise values show lower deviation and higher noise values show higher deviation (Figure 1C, iii, S1B, iii). The consequences of these distinct *λ* trajectories for network state dynamics are explored in the following sections.

We sampled monostable and bistable models separately from RACIPE, weighted by the relative frequencies of each attractor configuration. We then performed stochastic simulations of these networks by periodically perturbing *λ*. For each parameter set, we generated 100 stochastic trajectories starting from randomly chosen initial conditions. We use the RACIPE ensemble-derived thresholds (Figure 1B, ii, S1) to discretize the stochastic trajectories as well. Based on the number of discretized steady states allowed by a given parameter set in the deterministic simulations, each parameter set was characterized as monostable (single steady state), bistable, tristable and so on. For each network, we sampled up to 1000 parameter sets per stability class (fewer where a stability class had fewer available parameter sets) and for each parameter set, we performed 100 stochastic simulations each to estimate the statistics.

### 2.2 High levels of stationary noise pushes the system towards hybrid states

Stationary noise mimics the classical assumption underlying GRN models, that the fitted parameters are the mean of a distribution whose variance does not significantly change the emergent phenotypic landscape from what the mean alone predicts. Thus, we started with analyzing the stochastic simulations under stationary noise to evaluate the validity of the deterministic representation of regulatory parameters. We analyzed the stochastic trajectories by first discretizing them using the RACIPE ensemble threshold values (Figure 2A). The resultant discrete state trajectories are then used to calculate the residence time of each state, defined as the fraction of time steps a given trajectory sees that state. An average of the residence time over 100 trajectories, the mean residence time (MRT) is used to evaluate the occurrence of a state for a given parameter set under a given noise type and noise level (Figure 2B). Note that we use the same MRT calculation for deterministic simulations here on (e.g., *σ* = 0). As deterministic simulations reach steady state by the burnin period (shaded in criss-crossed lines in Figure 2B), the MRT measures the relative size of the basin of attraction. For monostable parameters, MRT in deterministic simulations will always be 1, while multistable parameters will see fractional MRT corresponding to the basin size of the specific steady state in the attractor.

**Figure 2:**
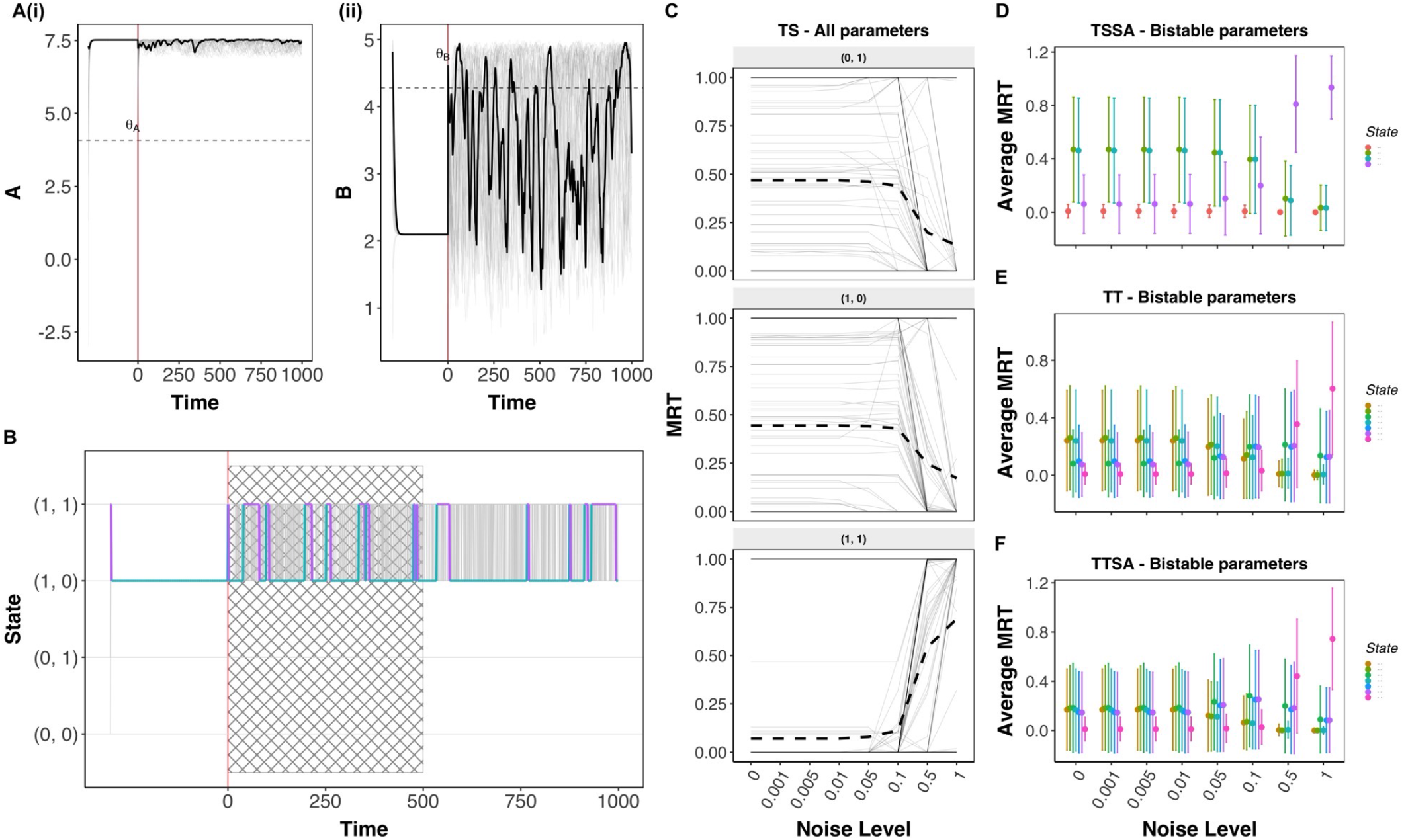
Effect of stationary noise on toggle switch and toggle triad networks. **(A)** Representative single-parameter trajectory for the TS network: (i) node A and (ii) node B expression over time (log_2_), with the deterministic (noise-free) segment at negative time and the stochastic segment at positive time. Gray lines show all sampled initial conditions; the black line highlights one representative trajectory (the one whose switch count is closest to the population median); dashed horizontal line marks each node’s discretization threshold *θ*. **(B)** Discrete-state trace of the same parameter, colored by state for the highlighted trajectory (gray for all other initial conditions). Crosshatched region marks the discarded transient-dominated window of the stochastic segment. **(C)** MRT of each state across noise levels. Each panel corresponds to the state indicated by its label. Grey lines show a random sample of 100 parameter-set trajectories (out of the full population, shown for legibility); the black dashed line shows the mean across the full parameter population, not just the sampled 100. Darker or apparently solid grey segments result from the overlap of multiple trajectories and do not denote a separate category. **(D–F)** MRT vs. noise level for TSSA (D), TT (E), and TTSA (F) bistable parameter sets, colored by state (top 4 states shown for TSSA, top 7 for TT/TTSA), mean *±* SD across parameter sets.

As expected, the dominant states for a toggle switch remain as (0, 1) and (1, 0) for low to moderate *σ* values under stationary noise. However, as the *σ* increases to 0.5 and above (i.e., the noise scales as 50% of the allowed range of *λ*), we see a drop in the MRT of the (0,1) and (1, 0) states and a corresponding increase in the MRT of the (1, 1) state - which represents a co-expression of both nodes that normally inhibit each other (Figure 2C). Note that a similar increase is not seen in the MRT of the (0, 0) state (Figure S2A), an asymmetric outcome despite the symmetry of the stationary noise. We further studied TSSA, TT and TTSA under stationary noise and found them behaving similarly. TSSA, in absence of noise enables bistability in a larger fraction of parameter space than TS (≈20% in TS vs *>* 50% in TSSA). TSSA also enables ≈14% of the state space to be tristable, with access to either (0, 0) or (1, 1) state in addition to the dominant (0, 1) and (1, 0) states. However, the basin size of (0, 0) state, is negligibly small, making (0, 1) − (1, 0) − (1, 1) the only real tristable configuration (Figure S2B - tristable parameters at *σ* = 0). Similar to TS, TSSA maintains the characteristics of its phenotypic landscape under stationary noise - dominant (0, 1) and (1, 0) states in monostable and bistable parameter sets, and dominant (1, 1) state in tristable parameter sets for realistic values of noise, i.e., *σ* ≤ 0.1 (Figure 2D). For higher *σ*, we see a dominance of the (1, 1) state.

Toggle triad (TT) and toggle triad with self-activation (TTSA) dominantly enable two classes of steady states - single-high states ((0, 0, 1); (0, 1, 0) and (1, 0, 0) and double-high states ((0, 1, 1); (1, 0, 1)*and*(1, 1, 0)). Previous studies have shown that the toggle triad motif is more susceptible to various perturbations due to the inherent frustration in the network [36, 22]. Here, we see that the MRT distribution of TT starts to change for *σ >* 0.01, showing a higher sensitivity to noise than TS and TSSA motifs (Figure 2E, F, Figure S2C-D). At 0.01 ≤ *σ* ≤ 0.1, the MRT of single-high states starts to go down and that of double-high states starts to come up, still qualitatively retaining the properties of the phenotypic landscape. However, *σ* ≥ 0.5 leads to an increase in the MRT of the all-high state ((1, 1, 1)) in both TT and TTSA, similar to that of TS and TSSA, with no emergence of the all-low state ((0, 0, 0)). Overall, we find that the phenotypic landscape of both binary and tertiary cell-fate GRNs remains robust to stationary noise of low to moderate levels of *σ*. The response to larger noise can be better understood based on our analysis of additive noise given in the next section.

### 2.3 Additive noise reshapes the phenotypic landscape toward high-expression states, modulated by reachability and threshold margin

Unlike stationary noise, additive noise starts to change the phenotypic landscape even at low *σ* values. For the toggle switch across a majority of the parameters, we see a decrease in the MRT of single-high states ((0, 1) and (1, 0)) and an increase in the hybrid all-high ((1, 1)) state MRT (Figure 3A). The mean MRT of all-high state crosses 0.5 for *σ >* 0.001 (0.1% of the *λ* range) and stabilizes at 0.75 for *σ >* 0.1. Given the biological relevance of hybrid states showing co-expression of all nodes in developmental and pathological contexts, this shift in the phenotypic landscape of cell-fate decision motifs is a very significant finding.

**Figure 3:**
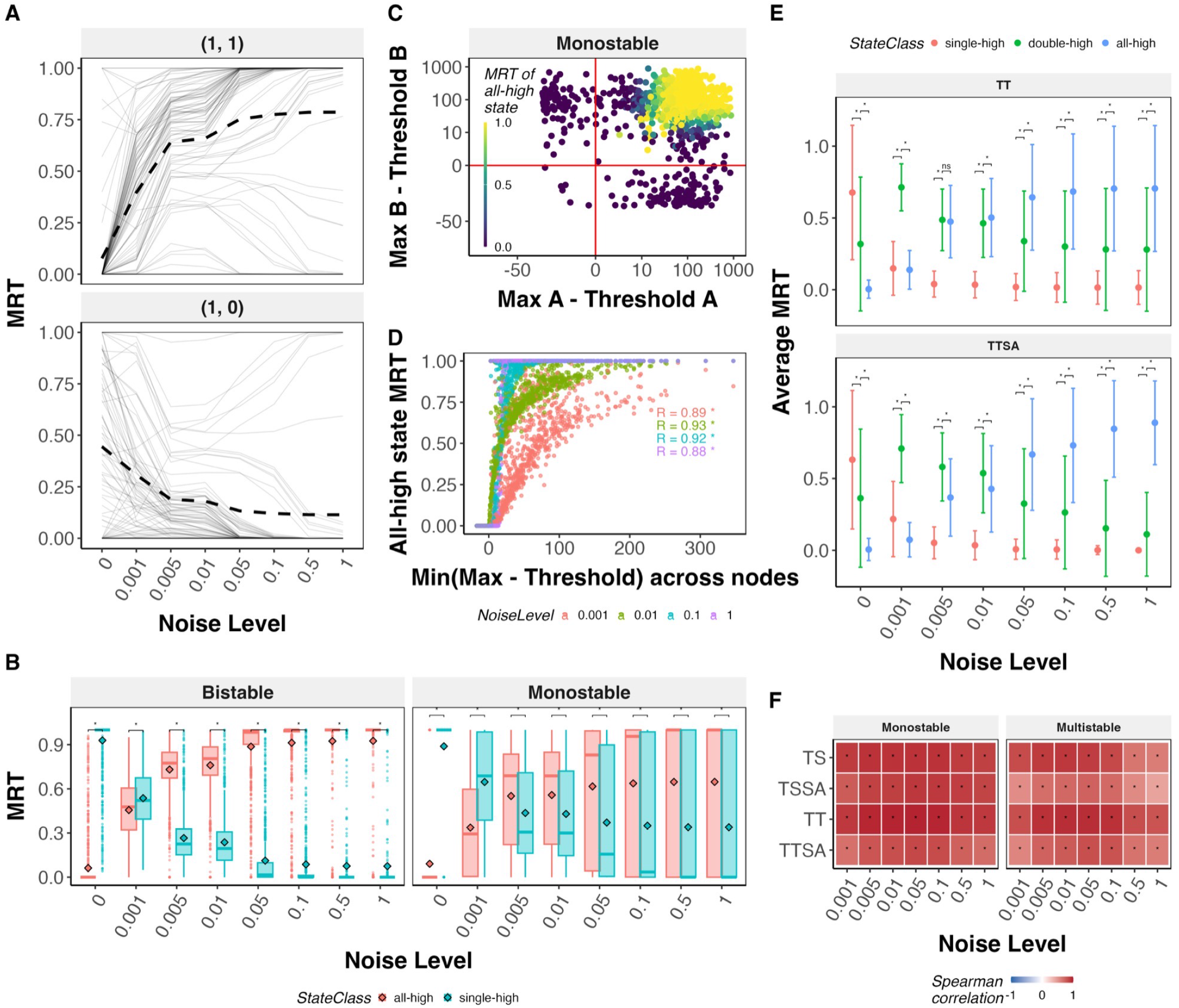
Effect of Additive noise on the phenotypic landscape of cell-fate decision networks. **(A)** MRT of (1, 1) and (1, 0) states vs. noise level for the TS network, a random sample of 100 parameter-set trajectories (gray) with the mean across the full parameter population (dashed). **(B)** MRT boxplots for the TS network, monostable vs. bistable parameter sets, all-high vs. single-high states, with paired Wilcoxon significance brackets; “*” *p* − *value <* 0.05, “ns” not significant. **(C)** Reachability scatter for TS monostable parameter sets at *σ* = 0.1: each node’s (max expression − threshold) margin on the x/y axes, colored by MRT of the (1,1) state. **(D)** Spearman correlation between the minimum reachability margin across nodes and MRT of the (1,1) state, across four noise levels (0.001, 0.01, 0.1, 1), TS monostable parameter sets. **(E)** MRT vs. noise level for TT and TTSA, single-high → double-high → all-high state classes shown as Mean *±* SD across all parameters, with paired Wilcoxon significance brackets between adjacent classes; “*” *p* − *value <* 0.05, “ns” not significant. **(F)** Heatmap of the Spearman correlation between minimum reachability margin and MRT of the all-high state, across networks (TS, TSSA, TT, TTSA), noise levels, and stability class (monostable vs. multistable); “*” *p* − *value <* 0.05, “ns” not significant.

Interestingly, we note that while most parameters gravitate towards a high MRT for the all-high state for *σ* ≥ 0.01, a smaller fraction of the parameters also show a significantly lower MRT of all-high state (Figure 3A). When we separately studied the parameter sets based on their multistability characteristics under deterministic dynamics, we found that monostable states always had a lower MRT for all-high states under additive noise as compared to that of multistable parameter sets (Figure 3B). Given how readily bistable parameters converge to the all-high state, we hypothesized that the reduced MRT for monostable states could possibly arise because the all-high state is not accessible to some monostable parameter sets. The upper limit of node expression in the ODE models we simulated is the ratio of the corresponding production and degradation rates. As we use a threshold derived from the ensemble steady state expression levels, it is possible that some parameter sets are sampled such that the upper limit of expression for one of the nodes defined by the production and degradation rates is lower than the threshold. Since the bistable parameters consist of two single-high states, bistable parameter sets never have this shortcoming.

Thus, we wanted to understand whether it really is the case that some of the sampled monostable parameters reflect systems that cannot access the all-high state and in turn, if the access to all-high state is sufficient for the additive noise to push the system towards all-high state. To do this, we plotted the sampled parameter sets on the axes of “reachability” defined by the difference between the upper limit of a node’s expression level and the threshold used to discretize that node. Reachability was not satisfied for all sampled parameters: for the TS network, 89.0% of parameter sets had a production/degradation-derived expression ceiling exceeding threshold for both nodes, 10.8% reached threshold for only one of the two nodes, and 0.3% reached threshold for neither; among monostable parameter sets specifically, those that are not reachable for both nodes correspondingly show an MRT of exactly 0 for the all-high state, confirming that unreachable parameter sets are the origin of the low-MRT tail seen in the monostable population (Figure 3C, S3A). Among the reachable parameter sets, however, we see that the MRT of the all-high state increases with increasing distance between the maximum allowed value of the nodes and the corresponding thresholds. Across *σ* values, the MRT of the all-high state correlated strongly positively with the smallest of the two difference values (Figure 3D).

TSSA showed a similar increase in hybrid (all-high) state MRT under additive noise. At low to moderate *σ*, TS and TSSA showed very similar MRT distributions. At high *σ* values, TSSA monostable parameter sets showed a higher MRT of all-high states compared to TS (Figure S3B, i). This increase can be attributed to the inherent affinity TSSA has towards all-high state as discussed earlier. For TT and TTSA, we observed a transition of states from single-high to double-high to all-high, suggesting that while additive noise does push the system towards the all-high state, the transformation can become more gradual as the network size increases (Figure 3E, S3B, ii, iii). TT and TTSA have similar MRT distributions for low to moderate values of *σ*, with similar average MRT of all-high and double-high states. But at high values of *σ*, TTSA showed a higher MRT of the all-high state, with the double-high state average MRT approaching 0. TT on the other hand maintains a non-zero average MRT of double-high states even at higher *σ* values (average MRT across all parameter sets - all-high 0.7 vs double-high 0.3 for TT vs all-high 0.9 and double-high 0.15 for TTSA). For all networks, we found a strong positive correlation between the difference quantity (max node expression minus threshold expression of the node) and the MRT of all-high states in monostable parameter sets. Multistable parameter sets showed relatively weaker correlations (Figure 3F). Overall, our analysis clearly shows that additive noise increases the MRT of hybrid, all-high states in cell-fate decision networks.

### 2.4 Increased MRT of all-high states is caused by both the *λ* distribution and stochasticity

Following the strong correlation we observed between parametrically determined upper bounds of node expression and the MRT of all-high state, we hypothesized that the drastic increase in the MRT of all-high states in comparison to deterministic simulations might also be explained by variation in the parameter space. Additive noise models a random walk in *λ*, bounded between 0.01 and 1 for inhibitory links and 1 and 100 for activating links. Any step crossing the boundary is absorbed at the boundary. For example, if *λ*(*t*) + *η*(*t*) *<* 0.01 then *λ*(*t* + *dt*) = 0.01. Due to the nature of random walk, the variance of *λ* increases over time, unlike the stationary noise case. Furthermore, the absorbing boundaries increase the chances of *λ* being at either of the borders than in the interior. The probability density of *λ* at the borders increases with increasing *σ*, as at larger *σ* values, there is a higher chance of taking larger *λ* steps and correspondingly higher chances of hitting the boundaries (Figure 4A). Same boundary conditions are applied to other noise forms as well, and we see such boundary behavior in stationary noise at high *σ*, where *λ* localizes at either of the boundaries. While *λ* closer to zero leads to a very strong inhibition by drastically reducing the net production rate, *λ* closer to 1 leads to very weak inhibition and lets the production rate reach the highest values. Despite this symmetry in the emergent *λ* distribution, the shift in the landscape is asymmetric towards all-high states. One potential reason could be the way the *λ* parameter is sampled in RACIPE for inhibitory interactions. The fold-change parameter is first sampled between 1 and 100 and then the inverse of that number is used for inhibitory links. Thus the distribution of *λ* for inhibitory links is tailed heavily towards lower values of *λ* with a mean of ≈ 0.05. As the stationary distribution of *λ* in additive noise has a mean of 0.5, as a simulation progresses the inhibitory links experience a significant increase in the average *λ*, which in turn leads to weakening of the inhibition.

**Figure 4:**
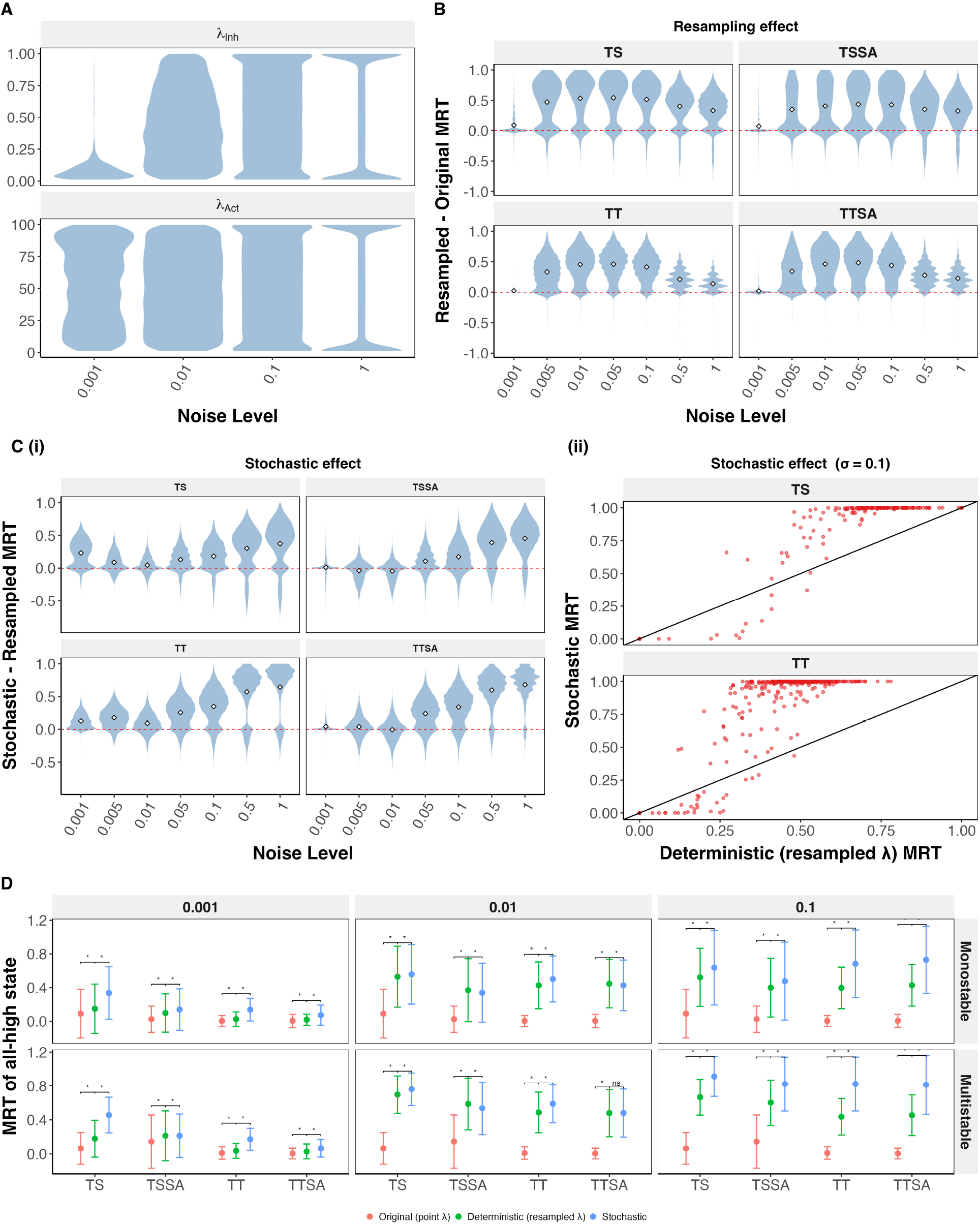
The additive noise effect combines a *λ*-distribution shift with an additional stochastic contribution. **(A)** Distribution of *λ* (Hill-function threshold) values across the parameter ensemble, for (i) inhibitory and (ii) activation-type edges, at four noise levels (Additive noise). **(B)** Distribution of the resampling effect - difference in deterministic MRT after *λ*-resampling and the original point-*λ* deterministic MRT for the all-high state, shown as violin plots across all noise levels, for TS, TSSA, TT, and TTSA. **(C)** (i) Same as B, but for Stochastic effect (stochastic MRT minus resampled-*λ* deterministic MRT). (ii) Scatter plot of resampled-*λ* deterministic vs. stochastic MRT of the all-high state at *σ* = 0.1, for TS and TT, illustrating the underlying sigmoidal relationship the violin’s signed difference alone does not show. Each point is a parameter set. **(D)** All-high-state MRT (mean *±* SD over all monostable and multistable parameter sets) for TS, TSSA, TT, and TTSA, comparing the original point-*λ* deterministic run, the *λ*-resampled deterministic run, and the full stochastic simulation; faceted by stability class of the parameter sets (rows) and noise level (columns: 0.001, 0.01, 0.1), with paired Wilcoxon significance brackets; “*” *p* − *value <* 0.05, “ns” not significant.

Given the above, we wanted to know if this shift in *λ* distribution is enough to completely explain the increase in the MRT of all-high states. For each parameter set used for stochastic simulations, we sampled *λ* from the additive noise stationary distribution shown in (Figure 4A). For each such *λ* combination, we simulated the networks deterministically (*σ* = 0) at multiple randomly sampled initial conditions and calculate the mean frequency of each discrete state across initial conditions, which is further averaged over different *λ* values. While the original RACIPE parameters for TS and TT have a very small fraction of parameter sets with non-zero frequency of all-high states, the majority of resampled simulations led to a non-zero frequency of all-high states, reaching up to 0.9 in TS and up to 0.75 in TT (Figure 4B, S4A). For TSSA and TTSA, which inherently stabilize the all-high states due to the self-activation links, resampled simulations still led to an increase in the frequency of all-high states for a majority of the parameter sets (Figure 4B, S4A). Thus, the change in the *λ* distribution does significantly contribute to increasing the MRT of all-high state. This might be relevant if a GRN is subjected to the onset of regulatory noise due to its interaction with some additional dynamical factor

Interestingly, however, when we compared the MRT of all-high states in stochastic simulations and the resampled deterministic simulations, we found that the stochastic MRT was pushed even higher across a large fraction of parameters for all four networks (Figure 4C, i, S4B). Furthermore, a sigmoidal relationship emerges between the stochastic MRT and the deterministic resampled frequency of all-high state (Figure 4C, ii, S4C). Below a threshold value of all-high state frequency under deterministic simulations with resampled *λ*, stochastic MRT is lower than or equal to deterministic frequency, and vice-versa (Figure 4C). The specific threshold MRT is network and *σ* dependent (Figure 4C, ii, S4C). Across all four networks and *σ* values, the MRT of all-high states increased significantly from original to resampled deterministic to stochastic simulations (Figure 4D, S5). We further tested the effect of stochasticity by reducing the frequency of update of *λ*, i.e., the *dt* value in Equation 3, from 0.01 to 1.0 and 10.0 (Figure S6). As the frequency of update decreased, the system had more time to relax towards a steady state, thereby losing the effects of stochasticity. Correspondingly, we see a reduction in the MRT of all-high state and increase in the MRT of the native states of these networks at lower *σ* values. At higher *σ* values, the noise compensates for the relaxation, pushing the systems back to hybrid states. These results indicate that, while the distribution of *λ* does shift the landscape towards the all-high hybrid state, the stochastic simulations serve to further enhance this shift.

To isolate the mechanism by which regulatory (*λ*) noise redistributes occupancy toward high-expression states, we analyzed the toggle switch in the Boolean limit of node expression and *λ* distribution, where the exact stationary distribution can be computed in closed form for both the deterministic (fixed-*λ*) and stochastic (fluctuating-*λ*) cases (see Supplementary section S2). We noticed a crucial asymmetry in the stochastic simulations that is absent in deterministic simulations. For an inhibitory edge, the shifted Hill function entering the production term is

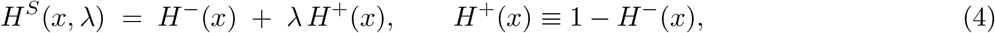

Under the extreme case of *σ* = 1, where the *λ* walk is confined to the two boundaries of its range, *λ* ∈ {*λ*_min_, *λ*_max_} with *λ*_min_ ≈ 0.01 and *λ*_max_ = 1. Evaluating Eq. (4) at these two limits exposes a structural asymmetry:

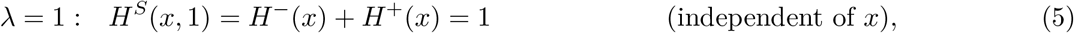

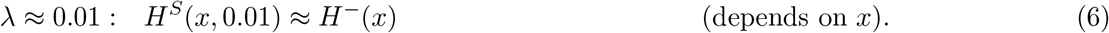

The *λ* = 1 boundary abolishes regulation *unconditionally*: the target node is produced at its full rate regardless of the regulator’s expression. The *λ* ≈ 0 boundary imposes repression only *conditionally*: it lowers the target’s production rate only when the regulator *x* is high, and has essentially no effect when the regulator is low (*H*^−^(*x*) ≈ 1). An excursion toward de-repression (*λ* = 1) therefore always “fires,” whereas an excursion toward repression fires only when the partner node happens to be expressed. This unconditional-up / conditional-down asymmetry is the microscopic origin of the enhancement. We see this clearly in the Boolean approximation of node expression (i.e., each node can only take two expression levels - either 1 or 0). In the extreme *σ* condition, resampled deterministic simulations can only converge to the all-high state if they start from the all-high state (a frequency of 0.25). However, in stochastic simulations, asymmetry in the response of shifted Hill function ensures that a strongly repressed state is maintained only when the source node is on, or when both are off. This asymmetry leads to a higher probability (equivalent to MRT) of all-high state in the stationary distribution of the stochastic process. Even at lower values of *σ*, this asymmetry holds true by making the de-repressed state more easily accessible to the nodes of the network than the repressed states. Therefore, we conclude that the drift in the phenotypic landscape caused towards the all-high state caused by additive noise is a combined outcome of the shift in the parameter space and the asymmetry exploited by stochastic simulations.

### 2.5 Multiplicative noise drives the phenotypic landscape toward single-high states - an opposite effect as that of additive noise

We then analyzed the effect of multiplicative noise in *λ* on the phenotypic landscape. Briefly, multiplicative noise models a random walk in *λ* with a state-dependent step size. For inhibitory interactions, the noise amplitude is smaller near *λ* ≈ 0 and larger near *λ* ≈ 1. This asymmetry makes *λ* approach 0 over time, unlike additive noise where *λ* visits both the boundaries and the intermediate values at intermediate *σ* (Figure 1D). While this concentration of *λ* near 0 may seem like a deterministic behavior, we will see below that stochasticity has a strong impact on the outcome of multiplicative noise, even more so in certain cases than additive noise.

Intuitively, multiplicative noise enhances repression in the edges of toggle switch network, thereby driving the system strongly towards single-high states (Figure S7A, i). This does not affect the phenotypic landscape of TS, as single-high states were dominant to begin with (Figure 1C). TSSA also showed a similar gravitation towards single-high states despite TSSA also stabilizing the all-high state in absence of noise (Figure S7A, ii). The enhancement of single-high state by multiplicative noise is prominently seen in the case of TT and TTSA. Both these networks have a significantly high frequency of double-high state along with single-high state in the absence of noise (Figure 1C). In the presence of multiplicative noise, both these networks converge to single-high states, achieving a high MRT by *σ* = 0.01 (Figure S7A, iii, 5A). Thus, consistently across the four networks studied so far, we found that multiplicative noise promotes the frequency of single-high states (Figure 5B). At very high *σ* values, TTSA shows a decrease in the frequency of single-high states with a corresponding increase in double-high state MRT. At high *σ, λ* starts to approach higher values purely based on the fluctuations (Figure S7B). These excursions weaken enough inhibitory interactions to increase the occupancy of double-high states, but they are not sufficiently widespread to permit all nodes to be highly expressed simultaneously. We therefore observe an increase in double-high-state MRT without the emergence of the all-high state.

**Figure 5:**
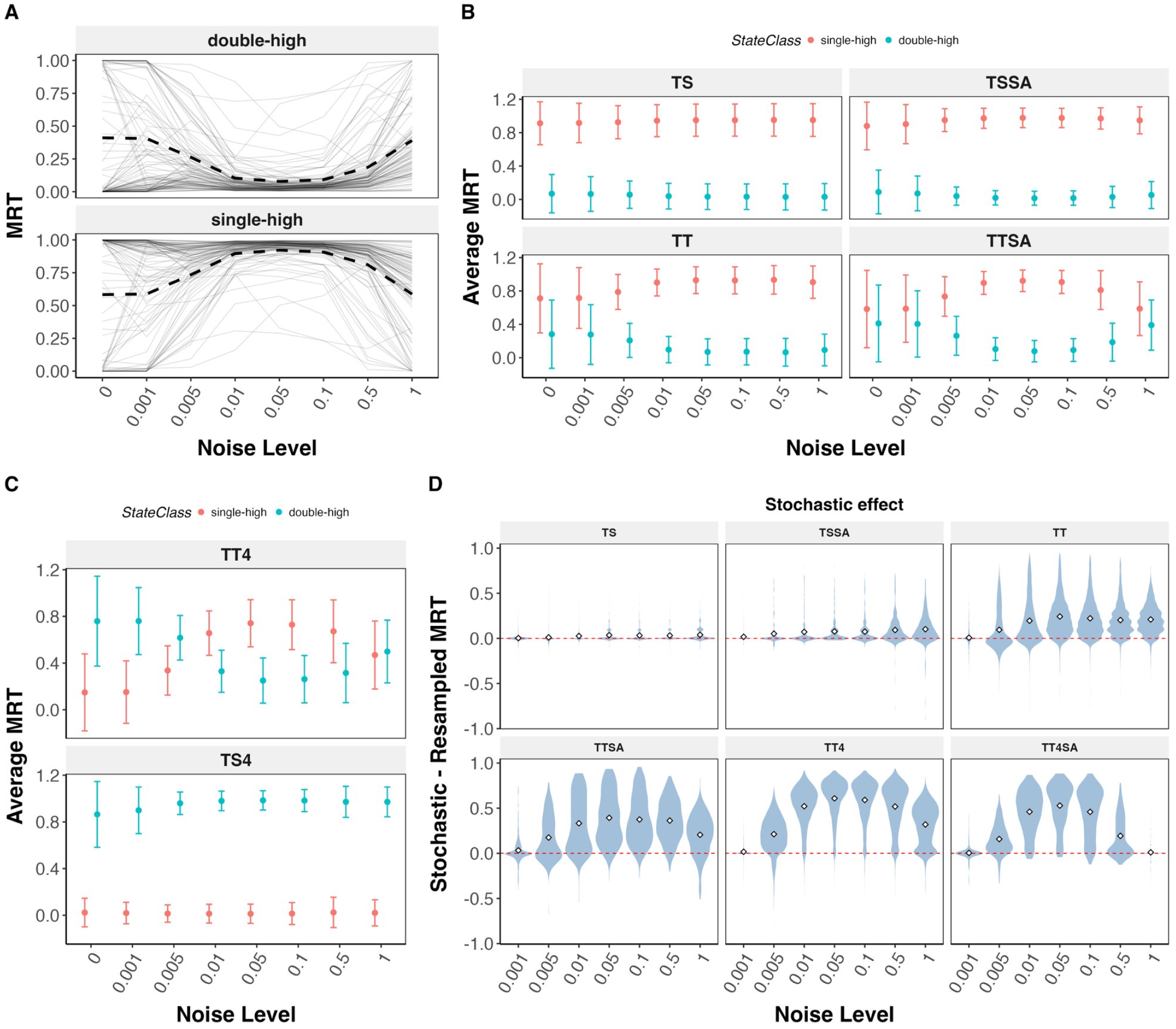
Effect of multiplicative noise on the phenotypic landscape of cell-fate decision networks. **(A)** MRT vs. noise level for the TTSA network, single-high and double-high states; grey lines show a random sample of 100 parameter-set trajectories, with the mean across the full parameter population shown separately. **(B)** Average MRT of single-high vs. double-high states vs. noise level, for TS, TSSA, TT, and TTSA. **(C)** Same, for the 4-node networks TT4 and TS4. **(D)** Stochastic effect (as in Fig. 4C(i)) for the single-high state, across noise levels, for TS, TSSA, TT, TTSA, TT4, and TT4SA.

To further validate this result, we simulated three new networks that are seen in cell-fate decision systems. The first network is an extension of TT network in the context of T cell differentiation, including the regulation of T-reg cells through the transcription factor Foxp3. The resultant four-node network is referred to as toggle tetrahedron (TT4), and is composed of four nodes mutually inhibiting each other. Previous studies have explored this network as well as its self-activation counterpart (TT4SA), our second 4-node network, as candidates for four-fate differentiation [40] (Figure S7C, i, ii). However, this and other candidate networks for multi-fate systems have so far failed to support the emergence of single-high states representative of fully differentiated cell-fates. TT4 and TT4SA, specifically support only double-high steady states (Figure S7D). Given the properties of multiplicative noise seen so far, we wanted to study if the double-high states of TT4 and TT4SA could also be driven towards single-high states. The third network is a combination of two mutually inhibiting toggle switches that we refer to as TS4, a representation of coupled cell-fate decision systems often seen in biology [41, 42]. TS4 also supports double-high states (Figure S7D). However, unlike TT4 where all six possible double-high states have equal frequency, TS4 only supports a limited number of double-high states that represent the coexistence of the two binary cell-fate systems. Unlike sequential differentiation, coupled systems are expected to have co-existing phenotypes governed by the network structure. Thus, we asked whether multiplicative noise pushes these coexisting double-high states into single-high states, irrespective of the topology. Under additive noise, all three networks lead to a consistent increase in all-high state, despite the differences in the network structure (Figure S8A). Under multiplicative noise, TT4 saw an increase in the single-high state MRT with *σ*, followed by a decrease brought on due to the boundary sampling of *λ* (Figure 5C, S8B). However, TT4SA maintained a double-high dominant landscape (Figure S8B). TT4SA did show a decrease in double-high state MRT as multiplicative noise increased, with a corresponding increase in the MRT of single-high states. However, this change never put single-high states in a dominant position. Furthermore, TS4 also maintained a double-high dominant landscape, with no change in the MRT with *σ*, preserving the behavior of internal toggle switches. Thus, unlike additive noise, the effect of multiplicative noise on the phenotypic landscape is strongly dependent on the network topology.

Given the potentially deterministic nature of multiplicative noise, we tried to dissociate the effects of the altered *λ* distribution from that of stochasticity in these simulations. Thus, we generated resampled deterministic simulations similar to that of Figure 4C. For TS and TSSA, the difference between stochastic and resampled deterministic MRT was very low, as expected (Figure 5D). However, TT, TTSA and TT4 showed large differences even at lower values of *σ* (< 0.05, Figure 5D). In comparison, additive noise simulations required higher *σ* ≥ (0.1) to achieve this level of difference (Figure 4C). TT4SA and TS4 also showed minimal difference, in accordance with their MRT trends. These differences were mirrored by corresponding decreases in the MRTs of single-high states (Figure S8D, E). Interestingly, however, the MRTs of original deterministic simulations and resampled simulations were nearly identical (Figure S8F, G), indicating that this “fixation” towards terminal differentiation seen in multiplicative noise is entirely due to the stochasticity, unlike the additive noise where the *λ* distributions also contributed to the increase in MRT of all-high states. We further tested this behavior by simulating these networks at larger dt values (Figure S9). Unlike additive noise, increasing dt did not reduce the excess of stochastic MRT over the resampled-deterministic MRT for multiplicative noise. Since larger dt gives node expression more time to relax toward the steady state set by the current *λ* before the next update, this indicates that multiplicative noise’s dynamical contribution — unlike additive noise’s — is not primarily a consequence of expression lagging behind a fast-moving *λ*. The specific mechanism by which individual stochastic realizations of *λ* produce this persistent excess remains to be characterized, but our results indicate it is distinct from the timescale-separation (non-adiabatic) effect identified for additive noise.

### 2.6 Multistable double-high states resolve toward their nearest single-high fate across network motifs

We next asked which single-high states the hybrid double-high states converge to under multiplicative noise. Is a double-high “progenitor” restricted to one terminal fate, or can it access several? We focused this analysis on TT4, which has a high double-high and low (≈ 0) single-high frequency in deterministic simulations (Figure S7D) and gives the cleanest signal of double-high to single-high transition in the network motifs we studied so far (Figure 5C). To confirm that this terminal commitment reflects genuinely stable fates rather than continued flickering between states, we examined switching-event rates, measured the mean number of switches in discrete state space observed during stochastic simulations at different noise level (Figure 6A, S10). Under additive noise, switching rates rose with noise level before leveling off or mildly declining by *σ* = 0.1, across every network and stability class. Under multiplicative noise, by contrast, switching events remained close to zero throughout this same range, regardless of network or class. A notable trend however is that while additive noise showed a decrease in the switching rate as noise increased, multiplicative noise showed an increase owing to the spread of *λ* distribution discussed earlier.

**Figure 6:**
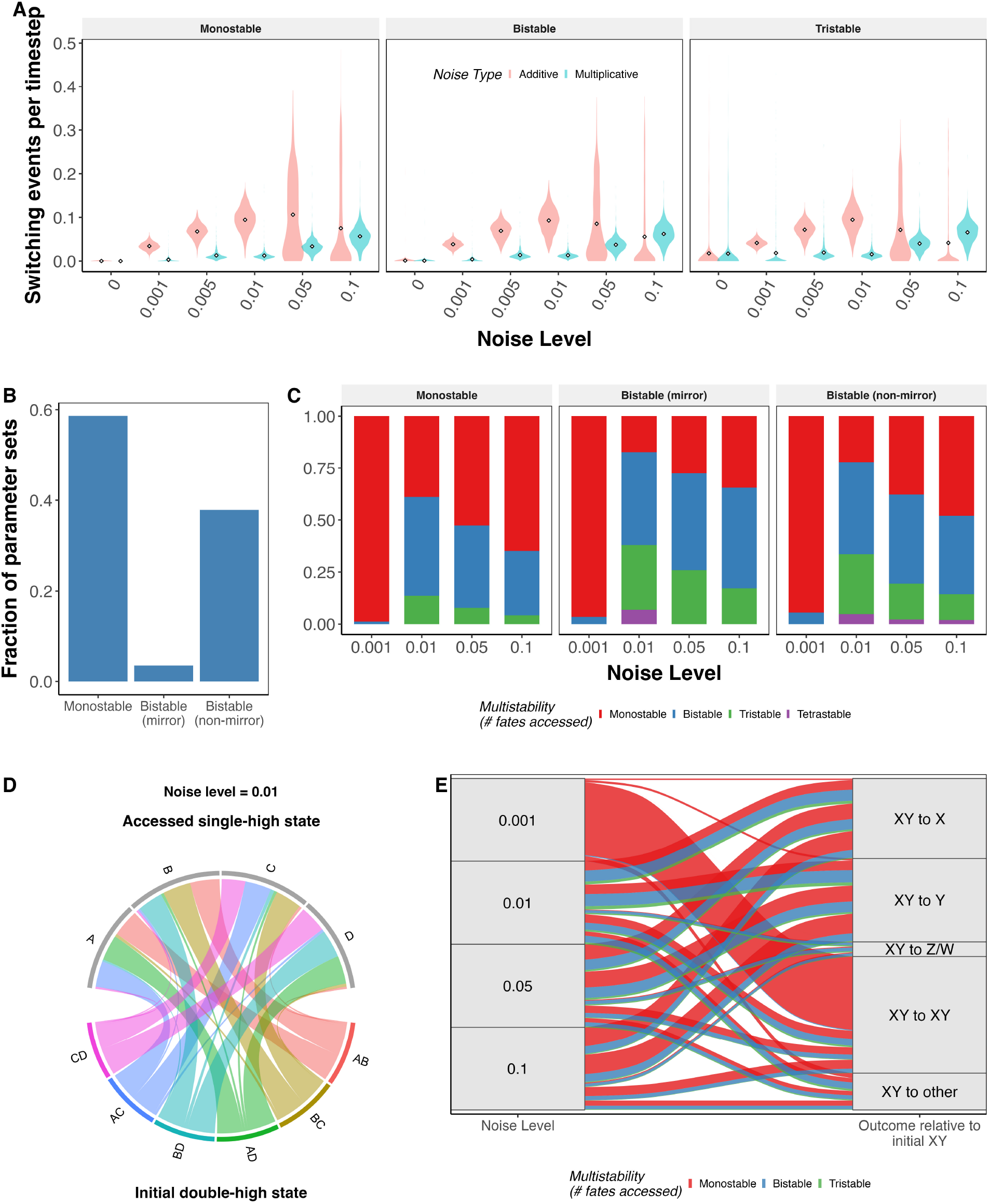
MRT-based multistability and transition analysis for the TT4 network. **(A)** Distribution of switching-event counts by noise level (Additive vs. Multiplicative), after excluding the initial burn-in window. **(B)** Frequency of the three deterministic stability classes among double-high parameter sets. **(C)** For each stability class, the fraction of parameter sets accessing 1 (Monostable), 2 (Bistable), or 3+ (Tristable/Tetrastable) distinct single-high fates under noise. **(D)** Circular chord diagram of double-high → single-high MRT flow among monostable double-high parameter sets, at noise = 0.01. **(E)** Noise-level → outcome alluvial for the pooled monostable double-high population (outcomes labeled relative to the initial double-high state XY, e.g. “XY to X”), colored by the multistability classification from panel C.

We then moved to characterize the nature of transitions from double-high to single-high states in TT4. We classified the parameter sets of TT4 as monostable double-high, bistable mirror double-high and bistable non-mirror double-high. TT4 parameter sets sampled by RACIPE were predominantly monostable and bistable non-mirror, with a small mirror-bistable population (Figure 6B). Intuitively, we expect bistable parameters to have access to more than two single-high states. Given the relative differences in expression patterns (i.e., the closest single-high states for a set of double high states making up the attractor), bistable mirror parameter sets have a higher chance of accessing all four single-high states, potentially representing a true progenitor configuration. Therefore we first quantified the number of single-high states each of these parameter classes accessed under multiplicative noise (Figure 6C). At *σ* = 0.001, all parameter classes only accessed one single high state. As noise level increased, the fraction of parameters accessing multiple single-high states increased. Monostable parameter sets primarily accessed one or two single-high states, while bistable parameter sets accessed two and three single-high states. While the mirror bistable parameter sets did not have access to all four single-high states as we intuitively predicted, a larger fraction of these parameter sets accessed more than one single-high state under multiplicative noise.

We then characterized the nature of the single-high steady states accessed by monostable double-high parameter sets. For ease of reference, we adapt an alphabetical notation to represent states where we use the set of nodes which have a high expression in that state. For example, ′*A*′ represents the single high state (1, 0, 0, 0) while ′*AD*′ represents the double high state (1, 0, 0, 1). The single-high state(s) accessed under multiplicative noise (*σ* = 0.1) were majorly the “nearest” ones – reached by silencing one of the two active nodes rather than switching to an unrelated fate (Figure 6D, for example, CD monostable parameter sets predominantly access C and D states, with a very few accessing A and B state). We found this trend consistently across all networks studied (Figure S11). However, these transitions varied significantly with noise level (Figure 6E). At lower noise levels (*σ* = 0.001), most parameter sets retained their original double-high attractor (XY to XY, where X and Y are any two of the four nodes in TT4). As noise level increased, parameter sets split between three routes - going to the closest single-high state (XY to X and XY to Y) or remaining in the original double high state, with the choice of which single high state to access attributed, perhaps, to the specific parameter sets. A smaller fraction of parameters also transition to other double-high states. Parameter sets accessing more than one single-high state also showed the proximity bias. Correspondingly, access to non-proximal single-high state remained low across all parameter sets and noise levels. This picture is consistent with multiplicative noise leading to strengthened inhibition. The relative strengths of inhibitions in the network are primary determinants of which double high state is accessed, with the inhibition on low expression nodes stronger than the inhibition on high expression nodes [40]. Intuition suggestst that, a further strengthening of inhibitions uniformly across all edges by multiplicative noise is bound to induce a single high state where one of the two weaker inhibitory links gets strengthened, with the choice of which node gets inhibited depending on which of inhibitions incoming to the two high expressing nodes was already stronger. Switching to a state different from the three closest choices (XY to XY, X or Y) would require a larger fluctuation in *λ* allowing some of the inhibitions to weaken instead, which can only happen at higher noise levels where tail in *λ* distributions towards weaker inhibitions grows (Figure S7B). Consistently, we find that the non-proximal transitions increase with increasing noise levels (Figure 6E). We confirmed this same pattern – nearest-fate commitment, rare late-stage switching, no single-to-single transitions – in TT, TTSA, and TT4SA, indicating that multiplicative noise’s ability to push cell-fate systems from hybrid, progenitor-like states toward terminally differentiated ones is a general property of these topologies rather than one specific to TT4.

We extended the noise-level → fate alluvial analysis of Figure 6E to the bistable double-high classes (Figure S12). Mirror bistability is structurally impossible in the 3-node TT and TTSA networks (any two of three nodes must share at least one active node), so only their non-mirror class is shown (Figure S12A, B); TT4 has enough parameter sets in both classes to show them separately (Figure S12C), while TT4SA’s mirror-bistable population is too sparse (≈ 13 parameter sets) to interpret alone and is pooled with its non-mirror population (Figure S12D). Across all of these, the “Self” (return-to-double-high) outcome remains a substantial fraction of the bistable-class population even at *σ* = 0.1, more so than for the monostable class in Figure 6E – consistent with our choice to base the quantitative claims above on the monostable subclass, where the fate-resolution signal is cleanest.

### 2.7 Effects of regulatory noise on the phenotypic landscape scale to larger biological networks

While the toggle switch and TSSA are well-studied representatives of binary cell-fate decision systems, the regulatory circuits underlying real cell-fate transitions are rarely small or as isolated. Phenotypes emerge from coordinated expression across tens to hundreds of genes, many of which function as transcription factors that individually regulate transitions between phenotypes. It is therefore important to ask whether the noise-driven dynamics observed in small motifs generalize to larger, more complex GRNs. To address this, we asked whether biological GRNs governing two well-studied cell-fate decisions – epithelial-mesenchymal plasticity [43] and Gonadal cell-fate decision [44] – behave similarly to our studied examples.

Large-scale GRNs underlying binary cell-fate decisions often share a common coarse-grained structure: nodes can be partitioned into two “teams”, with intra-team interactions predominantly activating and inter-team interactions predominantly inhibitory (Figure S13). At a coarse-grained level, this structure is analogous to a TSSA, with each super-node representing the collective of genes defining one phenotype. The biological GRNs adhere to this structure at a coarse-grained level as well. One specific EMT model, a 23-node network underlying the epithelial-mesenchymal transition (EMT) [31], has two teams of nodes corresponding to the epithelial and mesenchymal phenotypes respectively. In fact, we previously showed that many large-scale EMT GRNs constructed to date could be coarse-grained into two mutually inhibiting teams, and that the presence of these teams gives rise to a robust, low-dimensional phenotypic landscape despite the size of these networks [43, 45, 46]. We also showed that these teamed networks are robust to noise in the network state, and extensive mutations disrupting the network of positive feedback present in these GRNs are required to stabilize what is otherwise a weakly stable hybrid phenotype [47, 48]. The 19-node gonadal network, governing cell-fate decisions between Sertoli and Granulosa fates that are upstream of male and female development respectively [44], has two teams of nodes corresponding to Sertoli and Granulosa phenotypes, and carries the advantages of team structure across simulation formalisms [49, 37].

Given the size of these networks, exploration of regulatory noise using similar ODE based models as with small-scale networks earlier would be computationally infeasible. Hence, we utilized a threshold-based Boolean formulation to analyze these networks instead [50]. In this formalism, each node is allowed to take either -1 (low) or +1 (high) expression levels. The edges of the network are given a weight of +1 (activation) or -1 (inhibition). The state of the network is updated asynchronously, based on an Ising-model-inspired, additive rule (see Methods). The binary state vectors (*S* ∈ { 0, 1} ^*N*^) were converted into team scores — the fraction of nodes in each team that are active — and a phenotypic score defined as the difference between the two team scores. Values near *±*1 indicate terminal phenotypes (epithelial/mesenchymal or Sertoli/Granulosa), while values near 0 indicate hybrid states. Under deterministic conditions, the phenotypic landscape is largely concentrated in regions corresponding to terminal states with team scores *±*1.

Under additive noise, both biological networks show behavior consistent with the small network analysis. Trajectories initiated from terminal phenotypes of the unperturbed network rapidly migrate to hybrid phenotypic scores and remain there throughout the simulation, regardless of initial condition (Figure 7A). The transition occurs sharply: even at *σ* = 0.001, terminal state frequencies are markedly reduced (Figure 7B), and by *σ* = 0.005 the phenotypic score distribution has saturated into a stable hybrid-dominated configuration that changes little with further increases in noise (Figure 7C, S14A). Multiplicative and stationary noise produce minimal disruption to the phenotypic landscape, with trajectories retaining high terminal state frequency similar to the unperturbed network across all noise levels tested (Figure 7C, S14B, C).

**Figure 7:**
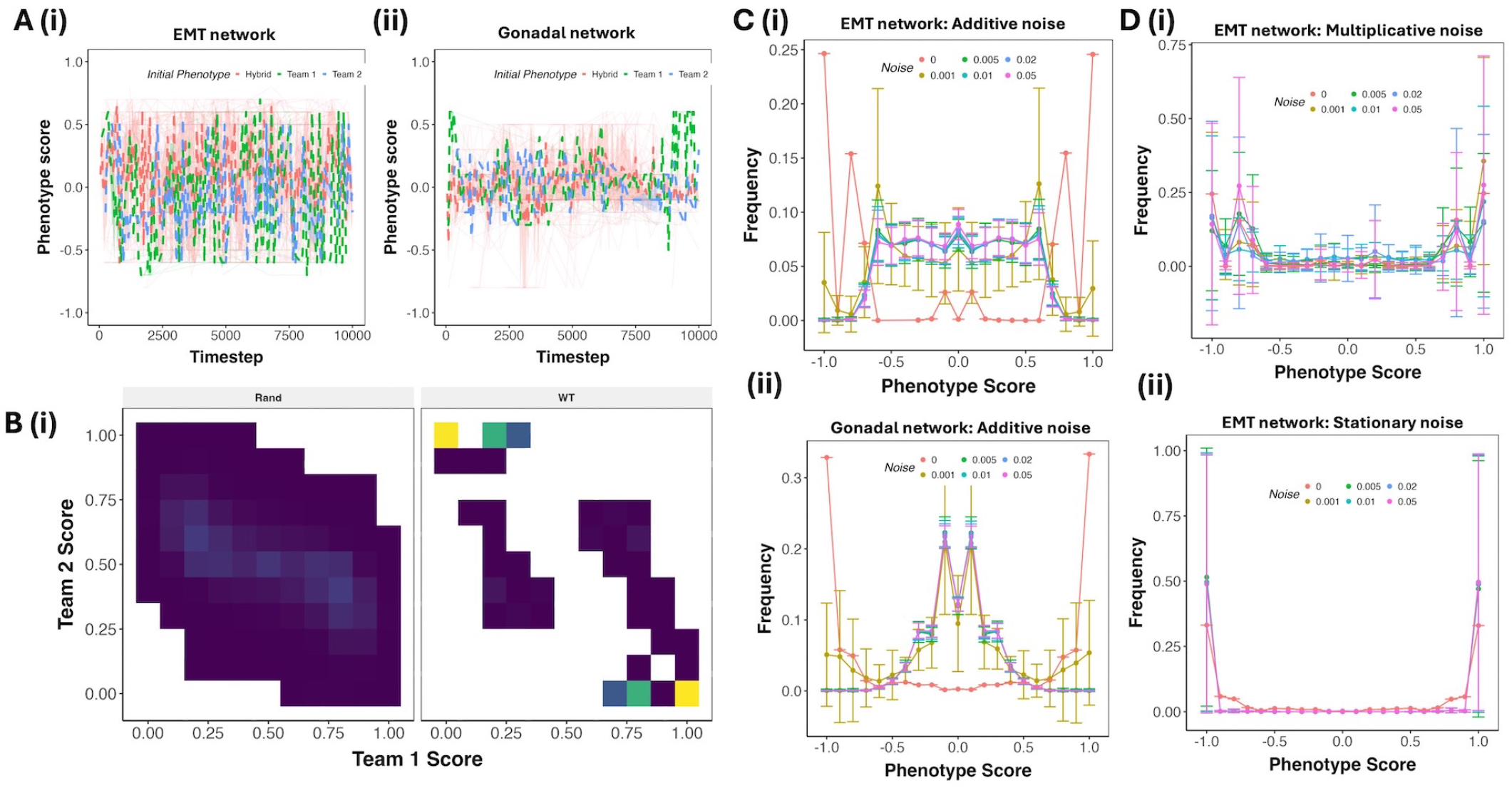
Regulatory noise in large-network Boolean models. A) Representative trajectories of the phenotypic score for EMT network (i) and Gonadal network (ii) at additive noise of 0.05 magnitude. B) Density of team scores across the representative trajectory of the EMT network, with frequency represented on a scale from low (blue) to high (yellow). WT represents the frequency of team score configurations obtained from the steady states of the unperturbed network, while Rand represents that of the simulations corresponding to the trajectories in A. C) Relative frequency (equivalent to MRT) of different phenotypic scores across 1000 simulation trajectories of additive noise over simulation time for (i) EMT network, (ii) Gonadal network. D) Same as C, but for Multiplicative and Stationary noise for EMT network.

The two networks differ, however, in the characteristics of their hybrid states. In the EMT network, the hybrid phenotypic score distribution retains some polarization toward *±*0.5–0.75, reflecting a tendency for one team to remain more active than the other even in the hybrid regime (Figure 7C(i), B). In the Gonadal network, the distribution collapses tightly to near zero, indicating near-equal team expression with no residual polarization (Figure 7C(ii)). Both networks have similar densities and team structures to each other, suggesting that network size and specific wiring topology beyond the team structure are likely contributors to this difference. A full characterization of the structural features that determine the degree of polarization within the hybrid regime remains an open question. Overall, our results indicate that regulatory noise can have a significant impact on the phenotypic landscape, with the direction of the impact determined by the nature and the magnitude of the noise. While additive noise can drive the cell-fate decision system up towards an undifferentiated phenotype, multiplicative noise can drive the system down towards a terminally differentiated state (Figure 8).

**Figure 8:**
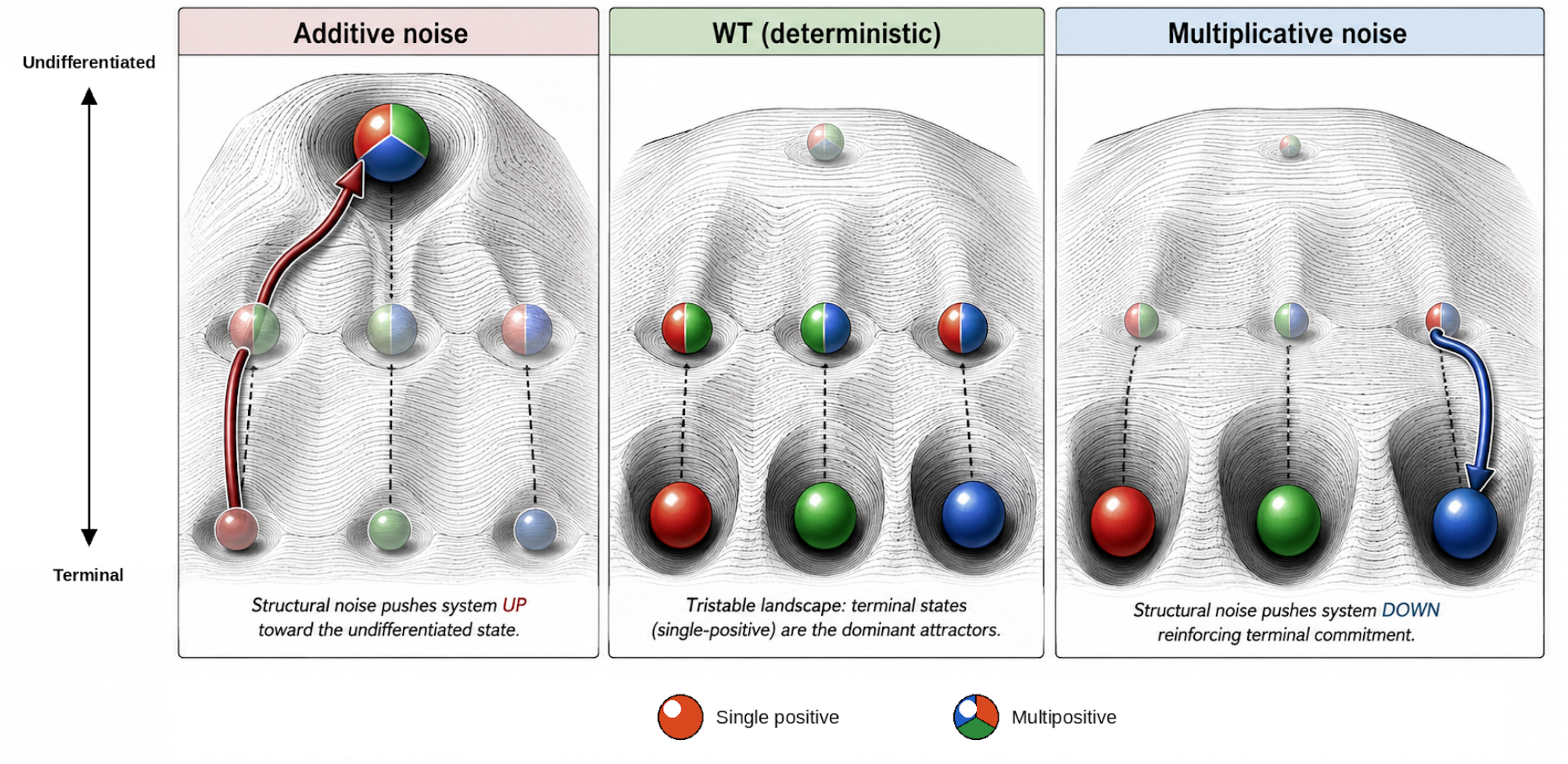
Schematic representation of regulatory-noise-induced remodeling of the phenotypic landscape. The landscapes reimagine Waddington’s phenotypic landscape [51] in the context of the toggle triad. In the deterministic (WT) system (center), hybrid (double-high) and terminally differentiated (single-high) states constitute stable attractors. Additive noise (left) reshapes the landscape by stabilizing an undifferentiated all-high state, whereas multiplicative noise (right) drives the transitions from hybrid states to terminally differentiated states. Colored sectors indicate the nodes expressed in each state.

## 3 Discussion

Historically, regulatory interactions in gene networks models have been remarkably successful at predicting cell-fate outcomes from EMT circuits to Boolean perturbation frameworks. Nonetheless, stochasticity in kinetic parameters is unavoidable. Therefore, here, we introduced regulatory noise into GRN models by allowing the fold-change parameter governing regulatory efficacy of a TF to be a stochastic process. We found that regulatory noise can significantly alter the phenotypic landscape, with qualitatively different outcomes depending upon the *mode* of regulatory noise. Across cell-fate decision motifs, additive noise drove networks toward hybrid co-expression states, multiplicative noise favored single-high terminal states, and stationary noise largely preserved the deterministic landscape at low-to-moderate noise. These effects were reproduced qualitatively in two well-characterized biological GRNs, namely EMT and gonadal cell-fate determination, suggesting that regulatory stochasticity can act not merely as a perturbation around a fixed landscape, but as a determinant of which regions of that landscape are occupied.

Prior theoretical work has shown that noise can alter the effective bistable range of regulatory circuits and generate switch-like population distributions even in deterministically monostable systems [52]. Our results extend this idea to stochasticity in the regulatory interactions themselves. The (1, 1) co-expression state, negligible in the deterministic toggle-switch landscape, becomes dominant under additive noise, whereas multiplicative noise can generate single-high states in networks such as the toggle tetrahedron where these states are not stable deterministically. Thus, similar population-level expression patterns can arise from different combinations of network topology and regulatory-noise regime. A positive correlation between two genes, for example, need not imply mutual activation if stochasticity of mutual inhibition can generate the same pattern. Inferring regulatory wiring from steady-state expression data without accounting for regulatory stochasticity may therefore attribute noise-induced phenotypes to structural features that are not actually present.

The three noise modes studied here can be viewed most usefully not as literal descriptions of individual molecular mechanisms, but as phenomenological limits of how fluctuations in regulatory efficacy evolve in time. This distinction is particularly relevant for intrinsically disordered TFs, for which regulatory function emerges from conformational ensembles spanning a hierarchy of timescales. Rapid conformational fluctuations may be effectively averaged over by the slower transcriptional response, whereas post-translational modifications can stabilize particular conformers for minutes to hours [7, 8]. Still slower processes, including chromatin remodeling and epigenetic feedback, can provide cellular memory across timescales much longer than that of an individual binding event.

Stationary noise represents the regime closest to the conventional deterministic approximation: regulatory parameters fluctuate around a preferred set of values and therefore retain memory of their underlying interaction strength. At low-to-moderate noise, such fluctuations have little effect on phenotype occupancy, consistent with a separation of timescales in which sufficiently rapid or sufficiently constrained molecular fluctuations are averaged into an effective deterministic parameter. Only at high noise does boundary sampling appreciably reshape the landscape.

Additive noise represents the opposite limit. Here, *λ* undergoes a bounded random walk whose long-term distribution loses memory of its initial value. For inhibitory interactions, this progressively increases access to weakly repressive configurations and drives the system toward hybrid co-expression states. A possible molecular analogue is unconstrained sampling of an IDP conformational ensemble or regulatory configurations generated by non-specific binding, in which successive changes in effective regulatory strength are only weakly anchored to the previous interaction state [5, 6]. Importantly, our Δ*t* analysis shows that the resulting phenotype shift depends on the relative timescales of parameter fluctuation and expression relaxation. When *λ* changes rapidly enough that expression cannot fully relax to the instantaneous landscape, additive noise produces an additional hybrid-state enrichment beyond that predicted by the stationary *λ* distribution alone. Thus, the ratio between the correlation time of regulatory fluctuations and the response time of the downstream GRN may itself be an important determinant of cellular plasticity.

Multiplicative noise produces a qualitatively different outcome because the magnitude of each perturbation depends on the current value of *λ*. For inhibitory interactions this creates a self-reinforcing drift toward stronger repression and consequently toward single-high states. PTM-mediated stabilization of particular IDP conformers provides one possible conceptual analogue: phosphorylation and other modifications can stabilize particular conformational substates over timescales substantially longer than rapid conformational exchange [7, 8], allowing past molecular states to influence future regulatory efficacy. We do not interpret multiplicative noise as a direct model of any specific modification process; rather, it captures the qualitative consequences of introducing state-dependent memory into the evolution of regulatory strength. Notably, its excess stochastic effect persists even when parameter updates are slowed, indicating a dynamical mechanism distinct from the non-adiabatic expression lag that contributes to the additive-noise phenotype.

This timescale perspective also clarifies why the stochastic simulations cannot be reduced to parameter resampling. Deterministic simulations sampled from the noise-induced *λ* distributions reproduced only part of the observed phenotype redistribution. Both additive and multiplicative noise produced state occupancies beyond those predicted by their stationary parameter distributions, demonstrating that the temporal ordering and persistence of regulatory states matter in addition to the values sampled. Regulatory noise is therefore characterized by at least three quantities: its magnitude, its dependence on the current regulatory state, and its timescale relative to gene-expression dynamics.

The additive-noise phenotype generalized from small motifs to the 23-node EMT and 19-node gonadal cell-fate networks despite the change from an ODE to a Boolean formalism. In both networks, additive noise sharply reduced terminal-state occupancy and generated hybrid-dominated landscapes, whereas stationary and multiplicative noise caused comparatively little disruption. The specific hybrid states nevertheless differed: the EMT network retained partial polarization toward one of its two regulatory teams, whereas the gonadal network approached approximately equal activation of both teams. Thus, the tendency of additive noise to increase phenotypic mixing appears robust across network scales, while the particular hybrid configurations accessed remain constrained by network structure.

This result is especially relevant to epithelial–mesenchymal plasticity. Hybrid epithelial/mesenchymal (E/M) states are associated with stemness, collective migration, metastatic competence, and therapy resistance [35, 33, 41, 53]. Our results identify fluctuations in regulatory efficacy as one possible route to their enrichment: additive-like noise is sufficient to shift both an isolated bistable motif and a full EMT network toward hybrid phenotypes without changing network topology. Mutations affecting intrinsically disordered regions may therefore alter phenotype distributions not only through conventional gain- or loss-of-function effects, but also by changing the magnitude, persistence, or correlation structure of fluctuations in regulatory activity [54].

The connection between regulatory noise and phenotype is likely to extend beyond TF conformational dynamics to the chromatin environment in which regulatory interactions occur. Pluripotent and progenitor cells maintain relatively open, dynamic chromatin and broadly accessible transcriptional programs [55, 56]. Recent work further suggests that fluctuations in chromatin accessibility can actively enable ectopic or multi-lineage-like transcriptional programs [13]. Such a permissive chromatin state increases the number of genomic configurations accessible to TFs and may therefore enhance fluctuations in effective regulatory strength. At a coarse-grained level, this resembles the additive-noise regime: regulatory interactions explore a broader range of efficacies, weakening the confinement imposed by the deterministic network and increasing access to hybrid states.

Differentiation, however, is accompanied not simply by reduced transcriptional heterogeneity but by progressive stabilization of regulatory programs through chromatin remodeling and epigenetic feedback. This introduces a much slower form of memory than IDP conformational fluctuations. A useful example comes from EMT, where epigenetic feedback has been shown theoretically and experimentally to make phenotype transitions increasingly difficult to reverse. In earlier work on the miR-200/ZEB EMT circuit, prolonged TGF-*β* exposure progressively reduced the ability of cells to revert toward the epithelial state following withdrawal of the inducing signal, consistent with epigenetic feedback stabilizing the mesenchymal program [57, 58]. Such “locking” changes the problem from one of fluctuations around a regulatory state to one in which the history of the system progressively reshapes its future transition probabilities.

This suggests a hierarchy of regulatory dynamics. Fast IDP conformational fluctuations can continuously perturb interaction strengths; intermediate-timescale modifications can bias the conformational ensemble and introduce regulatory memory; and slower chromatin or epigenetic changes can stabilize the resulting transcriptional state. The same GRN may therefore behave differently without any change in topology depending on which timescale dominates. An open, rapidly exploring regulatory environment may favor additive-like dynamics and hybrid-state accessibility, whereas progressively correlated or self-reinforcing regulatory changes may reduce that exploration and facilitate commitment. Epigenetic locking can then consolidate the selected attractor over still longer timescales.

These observations suggest a two-stage picture of differentiation. In the first stage, *fate selection*, extracellular signals and the structure of the GRN bias a plastic or lineage-primed population toward particular regions of phenotype space. Regulatory variability need not oppose this process; in an open chromatin environment, additive-like fluctuations may facilitate exploration of multiple lineage-associated programs and maintain the hybrid states from which alternative fates remain accessible.

In the second stage, *fate resolution*, the selected lineage must be converted from a bias into a stable terminal phenotype. Our results suggest that this step need not require the instructive signal to specify every component of the final state. A shift away from broadly exploratory regulatory dynamics toward state-dependent reinforcement can itself favor single-high terminal configurations. Slower chromatin remodeling and epigenetic feedback can subsequently lock the resulting transcriptional program, making reversal progressively more difficult. Thus, an external signal may primarily determine *which* fate is selected, while changes in regulatory-noise regime and epigenetic memory determine *whether and how robustly* that choice is resolved.

This division provides a possible reconciliation of instructive and stochastic views of differentiation [59, 60]. Stochastic regulatory dynamics may dominate the exploratory phase, signals can bias the available alternatives, and state-dependent or epigenetic feedback can stabilize the resulting choice. The relevant control parameter would then not simply be the magnitude of molecular noise, but the evolution of its correlation time and state dependence as the cell progresses from plasticity to commitment.

Despite the novelty and the promising potential, the present study, like any other theoretical endeavour, does have some limitations. For instance, the present framework is deliberately phenomenological, and therefore does not establish a one-to-one mapping between the three noise modes and specific molecular processes. Relating these modes quantitatively to IDP conformational dynamics, chromatin-accessibility fluctuations, and slower epigenetic feedback will require experimental measurements and more mechanistically explicit models. A second limitation is that we consider regulatory-parameter noise independently of conventional expression-level noise. Because these two forms of stochasticity act on different components of the dynamical system, their combined effects may differ from either mechanism in isolation and warrant systematic investigation. Finally, our analysis focuses primarily on inhibition-dominated cell-fate decision networks. Whether the contrasting effects of additive and multiplicative regulatory noise extend to activation-rich networks, such as many signaling, metabolic, or protein–protein interaction networks, remains to be determined.

## 4 Methods

### 4.1 Network models and RACIPE ensemble generation

Each network motif is modeled as a system of ordinary differential equations (ODE), one ODE per node, following the shifted Hill formalism of RACIPE [31]:

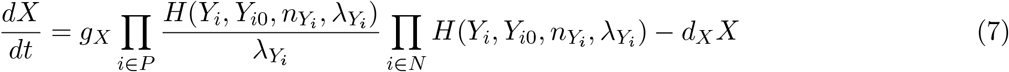

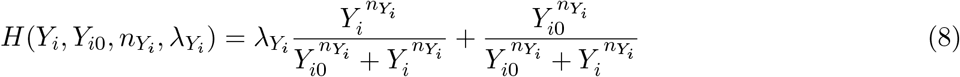

where *g*_*X*_ and *d*_*X*_ are the production and degradation rates of node *X, P* and *N* index its activating and inhibiting regulators, and *λ*_*Y*_ is the fold-change by which regulator *Y*_*i*_ modulates *X*’s production (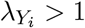 for activation, 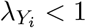 for inhibition).

For each network topology, RACIPE [31] was used to generate an ensemble of 10,000 parameter sets by uniform sampling within standard biologically plausible ranges, with each parameter set solved from 100 random initial conditions to identify its steady-state attractor(s). The resultant ensemble of steady states was used to calculate thresholds of individual node expressions (*θ*_*X*_) as the average expression across the ensemble, which were then used to discretize the steady states into strings of 1’s and 0’s. Based on the number of such discretized steady states allowed, each parameter set was assigned a multistability class (monostable, bistable, tristable, tetrastable). A representative subset of up to 1000 parameter sets was selected from each stability class present for a given network for downstream stochastic analysis (fewer where a stability class had fewer available parameter sets).

### 4.2 Regulatory noise model

To simulate regulatory noise, we allowed the fold-change parameters 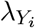 to vary stochastically over time rather than remaining fixed at their RACIPE-sampled value, using a periodic update applied every *dt* = 0.01 (a.u.) during ODE integration. Three noise modes were implemented, differing in how each update depends on the parameter’s current value:

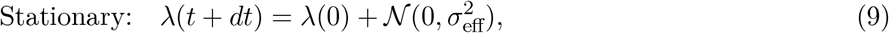

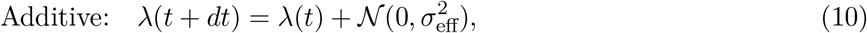

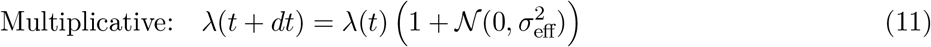

where *σ*_eff_ = *σ*·*λ*_max_, with *λ*_max_ the upper bound of the relevant fold-change (inhibitory: *λ*_max_ = 1, so *σ*_eff_ ≈ *σ*; activatory: *λ*_max_ = 100), so that a given nominal noise level *σ* corresponds to a comparable proportion of each parameter’s admissible range regardless of whether it governs activation or inhibition. After each update, *λ* was clamped to its biologically admissible bounds ([0.001, 0.999] for inhibitory, [1.001, *λ*_max_] for activatory parameters). Noise levels *σ* ∈ {0, 0.001, 0.005, 0.01, 0.05, 0.1, 0.5, 1} were tested for each noise mode and network.

### 4.3 Stochastic simulation and mean residence time

For each selected parameter set and noise level, 100 independent stochastic trajectories were simulated, each initialized from an independently drawn initial condition *u*_*i*_(0) = *U* (0, 1) *×* (*g*_*i*_*/d*_*i*_) for every node *i* (i.e. uniformly between zero and that node’s maximal attainable expression). Trajectories were integrated over *t* ∈ [0, 1000] (a.u.) using the Tsit5 adaptive-step solver (relative tolerance 10^−4^, absolute tolerance 10^−5^). Each trajectory was discretized into a sequence of Boolean node states using the RACIPE-derived thresholds, and the first 50% of each trajectory was discarded as transient. Mean residence time (MRT) for a given discrete state was computed as the fraction of the remaining trajectory (across all 100 stochastic realizations) spent in that state.

### 4.4 Deterministic *λ*-resampling comparison

To test whether a given noise mode’s effect on state occupancy could be explained purely by the stationary distribution of *λ* it induces, rather than by any dynamical (non-adiabatic) contribution, we compared stochastic MRT against a deterministic control: for each parameter set and noise level, *λ* values were resampled from the empirical stationary distribution generated by that noise mode (via `get_lambda_dist`), and the corresponding deterministic steady state was solved; this was repeated over 10 resampled draws per parameter set, and the resulting basin-occupancy fractions (MRT_det_) were compared directly against the stochastic MRT from the same parameter set and noise level.

### 4.5 Boolean simulations of large cell-fate networks

For larger networks, we used Boolean modeling to simulate network dynamics, as opposed to the ODE models of RACIPE. We used a threshold-based Boolean formalism [50], inspired by the Ising model of spin-glass systems. A network is defined by *N* nodes, *E* edges, and a topology, i.e., the arrangement of *E* edges among *N* nodes. The Ising model employs discretized expressions of nodes, with *s*_*i*_ ∈ {−1, 1}; *i* ∈ 1, …, *N*, where *s*_*i*_ = 1 and *s*_*i*_ = −1 represent an active node and an inactive node, respectively. A state at any time *t* is the set of expression levels of all nodes, *S*(*t*) = {*s*_*i*_(*t*) }; *i* ∈ 1, …, *N* .

The system state is updated in discrete time steps using an asymmetric adjacency matrix (Adj), which stores the nature of pairwise interactions between nodes. In a static network, activating and inhibiting edges carry edge weights of +1 or −1, respectively, with 0 indicating no interaction. The expression level of node *I* at time *t* is given by:

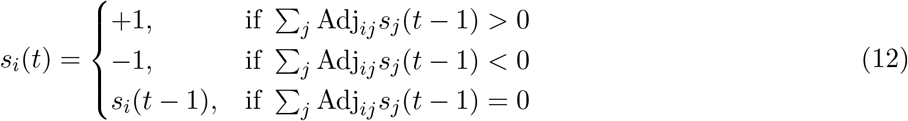

where ∑_*j*_ Adj_*ij*_*s*_*j*_ represents the net regulatory input to node *i*. A steady state is defined as a state *S*(*t*) satisfying *S*(*t* + 1) = *S*(*t*).

### 4.6 Large network metrics

#### 4.6.1 Influence matrix

The influence matrix is formulated as:

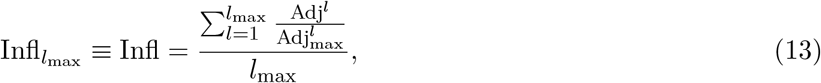

where *l* is the path length, representing a path of *l* edges connecting a pair of nodes, and *l*_max_ = 10 is the maximum path length considered. The influence matrix captures the net bidirectional effect between a pair of nodes mediated by all paths of length *l* ∈ {1, …, *l*_max_}. 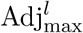 is derived from Adj^*l*^ by setting all non-zero entries to 1, and the division 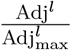 is element-wise.

#### 4.6.2 Teams

The wild-type influence matrices of the biological networks considered in this study are displayed in Figure S13. Each network can be partitioned into two “teams” of nodes, defined such that the influence between any pair of nodes is positive if they belong to the same team and negative otherwise. The team strength of a two-teamed network is defined as:

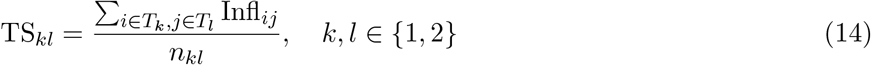

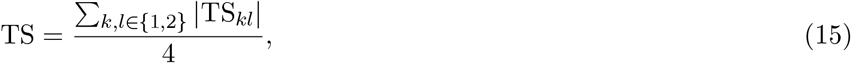

where *T*_*k*_ is the *k*-th team.

#### Team expression

For a given state *S*, team expression is defined as the fraction of nodes belonging to a team that are active (i.e., *s*_*i*_ = 1).

#### Nature of stable states

Stable states of two-team networks are classified as either dominant (most frequent) or hybrid (non-dominant). Due to the bimodal phenotypic distribution characteristic of toggle switch-like networks, dominant states are those in which only one team is active, i.e., states with team expressions of (0, 1) and (1, 0).

### 4.7 Stochastic simulation of large networks

#### 4.7.1 Inducing noise

In biological networks, the strengths of regulatory interactions are subject to temporal variation arising from fluctuations in protein concentrations, post-translational modifications, and other molecular sources. We model this regulatory noise by perturbing the edge weights of the adjacency matrix over time. Three noise modes are implemented, mirroring those used in the ODE-based simulations of small motifs (Section 4.1):

##### Additive (state-independent) noise

At each time step, a perturbation drawn from *N*(0, *σ*^2^) is added to each edge weight independently of its current value:

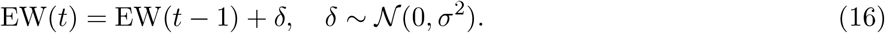

Edge weights are clipped to [−1, 1] after each update to preserve the biological interpretation of interaction strengths.

##### Multiplicative (state-dependent) noise

At each time step, each edge weight is scaled by a factor drawn from *N* (1, *σ*^2^), so that the magnitude of the perturbation scales with the current edge weight:

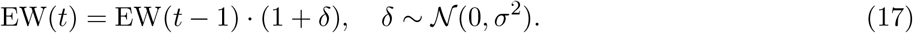

##### Stationary (mean-reverting) noise

At each time step, each edge weight is redrawn as its wild-type value plus an independent perturbation, so that the weight fluctuates around its nominal value without drift:

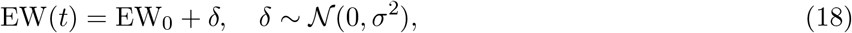

where EW_0_ denotes the wild-type edge weight. Unlike the additive mode, this formulation prevents long-range drift in parameter values and models a situation where molecular fluctuations are transient and the nominal interaction strength is effectively restored between perturbation events.

In all three modes, *σ*^2^ = 0.01 is used as the noise variance. Each biological network is simulated for 1000 trajectories of 10,000 time steps each; artificial networks are simulated for 100 trajectories of 1,000 time steps each.

#### 4.7.2 Method of state update

In practice, the system can be updated in two ways [61, 62]:

- **Synchronous update**: At each time step, every node is updated simultaneously using Eq. 12.
- **Asynchronous update**: At each time step, a single randomly chosen node is updated using Eq. 12.

While synchronous updates are computationally efficient, they can produce limit cycles from which the system cannot escape. Asynchronous updates are less efficient but avoid limit cycles and implicitly capture reaction noise by assuming that only one regulatory event occurs per time step. To combine the benefits of both schemes, we check for steady states synchronously every 200 time steps while updating the network asynchronously. For continuous perturbation simulations, synchronous updates are used throughout, as persistent perturbations make entrapment in limit cycles unlikely.

#### 4.7.3 Initial conditions

Peripheral nodes — those involved in interactions without feedback — are excluded from the analysis, as the focus of this study is on teaming behavior arising from mutual regulatory effects. Initial conditions are drawn from the terminal states and the most dominant hybrid states identified in the unperturbed network’s steady-state distribution.

## 5 Acknowledgments

MKJ was supported by the Ramanujan Fellowship (SB/S2/RJN-049/2018) awarded by the Science and Engineering Research Board (SERB), Department of Science and Technology, Government of India. MKJ was also supported by Param Hansa Philanthropies. KH and HL were supported by the Center for Theoretical Biological Physics, NSF PHY-2019745, and under Award Number MCB-2114191.

## 6 Data Availability

The code used to generate and analyze the simulations reported in this study, including the RACIPE-based stochastic simulation pipeline and all figure-generation scripts, is available at github.com/askhari139/Regulatory-noise-model (this repository, in the `codes/` folder). Raw simulation output data are available from the corresponding author upon reasonable request.

## S1 Derivation of the shifted Hill functions

A coarse-grained model of transcription and translation is used to describe gene regulatory networks. The shifted Hill function captures a non-zero basal expression rate together with a regulated expression rate that is higher (activation) or lower (repression) than the basal level. The derivation rests on the following assumptions.

### S1.1 Assumptions

1. *Quasi-steady-state binding:* DNA–TF binding and unbinding occur much faster than transcription and translation.
2. *TF in excess:* the transcription factor is abundant relative to the DNA, so its free concentration is approximately its total concentration.

### S1.2 Single-regulator model

For a single regulator (TF), the reactions producing protein *P* through transcription and translation of the target gene are

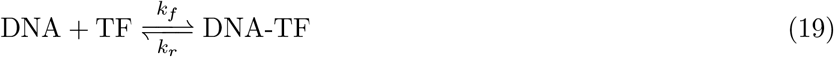

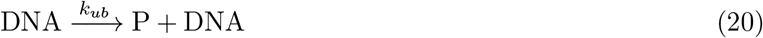

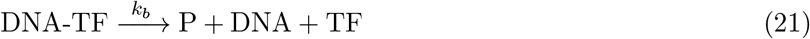

Reaction (19) describes formation of the DNA–TF complex. Reactions (20) and (21) together account for protein production: for an activating TF, *k*_*b*_ *> k*_*ub*_, and for an inhibiting TF, *k*_*ub*_ *> k*_*b*_. The corresponding production rate is

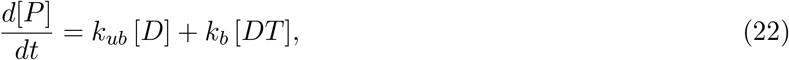

where [*D*] is the concentration of free DNA and [*DT*] that of the DNA–TF complex. The first term on the right-hand side is the basal expression and the second is the regulated expression (with the roles of “basal” and “regulated” set by whether the link activates or inhibits).

Applying the quasi-steady-state assumption to TF binding,

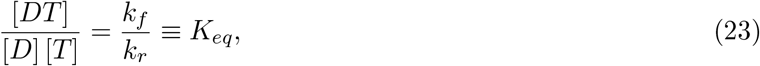

where [*T*] is the free TF concentration. Because the TF is in excess, [*T*] equals the total TF concentration. Conservation of DNA gives

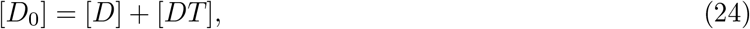

with [*D*_0_] the (constant) total DNA concentration. Substituting Eq. (23) into Eq. (24),

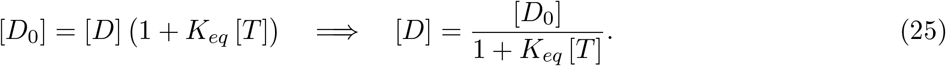

Using [*DT*] = [*D*_0_] − [*D*], Eq. (22) becomes

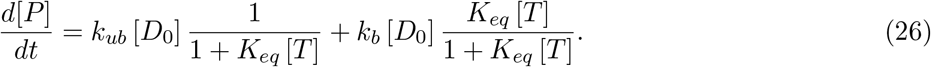

Defining the unbound and bound occupancy fractions

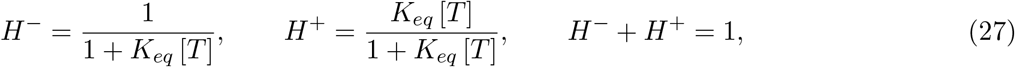

Eq. (26) reads

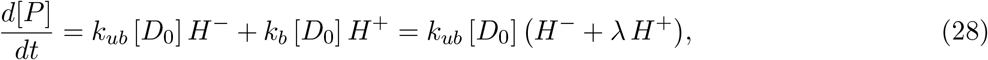

where *λ* ≡ *k*_*b*_*/k*_*ub*_.

Let *g* denote the maximal protein production rate. For an inhibitory link the maximum is the unbound rate, *g* = *k*_*ub*_ [*D*_0_]; for an activating link it is the bound rate, *g* = *k*_*b*_ [*D*_0_] = *λ k*_*ub*_ [*D*_0_]. Normalizing Eq. (28) by *g* therefore yields the two branches of the shifted Hill function,

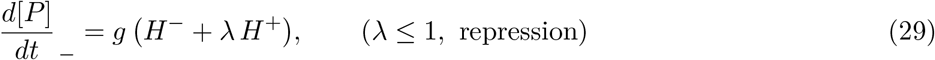

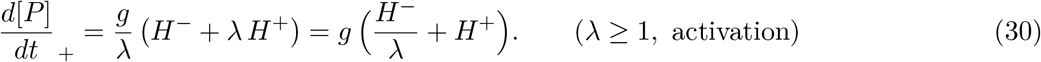

The two regulatory constants entering the production term are thus

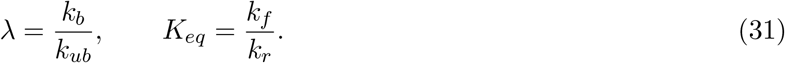

Here *λ* is the ratio of the transcription rate with the TF bound to that with it unbound—possibly set by TF-mediated condensate formation or RNA polymerase recruitment—while *K*_*eq*_ is the ratio of the TF’s DNA binding and unbinding rates. The half-maximal regulator concentration (threshold) follows from 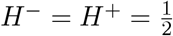, i.e. [*T*]_1*/*2_ = 1*/K*_*eq*_.

#### Remark (continuity and cooperativity)

Both branches coincide at *λ* = 1, where *H*^−^ + *H*^+^ = 1 and production equals the maximal rate *g* independent of [*T*]: the no-regulation limit. Cooperative binding of *n* TF molecules replaces *K*_*eq*_[*T*] by *K*_*eq*_[*T*] ^*n*^ in Eq. (27), giving the Hill coefficient *n* and recovering the usual sharp (switch-like) form used in the simulations.

## S2 A Boolean limit explains the noise-driven enhancement of the double-positive state

To isolate the mechanism by which structural (*λ*) noise redistributes occupancy toward high-expression states, we analyse the toggle switch in the Boolean limit, where the exact stationary distribution can be computed in closed form for both the deterministic (fixed-*λ*) and stochastic (fluctuating-*λ*) cases.

### S2.1 The two *λ* extremes are not symmetric

For an inhibitory edge, the shifted Hill function entering the production term is

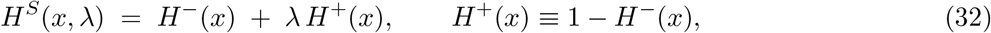

which is *exactly affine in λ* for fixed regulator level *x*. At the noise amplitude *σ* = 1 the clamped *λ* walk is confined to the two boundaries of its range, *λ* ∈ {*λ*_min_, *λ*_max_} with *λ*_min_ ≈ 0.01 and *λ*_max_ = 1. Evaluating Eq. (4) at these two limits exposes a structural asymmetry:

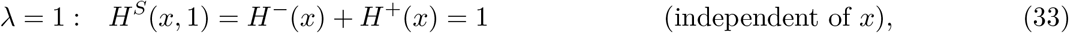

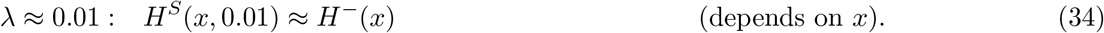

The *λ* = 1 boundary abolishes regulation *unconditionally*: the target is produced at its full rate regardless of the regulator’s expression. The *λ* ≈ 0 boundary imposes repression only *conditionally*: it lowers the target’s production rate only when the regulator *x* is high, and has essentially no effect when the regulator is low (*H*^−^(*x*) ≈ 1). An excursion toward de-repression therefore always “fires,” whereas an excursion toward repression fires only when the partner node happens to be expressed. This unconditional-up / conditional-down asymmetry is the microscopic origin of the enhancement derived below.

### S2.2 Boolean reduction

We coarse-grain expression to (*A, B*) ∈ {0, 1}^2^ and each edge’s regulatory strength to *λ*_1_, *λ*_2_ ∈ {0, 1}, where *λ* = 1 denotes the unconditional (de-repressed) branch and *λ* = 0 the conditional (repressive) branch. In the Boolean limit the fixed-*λ* update rule is

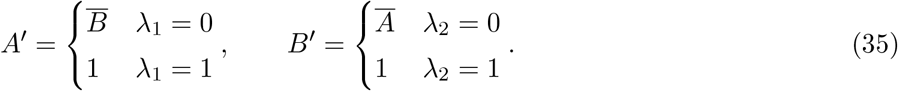

Each of the four *λ*-configurations defines a deterministic map on the four states (00, 01, 10, 11), with transition matrices

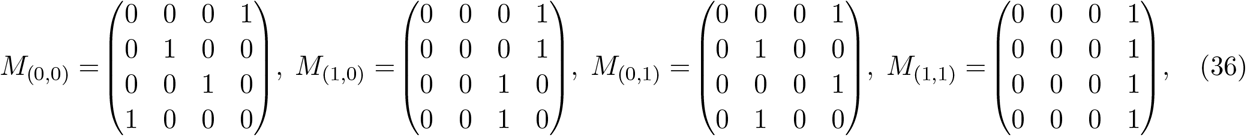

in row/column order (00, 01, 10, 11).

### S2.3 Deterministic case (*σ* = 1, *λ* drawn once and held fixed)

In the deterministic (quenched) picture each parameter set draws a single (*λ*_1_, *λ*_2_) ∈ {0.01, 1}^2^ and integrates to its attractor(s). The four configurations yield:

| $(\lambda_1, \lambda_2)$ | Effective dynamics | Attractor(s) |
| --- | --- | --- |
| $(1, 1)$ | $A, B$ both unconditionally on | $\{11\}$ |
| $(1, 0)$ | $A$ on; $B = \bar{A}$ | $\{10\}$ |
| $(0, 1)$ | $B$ on; $A = \bar{B}$ | $\{01\}$ |
| $(0, 0)$ | mutual inhibition (toggle) | $\{10, 01\}$ |

The (0, 0) configuration is the bistable toggle, whose stable attractors are the two single-positive fixed points {10, 01}; the states 00 and 11 are unstable and carry no basin. Weighting the four configurations equally (each boundary combination is equiprobable at *σ* = 1) and splitting the bistable config’s weight symmetrically between its two attractors gives the deterministic stationary distribution

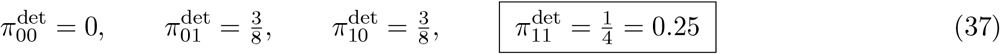

The double-positive state 11 is reached only under the single (1, 1) configuration; the conclusion 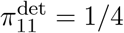 is independent of how the bistable config’s basins are partitioned, since (0, 0) never resolves to 11.

### S2.4 Stochastic case (*λ* fluctuating)

When *λ* fluctuates rapidly relative to the state dynamics, the effective one-step operator is the *λ-average* of the four deterministic maps in Eq. (36):

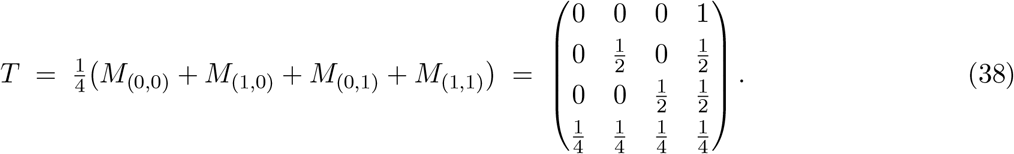

The chain is irreducible and aperiodic 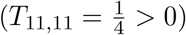, so a unique stationary distribution exists and equals the long-run occupancy. Solving *π* = *πT* with ∑_*i*_*π*_*i*_ = 1 gives 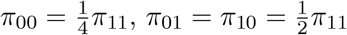, hence 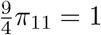 and

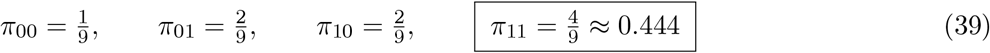

The mean residence time in the double-positive state is MRT(11) = *π*_11_ = 4*/*9.

### S2.5 Every state flows toward 11 under fluctuating *λ*

The enhancement is read directly off the last column of *T* (Eq. (38)): the one-step probability of entering 11 is positive from *every* state,

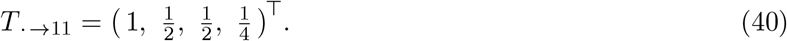

In particular the fully-repressed state 00 maps to 11 with probability 1: under the update rule (35), when neither node is expressed both targets are de-repressed under *all four λ*-configurations (the *λ* = 1 branch sets the target high unconditionally, and the *λ* = 0 branch sets it high because the repressor is absent), so 00 → 11 is *λ*-independent. By contrast, in the deterministic case 11 is reachable only under the isolated (1, 1) configuration. Fluctuating *λ* thus lets a parameter set that would deterministically settle into a single-positive fixed point (01 or 10) nonetheless accumulate occupancy in 11, because every re-draw of *λ* offers a fresh, largely unconditional route upward.

### S2.6 Comparison

| State | Deterministic $\pi^{\text{det}}$ | Stochastic $\pi$ | Change |
| --- | --- | --- | --- |
| 00 | 0 | $1/9 \approx 0.111$ | — |
| 01 | $3/8 = 0.375$ | $2/9 \approx 0.222$ | ↓ |
| 10 | $3/8 = 0.375$ | $2/9 \approx 0.222$ | ↓ |
| 11 | $1/4 = 0.250$ | $4/9 \approx 0.444$ | ↑ (1.78×) |

Fluctuating *λ* increases the occupancy of the double-positive state by a factor of 16*/*9 ≈ 1.78 relative to the deterministic prediction from the same clamped *λ* range, drawing the redistributed mass from the two single-positive states. This is a purely structural effect: the two calculations use an identical, symmetric *λ* ensemble confined to the same boundaries, and differ only in whether *λ* is quenched (Eq. (37)) or annealed (Eq. (39)). The gain reflects the asymmetry of Section S2.1—de-repression fires unconditionally while repression fires only conditionally—so that mixing over *λ* favours the state in which both nodes are high.

**Remark**. The small stationary occupancy of 00 (*π*_00_ = 1*/*9) arises from the synchronous discrete update, in which the (0, 0) configuration maps 11 → 00; in the continuous ODE this appears as a brief transient rather than a stable state. The robust, ODE-relevant conclusion is the redistribution of occupancy from the single-positive states to the double-positive state, not the absolute occupancy of 00.

## S3 Static Perturbation analysis of large-scale binary cell-fate decision networks

In this section, we investigate the impact of static perturbations (to edge weights and density; see §4.7.1) on phenotypic distributions, team strengths, and their interrelation. First, we qualitatively understand the effect of noise on the steady-state distribution. We will display results for one artificial and biological network each, and provide reference to the corresponding section in the supplementary material for other networks. Fig. **??** provides the frequency of (steady-state) phenotypic distributions of EMT(23N, 89E) (top) and ToggleSwitch(10N, 20E) (bottom) for three different noise levels (legend; arranged in decreasing order of noise level) of static perturbations to edge weights (left) and density (right). The dots represent the mean of the frequencies of the state across the iterations they have occurred. The upper and lower error bars represent the 75-th and 25-th percentile of this distribution, respectively. We characterize the steady states by the difference between the team expressions (see §4.6.2) of the bigger and the smaller teams, i.e., T1 − T2, as shown in the x-axis. The black dashed line belongs to the corresponding wild-type network.

Our general qualitative observations are: One, the dominant state in the unperturbed network remains dominant in perturbed networks up to TS_wild_ = 0.17 for biological networks and TS_wild_ = 0.61 for artificial networks, below which the state with T1 − T2 in the wild-type network closer to 0 becomes dominant. In the latter case, prediction of the exact dominant state across noise levels isn’t possible as they occur non-uniformly. Two, the number of hybrid states (states with team expressions that are neither (0, 1) nor (1, 0)) increases with an increase in noise level. Three, the states that occurred in unperturbed or relatively less perturbed networks continue to occur in networks with higher perturbations. Four, there is no pattern in the frequencies for a given hybrid state across changes in noise levels. The above observations are common regardless of the network-specific nuances and the differences between the artificial and biological networks. The one contrast between artificial and biological networks is that the dominant states are always the pure states in unperturbed artificial networks, whereas, in biological networks, there are some networks with dominant states that are not pure even for TS_wild_ above the thresholds mentioned above. This is probably because of the higher interconnectivity and smaller size of artificial networks than biological networks. The frequency of dominant states decreases with increasing noise in all cases except for biological networks subject to edge-weight perturbations. Another observation for perturbed networks with wild-type dominant states not being pure is that the frequency of pure states increases with increasing noise at the expense of the dominant state frequency. In conclusion, for networks with team strength above a threshold, the teaming behavior supports the unperturbed low-dimensional phenotypic distribution. In the absence of a ‘strong’ teaming behavior, the network tries to behave as one unit.

### S3.1 Static Edge-weight Perturbations

In this section, we quantify our qualitative observations for edge-weight perturbations, as summarized in Fig. **??**. We obtain the mean of team strength distributions (TS) from the iterations (see App. **??**) and normalize them by the corresponding team strength of the unperturbed (wild-type) network (TS_wild_). To quantify TS*/*TS_wild_ as a function of edge-weight noise levels, we fit the data to an empirical model defined as:

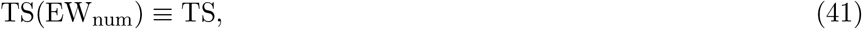

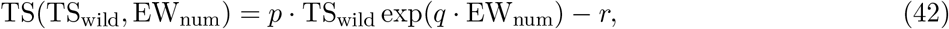

where EW_num_ is the numerical value of a given edge-weight range (see Eq. 47; §**??**). The team strength values obtained from the resulting fits are plotted (as black crosses) along with TS (blue dots) against the corresponding edge-weight ranges, across networks, as shown in the top-left panel in Fig. **??**. The edge-weight ranges (x-axis) are arranged in the ascending order of the noise level they introduce, i.e., the EW_num_. The black-dashed line is Eq. 42 calculated over the edge-weight noise range. The fit parameters are shown in Tab. **??**. We emphasize that the fit parameters aren’t independent of each other and hold no physical property. This demonstrates that the teams’ resilience to static edge-weight perturbations is attributed to its underlying team-based topology of the network, quantified using team strength. A natural expectation is that the emergent phenotypes from teams must also be resilient to static edge-weight perturbations, with the latter increasing with team strength, and we test this below.

The middle and bottom panels in Fig. **??** correspond to an artificial and a biological network, respectively. We use JSD to quantify the divergence of phenotypic distribution of the perturbed network from the corresponding wild-type network, with greater JSD implying greater divergence. Therefore, higher Jensen-Shannon Divergence (JSD) marks a lower resilience of the network to the introduced noise. (see §**??**). The top-right panel in Fig. **??** shows the JSD distributions across edge-weight noise levels. We see a general trend in the variance of the JSD distributions, increasing with increasing edge-weight noise level. We are interested in the role of team strength on JSD in the presence of perturbations. Therefore, we propose an empirical model to establish a relationship between JSD and TS for a given network, defined as:

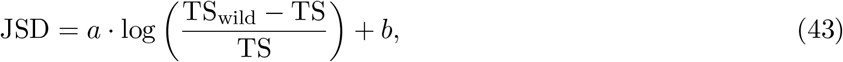

where *b* = JSD(TS = TS_wild_*/*2). Here, JSD refers to the mean of the JSD distribution, unless mentioned otherwise. We fit Eq. 43 to JSD and TS*/*TS_wild_ across edge-weight ranges (legend) as shown in the middle and bottom-left panel in Fig. **??**. The black dashed line interpolates Eq. 43 over the range of TS*/*TS_wild_, i.e., [0, 1]. Note that the orange dots at the extremes are not points obtained from our simulation but correspond to JSD(TS = 0) = 1 and JSD(TS = 1) = 0 according to Eq. 43. Tab. **??** displays the obtained parameters *a* and *b* from LMFIT along with their 1-sigma variations for each network. As expected, *b* decreases with an increase in TS_wild_. We see no trend in *a* as the first term on the right of Eq. 43 is independent of TS_wild_ (see Eq. 42). Thus, we can conclude that, for a given network, the resilience of the phenotypic distributions to static edge-weight perturbations increases with increasing team strength.

To check if the models Eqs. 42 and 43 can predict the dependence of JSD on TS across networks for a particular edge-weight range, Eq. 42 is plugged into Eq. 43 to result the model:

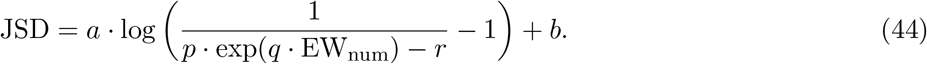

The JSD-values along with their error-bars are calculated for constant EW_num_s using parameter values from Tabs. **??** and **??**. These are displayed alongside JSD (obtained from our simulations) plotted against TS*/*TS_wild_ for a particular edge-weight range in the middle and bottom-right panel in Fig. **??**. We emphasize that no further fitting is done here, and Eq. 44 predicts our data within 2-sigma variations. While JSD decreases with an increase in TS_wild_ for artificial networks as expected, the trend is the opposite for biological networks. This observation remains unexplained.

## S4 Methods

### S4.1 Jensen–Shannon divergence

Jensen–Shannon divergence (JSD) is a symmetric, bounded measure of similarity between two probability distributions [63], derived from the KL divergence:

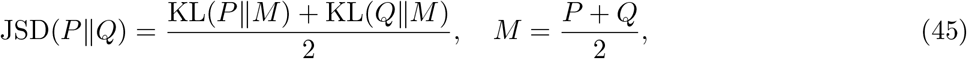

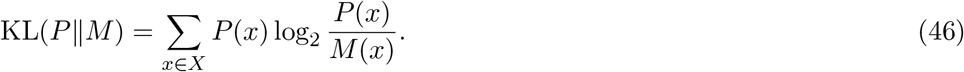

JSD takes values in [0, 1], with JSD = 0 indicating identical distributions. It is used here to quantify the divergence of the perturbed network’s phenotypic distribution from the wild-type, serving as a quantitative measure of network resilience: higher JSD implies lower resilience.

## S5 Fitting and error propagation

LMFIT [64] is used for model fitting. Error bars (1*σ* intervals) are calculated using Monte Carlo error propagation, which does not require the assumption that the distributions under consideration are Gaussian. Parameter values obtained from LMFIT are treated as Gaussian, and random samples drawn from these distributions are propagated through the model to generate a distribution for each derived quantity. The mean, median, and 16th and 84th percentiles are extracted from the resulting distribution; the upper and lower error limits are defined as the 84th percentile minus the median and the median minus the 16th percentile, respectively.

### S5.1 Static edge weight perturbation

To complement the continuous noise simulations and characterise the steady-state sensitivity of network phenotypes to fixed changes in interaction strength, we also performed static perturbation experiments. For each simulation run, a weight for each edge was independently sampled from a uniform distribution within one of the following ranges: [0, 0.25], [0, 0.5], [0, 0.75], [0, 1], [0.25, 1], [0.5, 1], [0.75, 1], and [0.95, 1], ordered from highest to lowest noise level. The topology and sign of interactions were held constant across all perturbations.

#### Assigning numerical values to edge-weight ranges

As the smallest interval considered is 0.05, the range [0, 0.05] is assigned value 1. The numerical value for an arbitrary range is defined as:

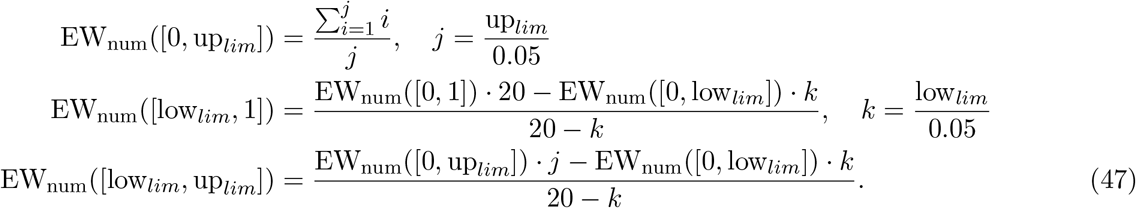

Note that this metric is undefined when low_*lim*_ = up_*lim*_.

##### Initial conditions

For networks with *N <* 20, all 2^*N*^ possible initial states are enumerated. For *N* ≥ 20, 2^20^ randomly generated initial states are used. In both cases, an unperturbed (wild-type) simulation with edge-weight moduli of 1 is run as a baseline against which phenotypic distributions under perturbation are compared.

## S6 Distribution of *λ* under additive noise

### Distribution of Additive noise *X* ∼ Uniform[0, 1], *Z* ∼ *N* (0, *σ*^2^)

#### S6.0.1 Notations

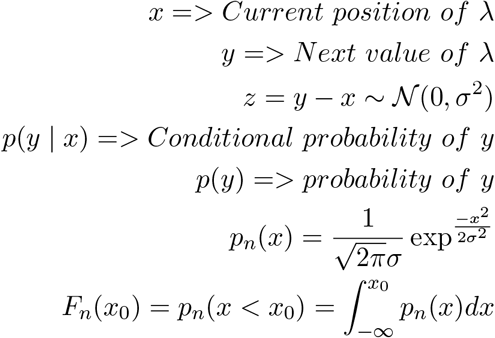

##### Conditional distribution and continuous density

For *y* − *x* = *z* ∼ *N* (0, *σ*^2^):

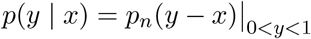

##### Point masses

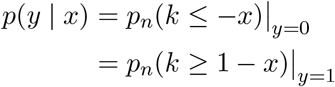

##### Marginal density for 0 *< y <* 1

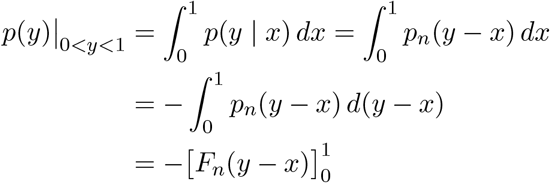

Hence:

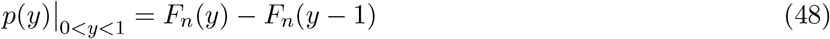

##### Point masses via integration by parts

**Mass at** *y* = 0:

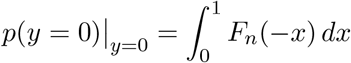

**Mass at** *y* = 1:

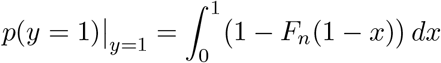

**Substitution** *u* = 1 − *x, du* = −*dx*:

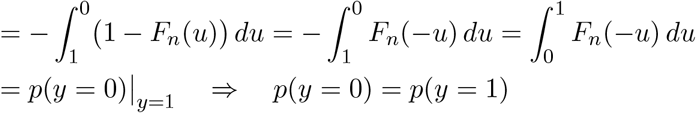

**IBP:** ∫*u dv* = *uv* − ∫*v du*, with *u* = *F*_*n*_(−*x*), *v* = *x*:

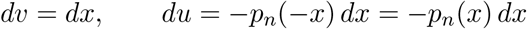

##### Gaussian identity

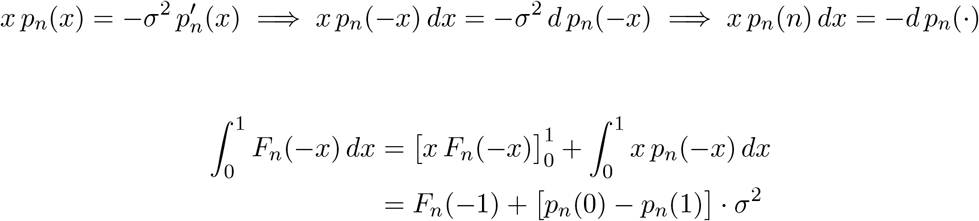

Hence:

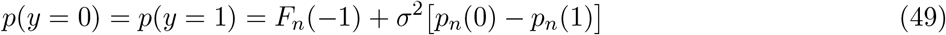

**CDF for** 0 ≤ *y*_0_ ≤ 1

Note: *F*_*n*_(*x*_0_) = *P*_*n*_(*x* ≤ *x*_0_).

**CDF:**

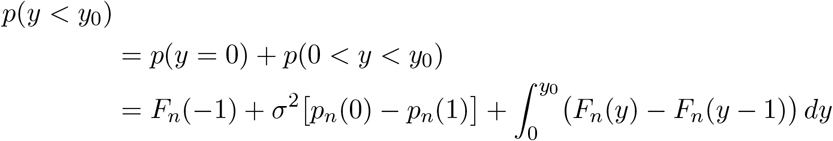

**Antiderivative of** *F*_*n*_ (IBP):

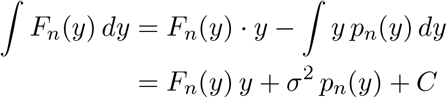

**Definite integral:**

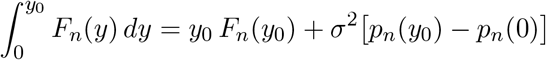

**Shifted integral** via *u* = *y* − 1, *du* = *dy, y* = 0 ⇒ *u* = −1, *y* = *y*_0_ ⇒ *u* = *y*_0_ − 1:

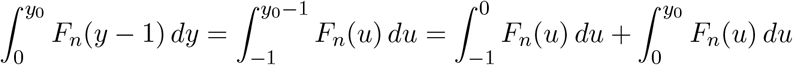

Therefore:

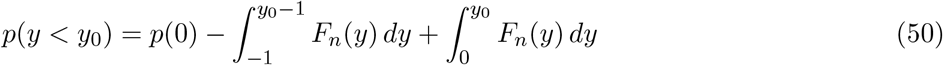

##### Evaluating the CDF explicitly

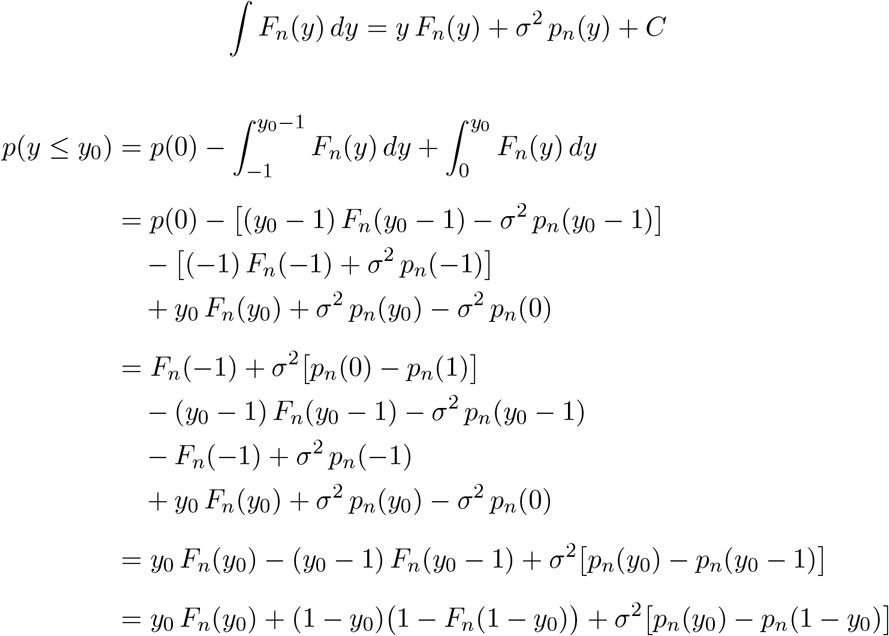

##### Final form and complement

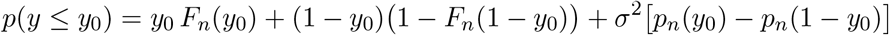

**Complement** *P* (*y*_0_ ≤ *Y* ≤ 1):

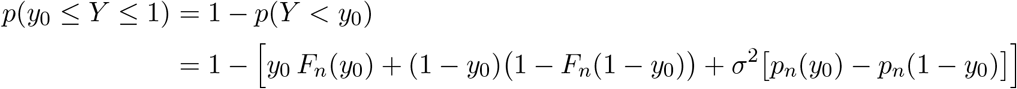

**Substitution** *y*_0_ = 1 − *y*′:

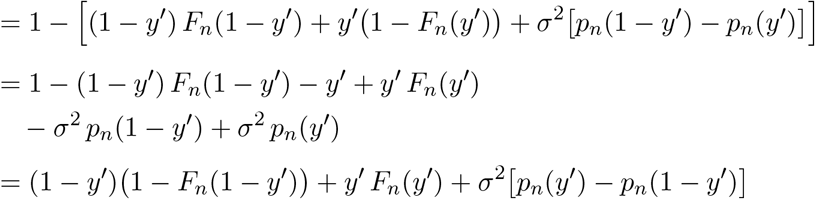

##### Summary and numerical example

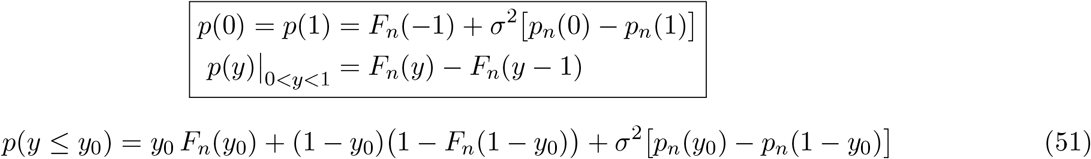

**Note:** this also equals *p*(*y*_0_ ≤ *Y* ≤ 1) by symmetry.

## S7 Supplementary Figures

**Figure S1:**
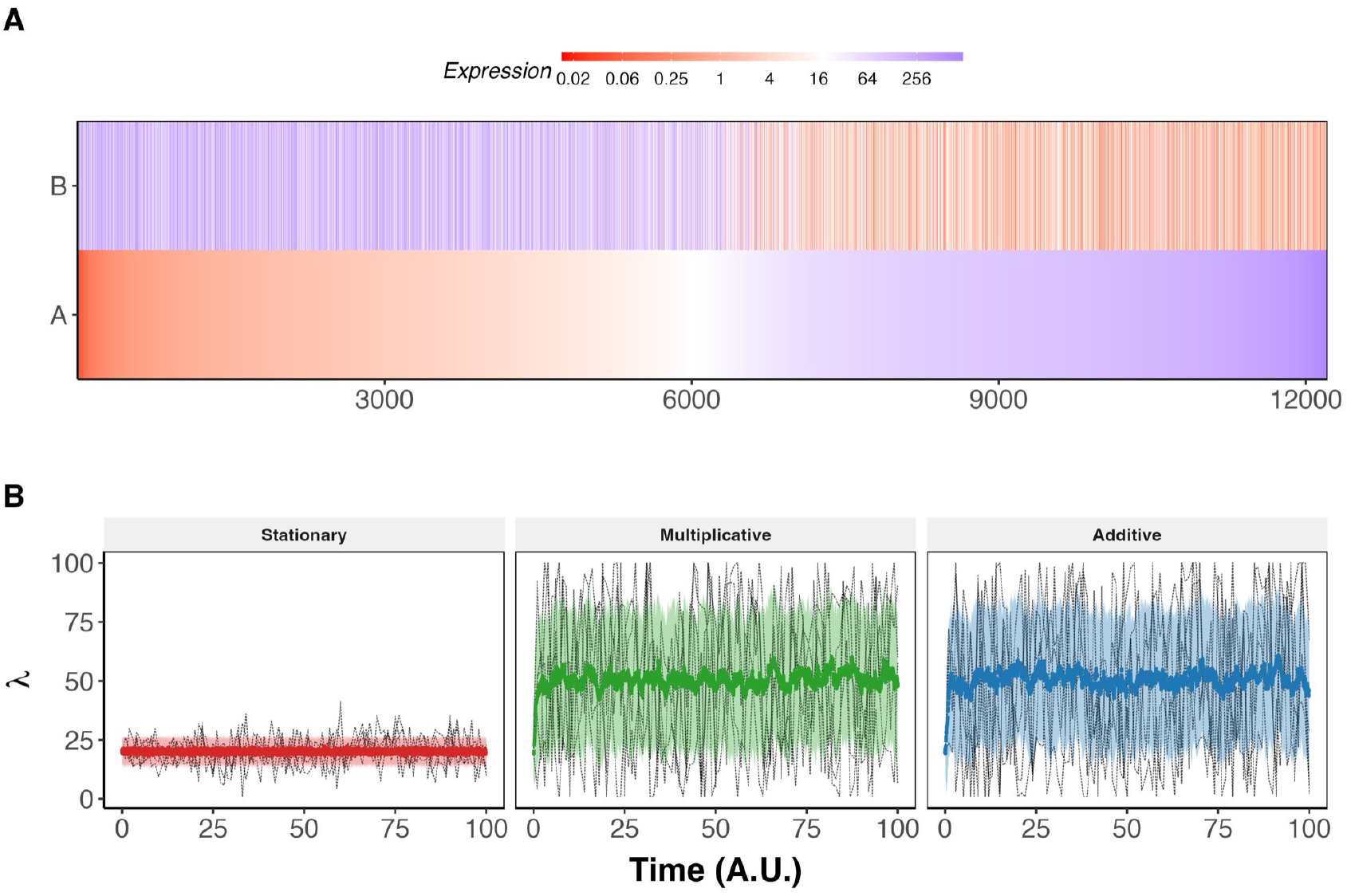
**(A)** Heatmap of RACIPE steady-state expression for nodes A and B in TS network, across all sampled parameter combinations. The steady states are sorted in increasing order of expression of node A. The color scale is in *log*_2_ scale and centered at the mean of the two nodes’ thresholds. **(B)** Same design as Figure 1D, but for a fold-change of an activatory edge (*λ* ∈ [1, 100], *λ*_0_ = 20).

**Figure S2:**
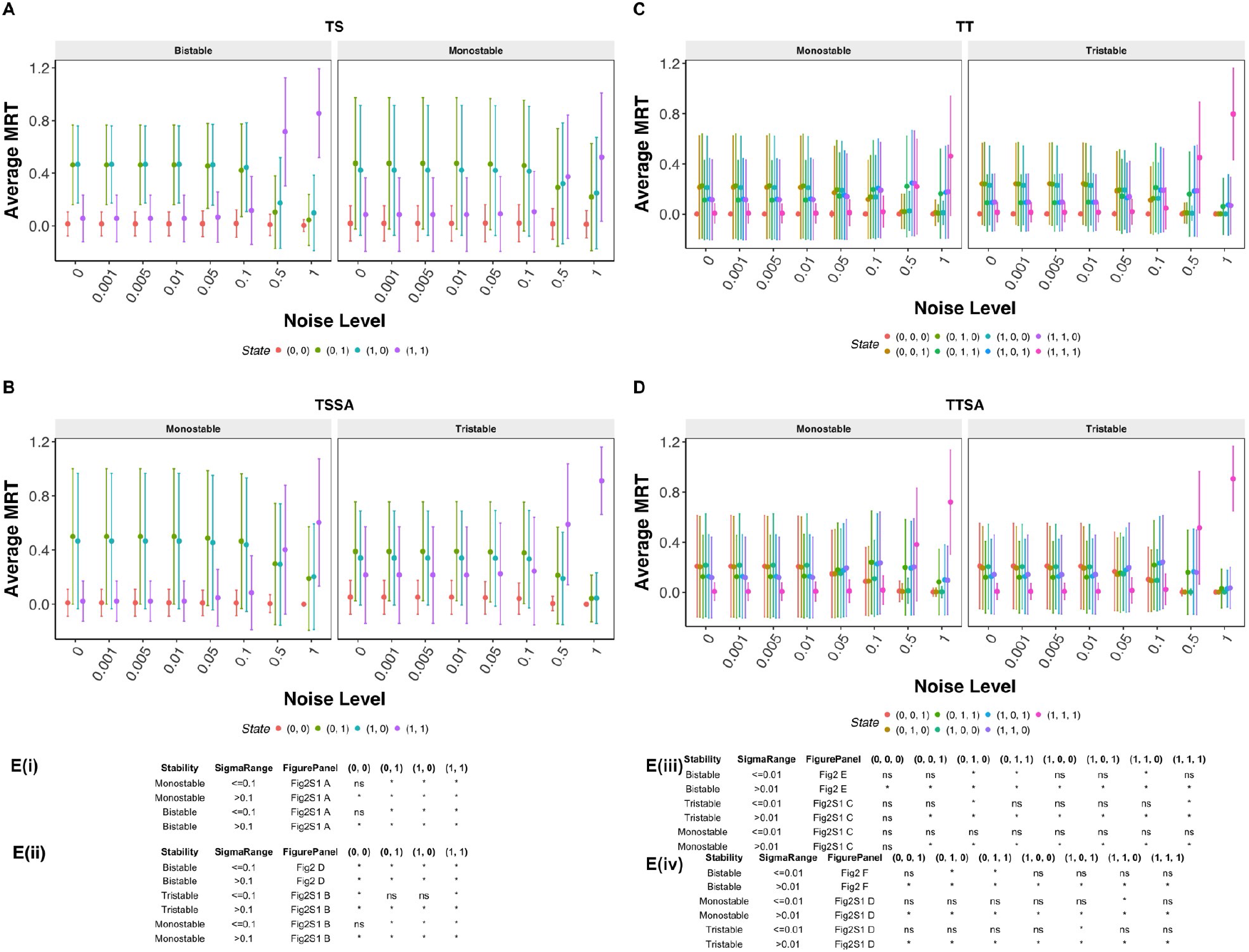
MRT vs. noise level, colored by discrete state, mean *±* SD across parameter sets, for **(A)** TS (all parameter sets), **(B)** TSSA (monostable + tristable), **(C)** TT (monostable + tristable), and **(D)** TTSA (monostable + tristable).

**Figure S3:**
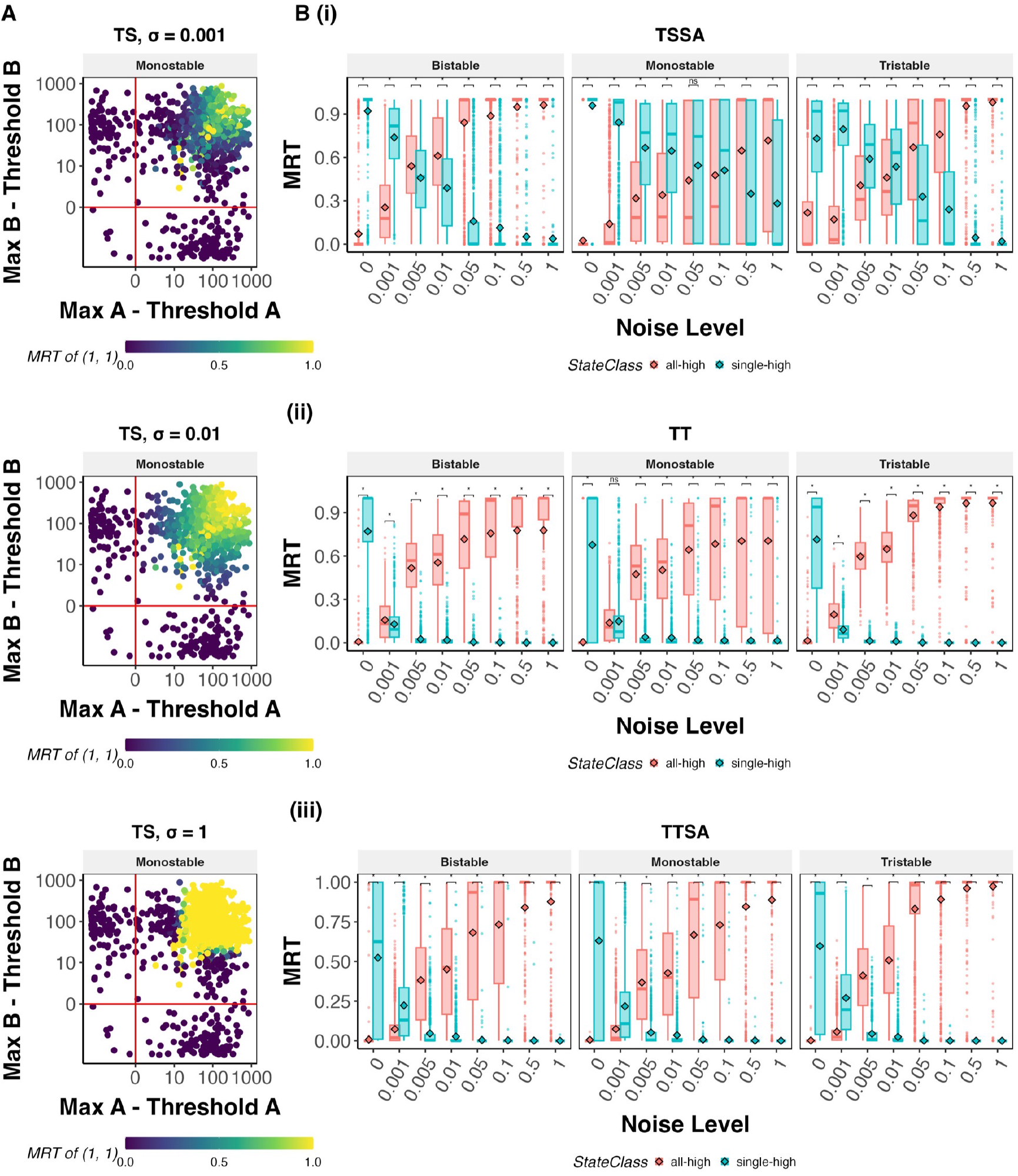
**(A)** MRT boxplots (as in Fig. 3B) for TSSA, TT, and TTSA, stacked vertically. **(B)** Reachability scatter (as in Figure 3C) for the TS network at *σ* = 0.001, 0.01, 1.0 (the noise levels not shown in the main figure), stacked vertically. **(C)** Reachability scatter for the TSSA network at *σ* = 0.001, 0.01, 0.1, 1.0.

**Figure S4:**
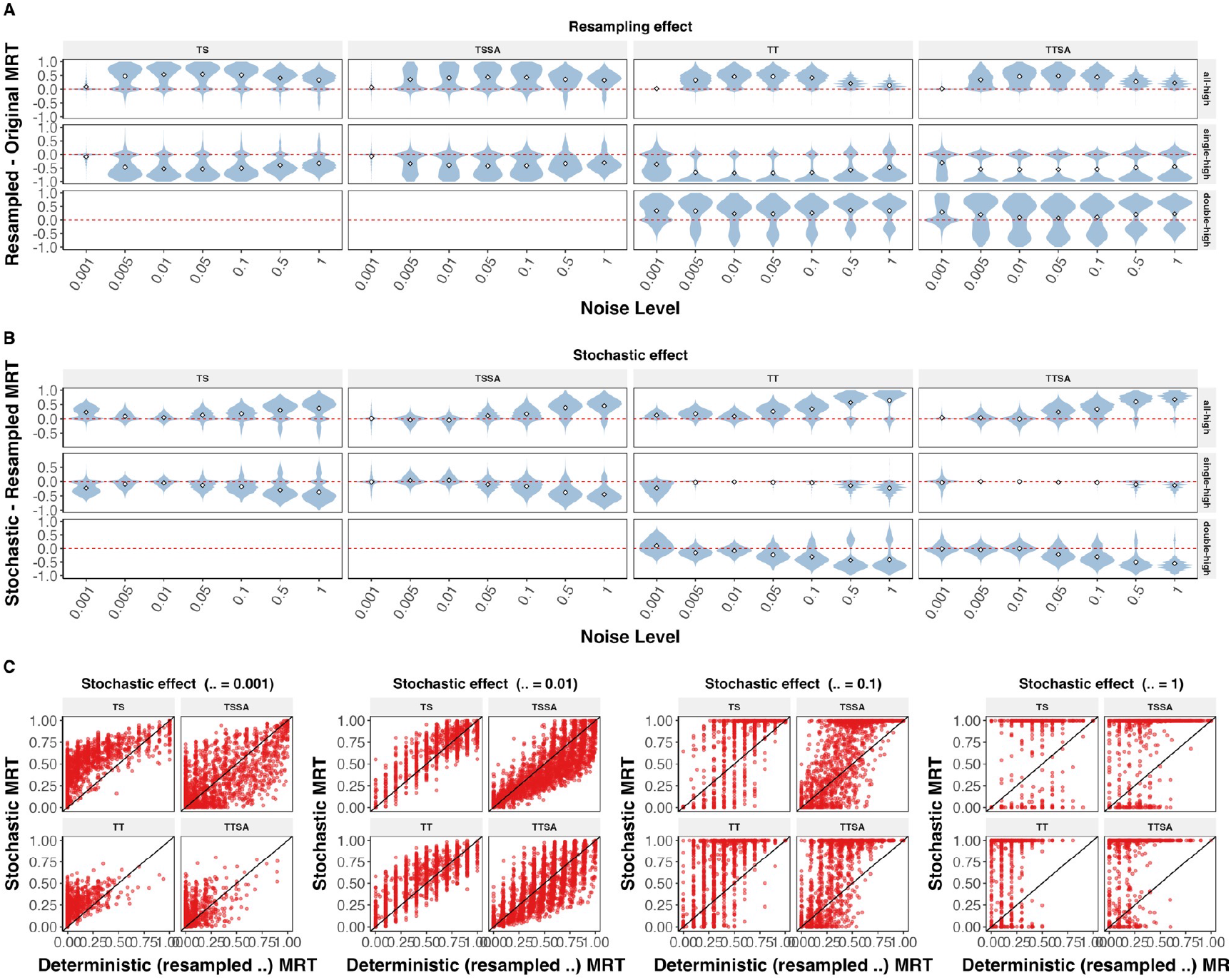
**(A)** Resampling effect (as in Figure 4B), collapsed to a single per-parameter-set difference and shown across all noise levels, faceted by state class (all-high/single-high/double-high) and network (TS, TSSA, TT, TTSA). **(B)** Same, for the stochastic effect. **(C)** Paramwise scatter of resampled-*λ* deterministic vs. stochastic MRT of the all-high state, all four networks, at each of four noise levels (0.001, 0.01, 0.1, 1.0).

**Figure S5:**
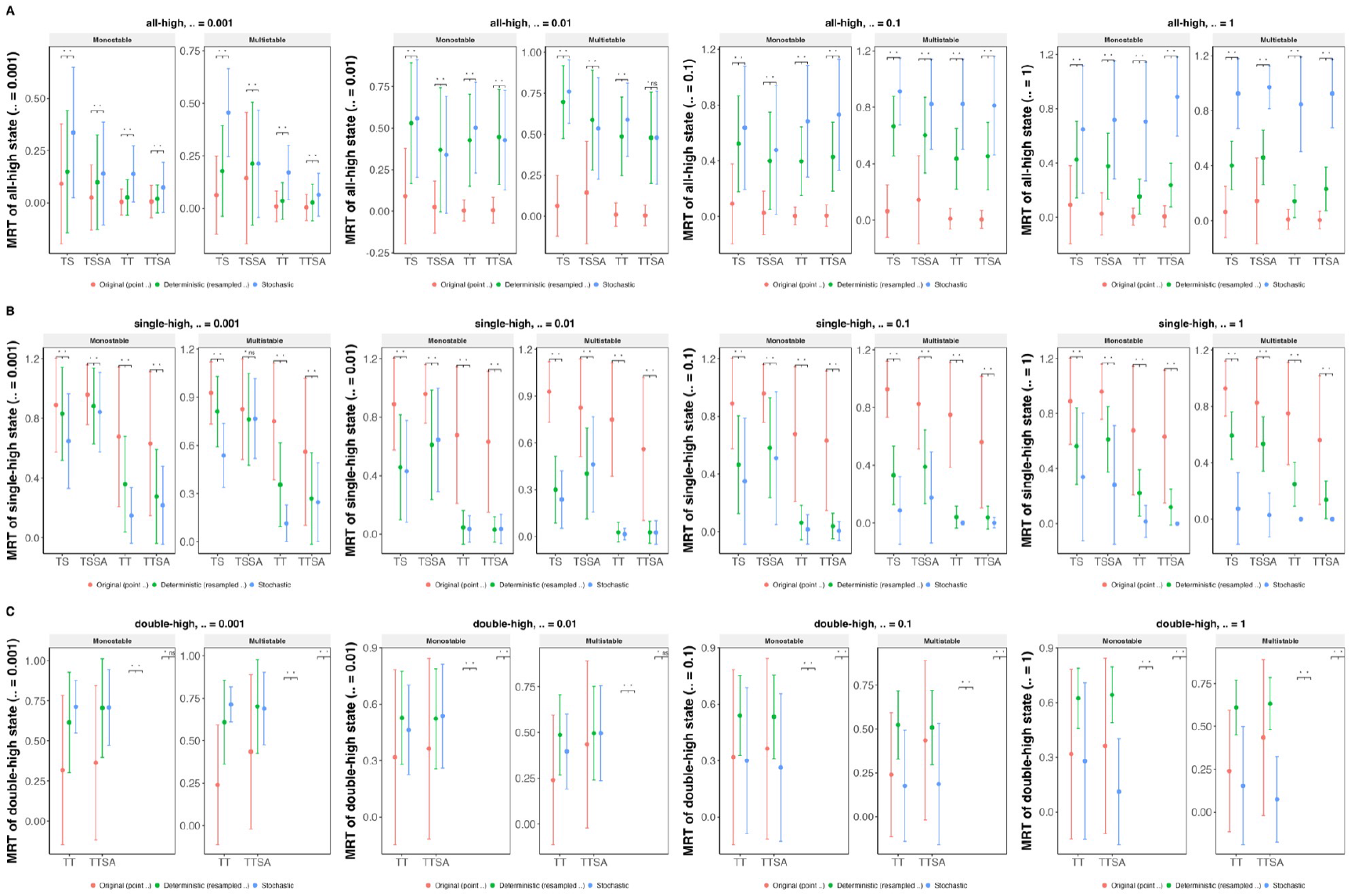
All-high **(A)**, single-high **(B)**, and double-high **(C)** state MRT comparisons (as in Figure 4D: original/resampled/stochastic, faceted by stability class), repeated at four noise levels (columns: 0.001, 0.01, 0.1, 1.0), for TS, TSSA, TT, TTSA. 2-node networks (TS, TSSA) have no double-high state distinct from all-high, so they are absent from that row.

**Figure S6:**
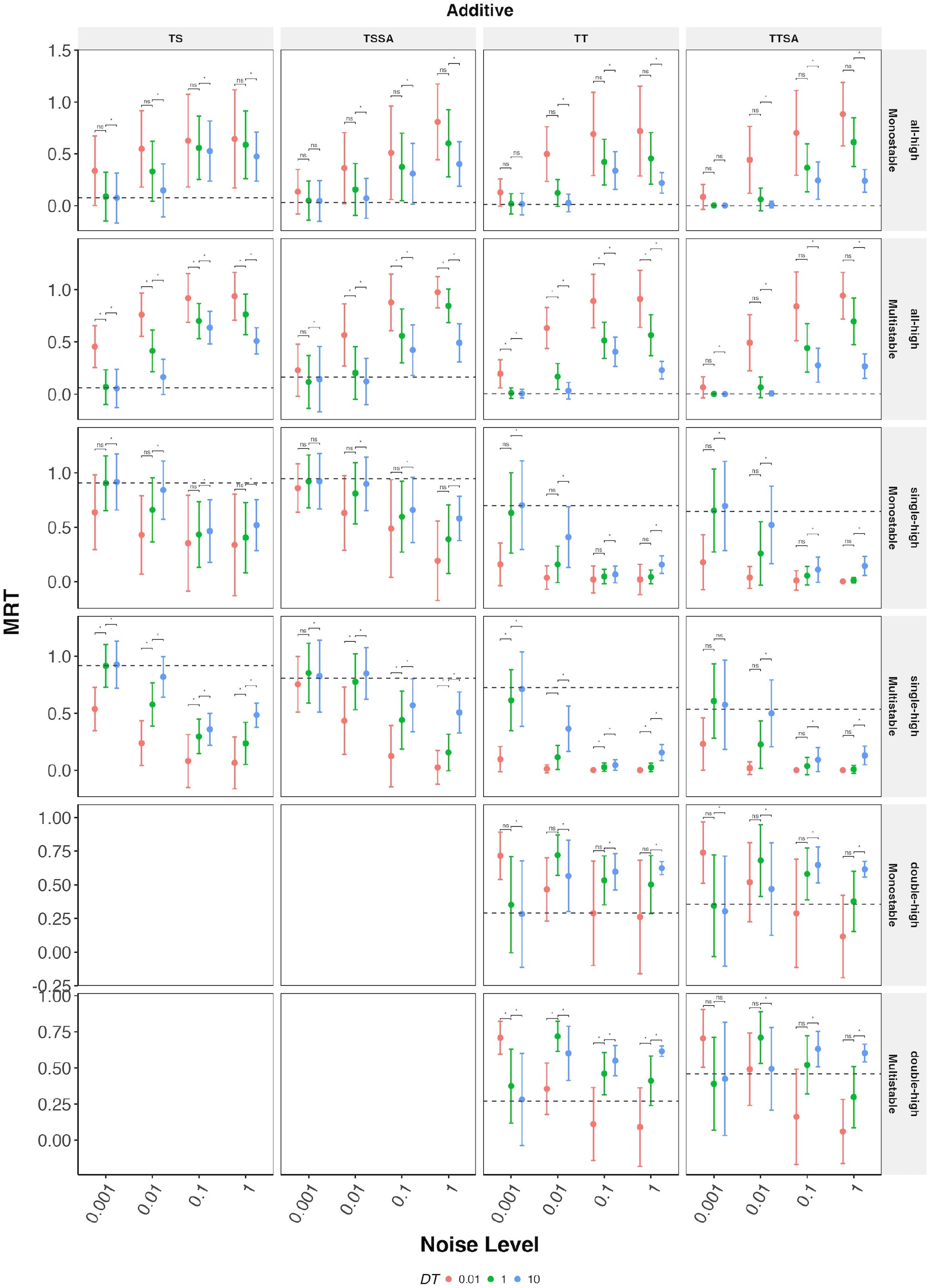
Comparison of MRTs for additive noise simulated at different values of *dt*

**Figure S7:**
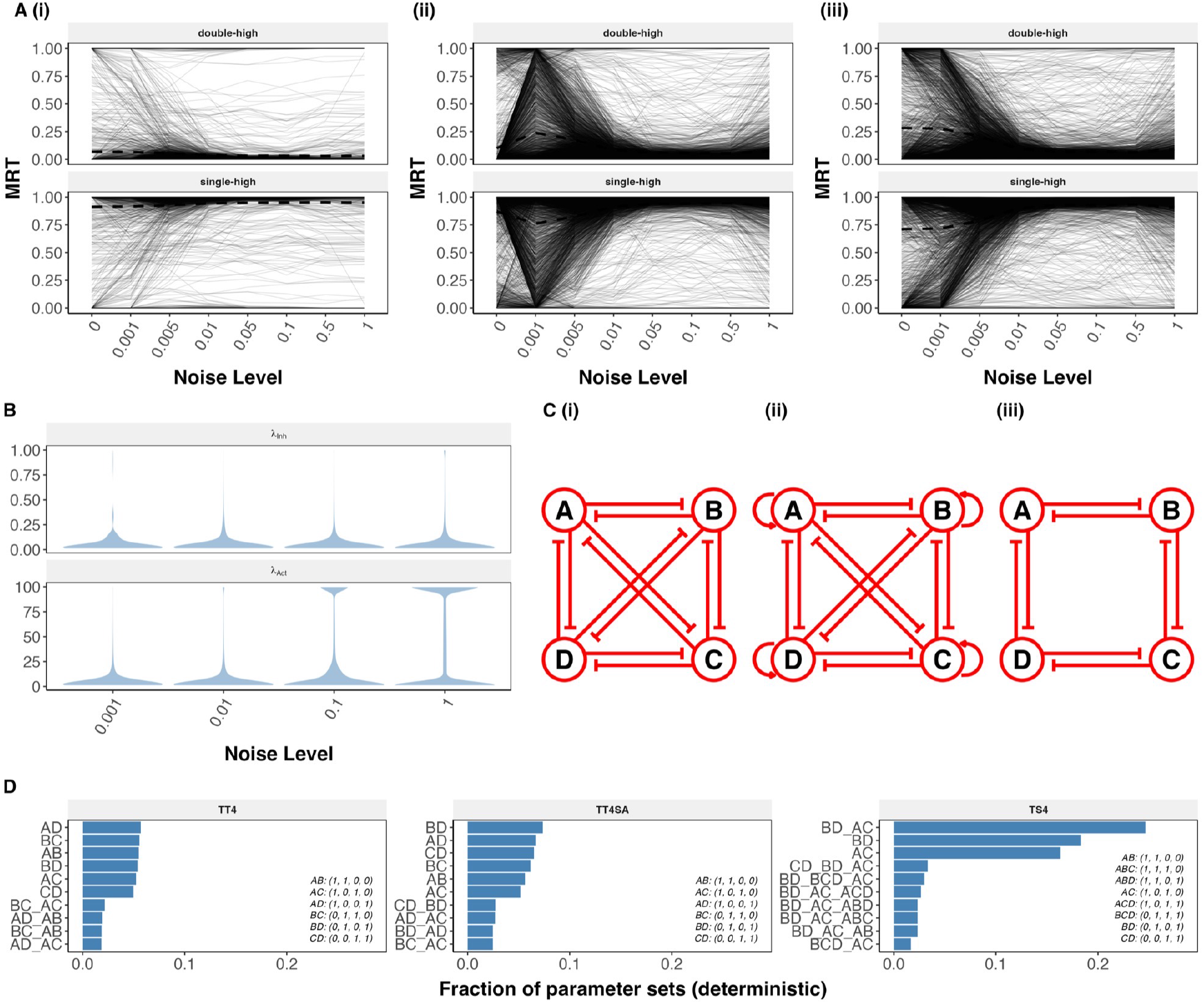
**(A)** MRT vs. noise level (as in Figure 5A, each showing a random sample of 100 parameter-set trajectories with the mean across the full population) for TS, TSSA, and TT; single-high and double-high state classes. For TS and TSSA, whose fully-active state is intrinsically “all-high” rather than a distinct “double-high” state, this facet is relabeled “double-high” for consistency with the other panels, but shows the same all-high data. **(B)** *λ* distributions (as in Figure 4A) under Multiplicative noise. **(C)** Network topology diagrams for TT4, TT4SA, and TS4. **(D)** Deterministic attractor-combination frequency (as in Fig. 1C, but from the noise-simulation pipeline’s zero-noise runs rather than RACIPE’s own steady-state search) for TT4, TT4SA, and TS4.

**Figure S8:**
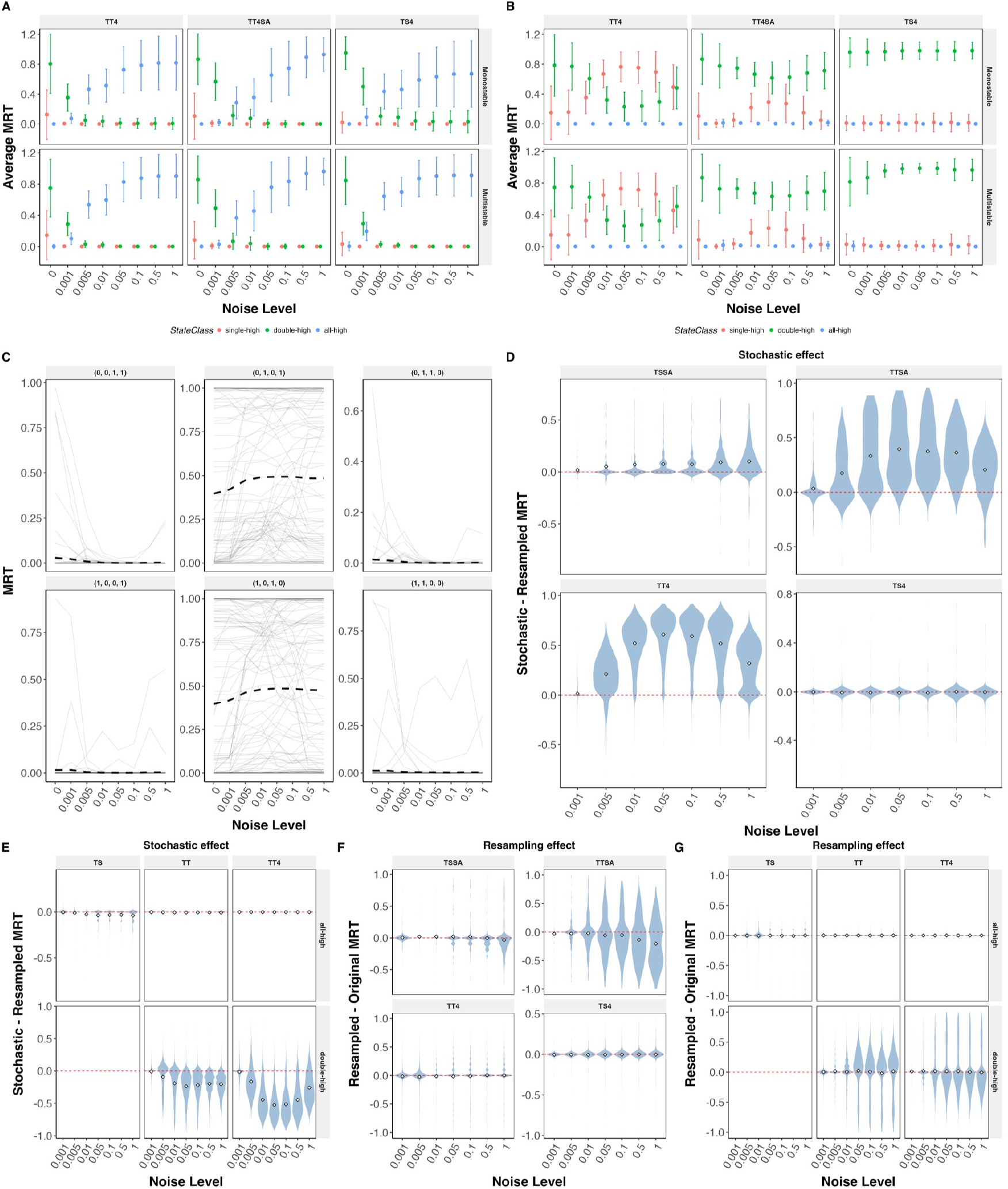
**(A, B)** MRT vs. noise level for TT4, TT4SA, and TS4; single-high, double-high, and all-high states, faceted by stability class, under Additive (A) and Multiplicative (B) noise. **(C)** State-wise MRT vs. noise level for each of TS4’s six double-high states individually (Multiplicative noise), showing that the same subset of states dominates at every noise level; grey lines show a random sample of 100 parameter-set trajectories, with the mean across the full parameter population shown separately. **(D)** Stochastic effect for the single-high state, across noise levels, for TSSA, TTSA, TT4, and TS4. **(E)** Stochastic effect for the all-high and double-high states, for TS, TT, and TT4. **(D**′, **E**′) Same networks/state classes as D and E, respectively, but showing the resampling effect instead of the stochastic effect.

**Figure S9:**
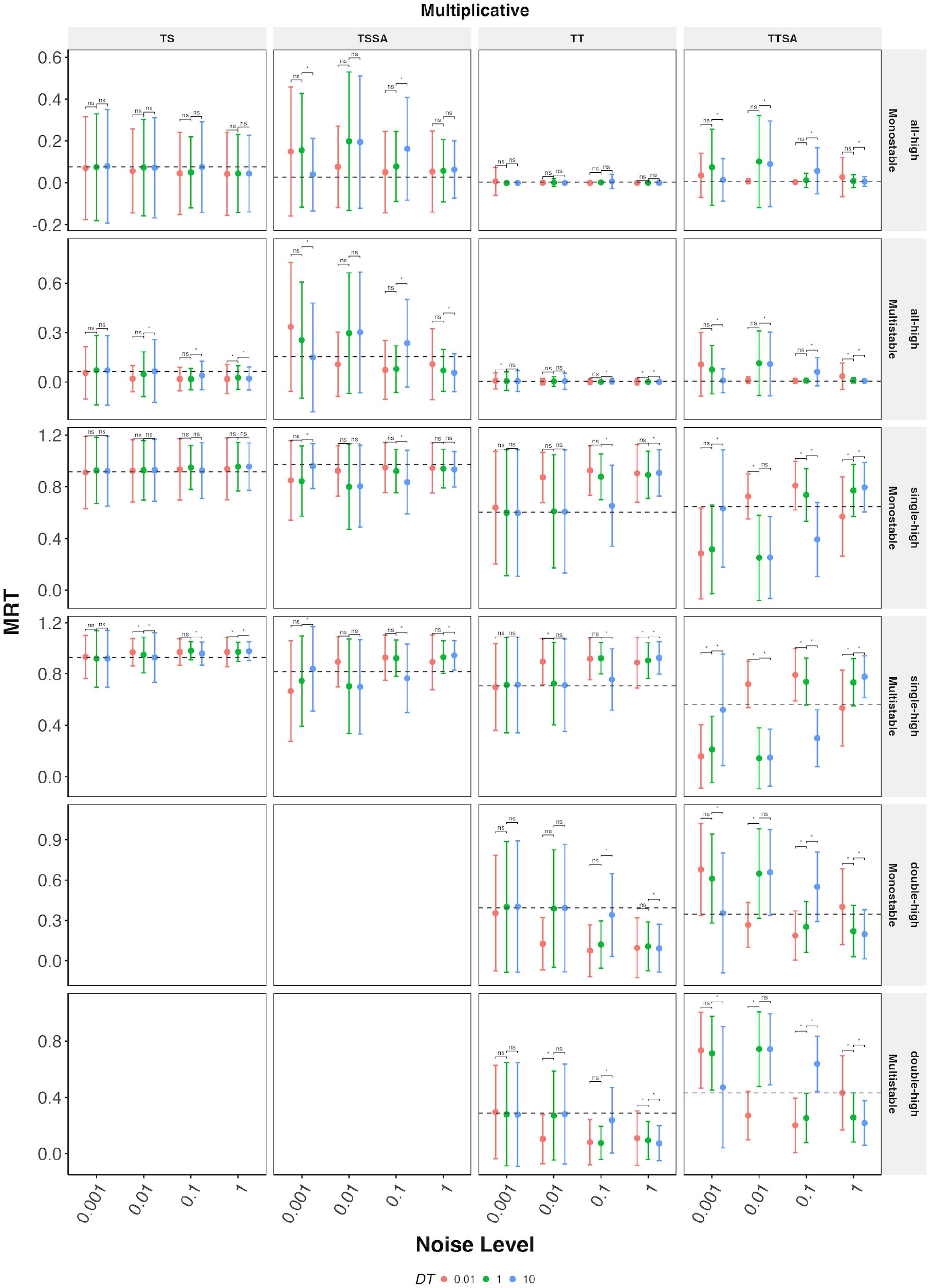
Comparison of MRTs for multiplicative noise simulated at different values of *dt*

**Figure S10:**
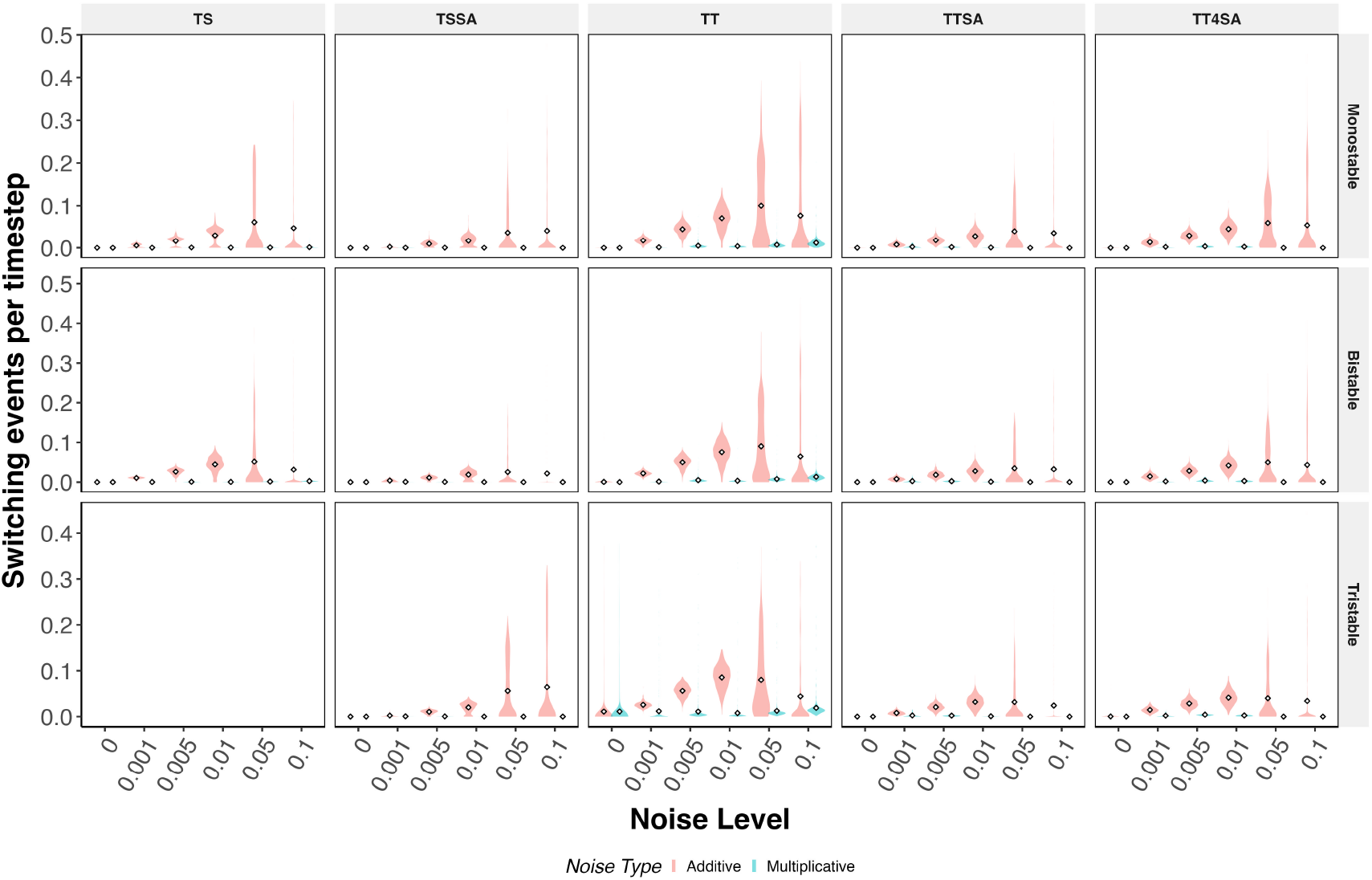
Switching-event rates generalize across networks and stability classes. Distribution of switching-event counts per timestep by noise level (Additive vs. Multiplicative), for TS, TSSA, TT, TTSA, and TT4SA, faceted by network (columns) and deterministic stability class (rows). TS has no Tristable class (a 2-node network cannot support 3+ distinct stable states), so that facet is empty.

**Figure S11:**
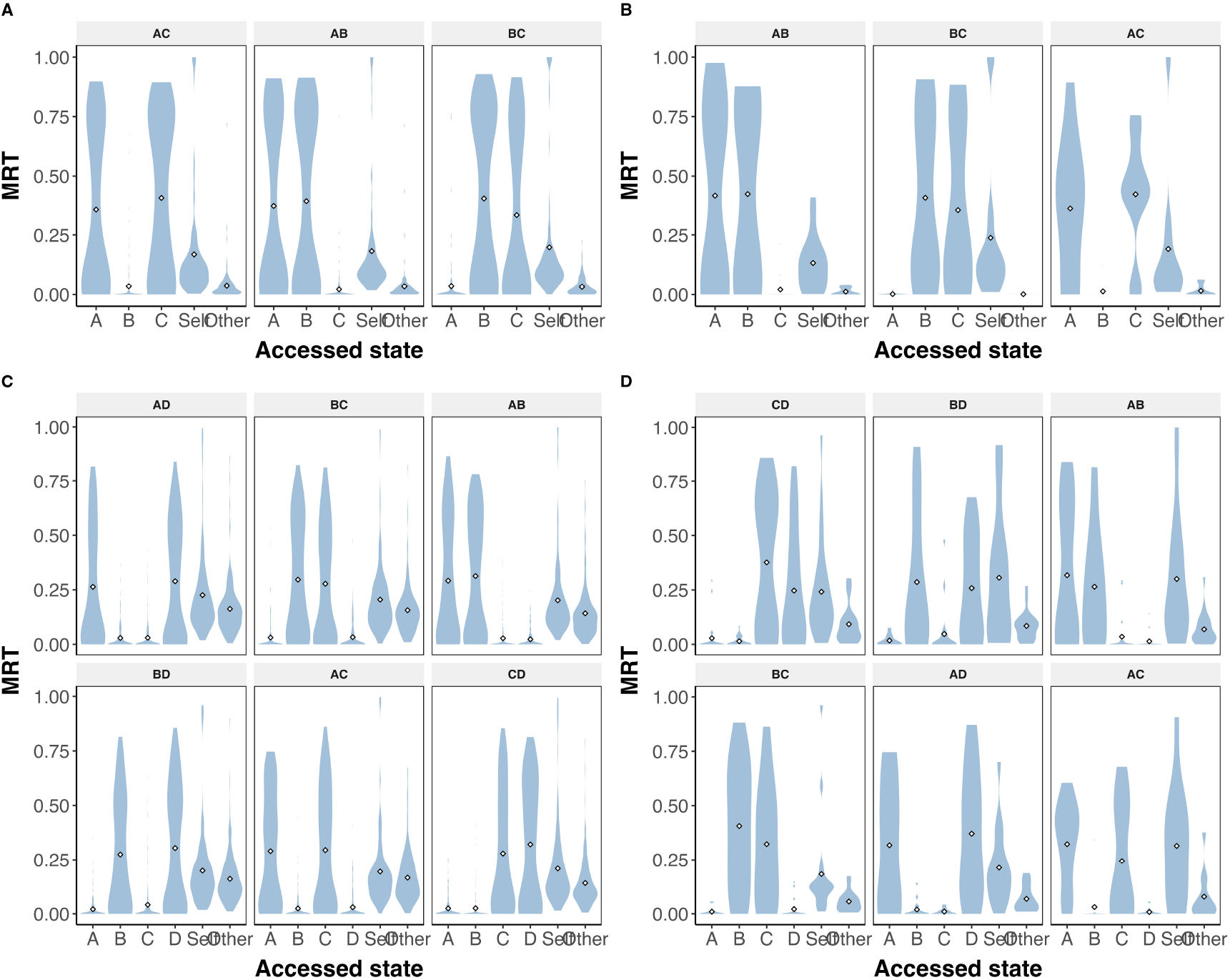
Nearest-fate resolution holds for every individual double-high attractor. Per-parameter MRT distribution at noise = 0.01, shown separately for each monostable double-high attractor combination (one facet per combination), for **(A)** TT, **(B)** TTSA, **(C)** TT4, and **(D)** TT4SA.

**Figure S12:**
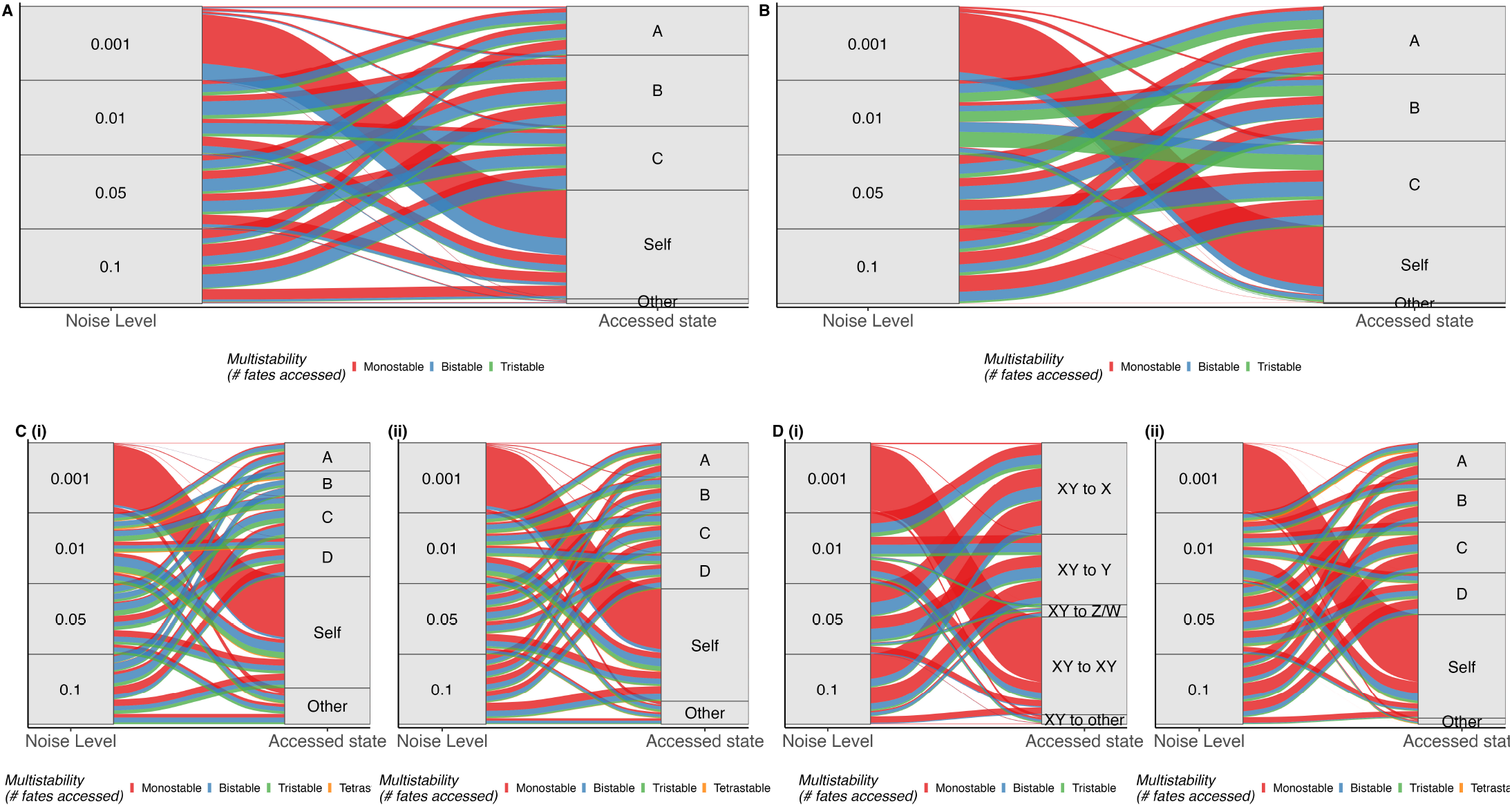
Noise-level → fate alluvials for the bistable double-high classes. Noise-level → outcome alluvial (as in Figure 6E), colored by the multistability classification from Figure 6C, for the bistable double-high stability class: **(A)** TT and **(B)** TTSA, non-mirror only (mirror bistability is structurally impossible for these 3-node networks). **(C)** TT4, mirror **(i)** and non-mirror **(ii)** shown separately. **(D)** TT4SA, monostable double-high class **(i)** (as in Figure 6E, for this network) and bistable double-high class **(ii)**, with mirror and non-mirror populations pooled (TT4SA’s mirror-bistable population alone is too sparse, ≈13 parameter sets, to interpret on its own).

**Figure S13:**
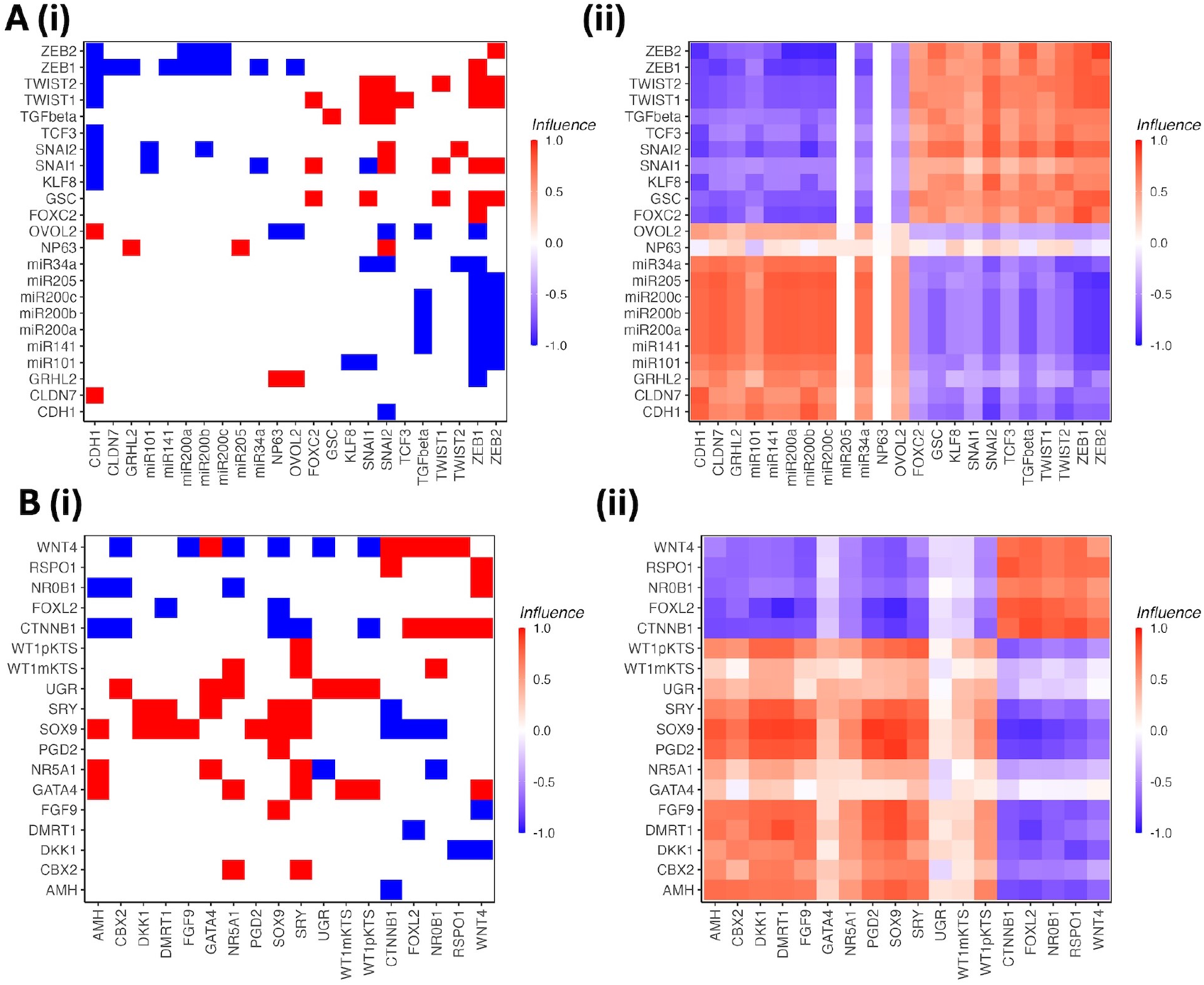
Adjacency and influence matrices for the EMT and Gonadal networks. A) EMT network (23 nodes). (i) Direct adjacency matrix showing the signed regulatory interactions between node pairs: red indicates activation (+1), blue indicates inhibition (− 1), and white indicates no edge. Nodes are ordered by team membership, with the epithelial team in the lower-left block and the mesenchymal team in the upper-right block. (ii) Influence matrix computed as described in the Methods, showing the net cumulative effect between each pair of nodes across all path lengths up to *l*_max_ = 10. Positive (red) values indicate net co-regulation; negative (blue) values indicate net counter-regulation. The block structure of the influence matrix confirms the two-team organisation of the network. B) Gonadal network (19 nodes), displayed in the same format as A, with the Sertoli-fate team and the Granulosa-fate team forming the two blocks.

**Figure S14:**
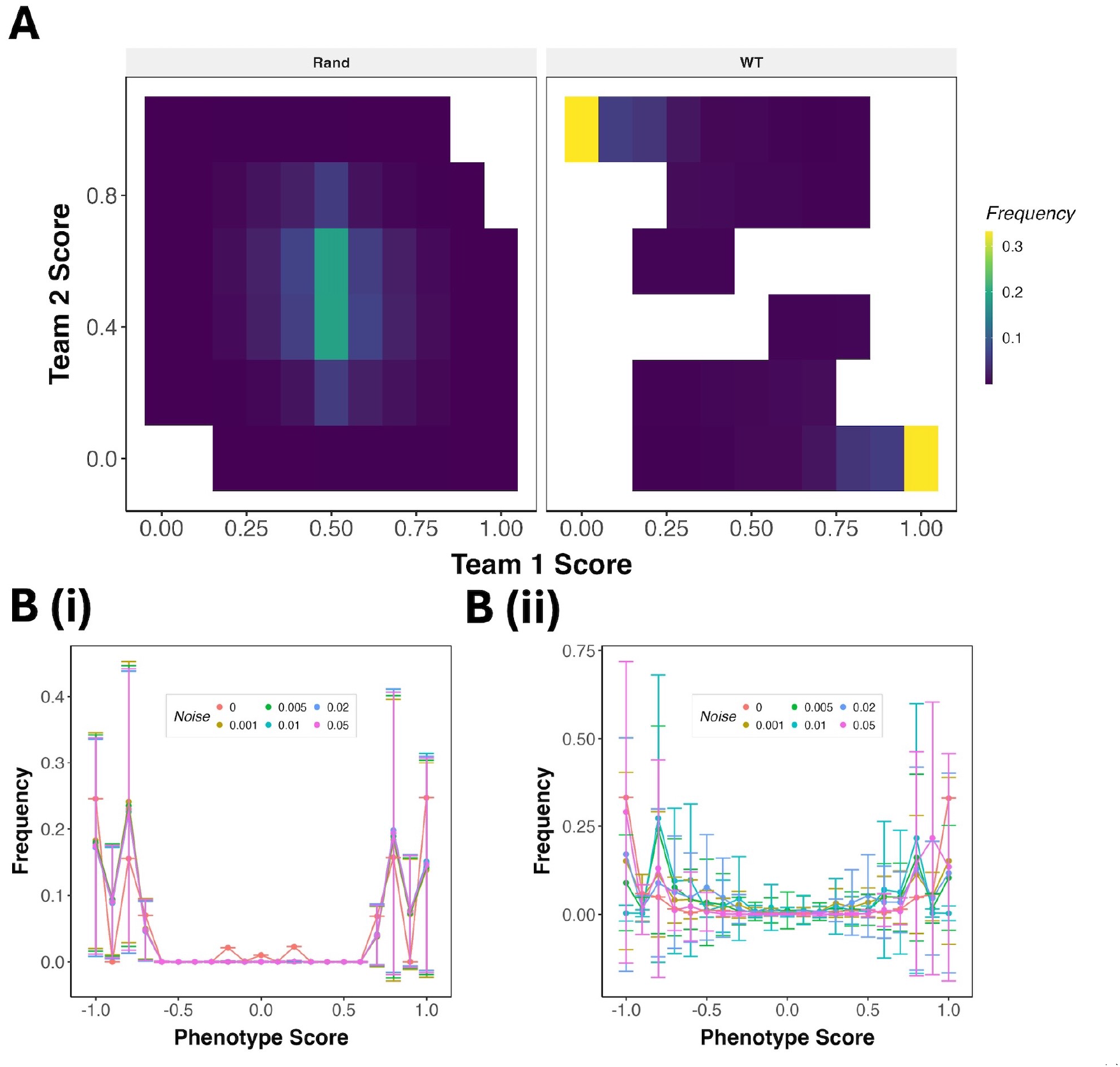
Effect of multiplicative and stationary noise on phenotypic distributions in biological networks. A) Team score density maps for the Gonadal network under additive noise at *σ* = 0.05, displayed as in Figure 5B. Left panel (“Rand”) shows the team score distribution sampled from stochastic simulation trajectories; right panel (“WT”) shows the distribution obtained from the steady states of the unperturbed network. B–C) Relative frequency of phenotypic scores across 1000 simulation trajectories for the EMT network (i) and Gonadal network (ii) under multiplicative noise (B) and stationary noise (C), shown at the same noise levels as in Figure 7C. Colours indicate noise level as shown in the legend. Trajectories are initiated from steady states of the unperturbed network as in Figure 7.

## S8 Supplementary Tables

**Table S1:**
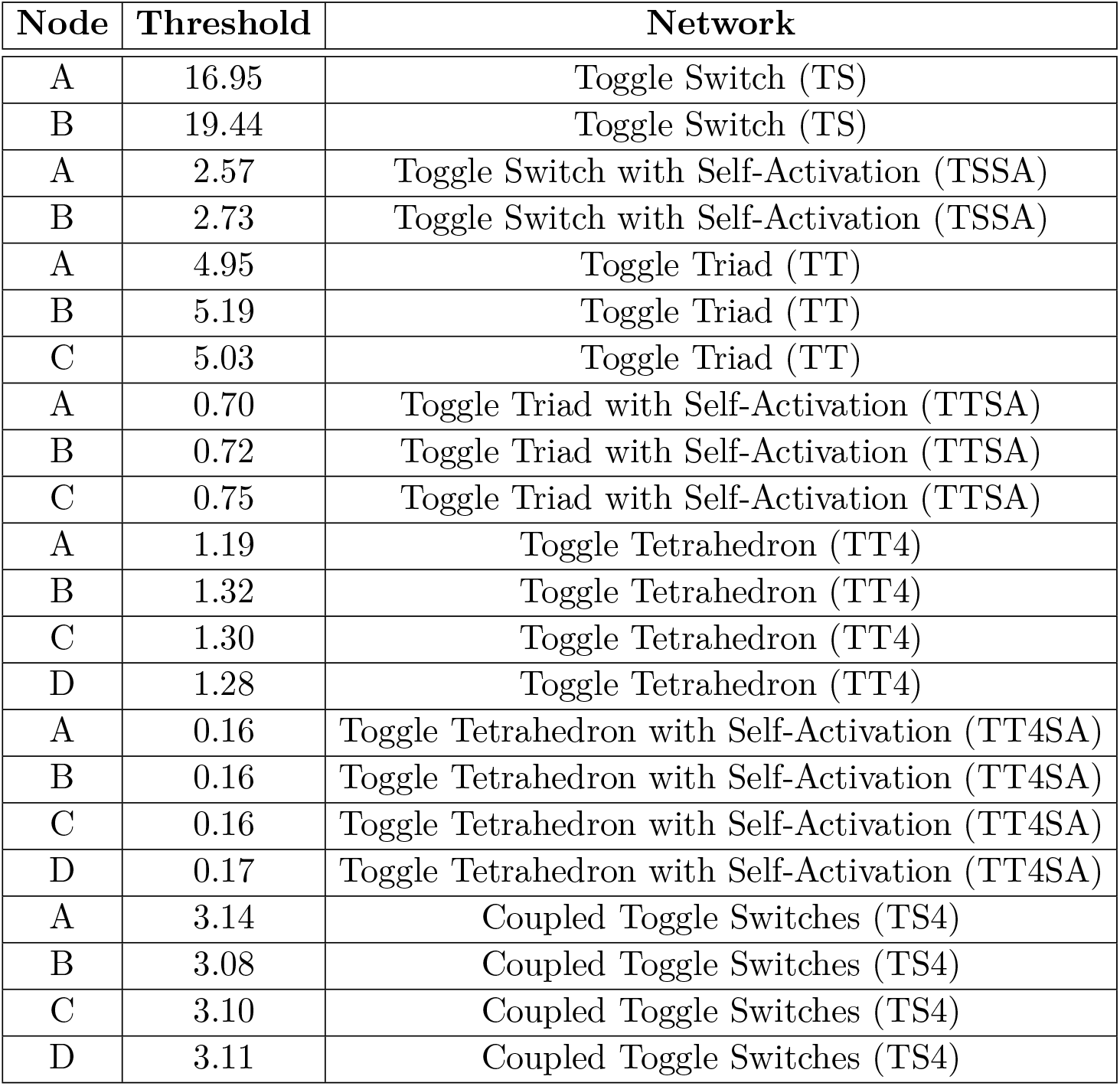
Threshold values used to discretize node expressions for different states in small cell-fate network motifs.

| Node | Threshold | Network |
| --- | --- | --- |
| A | 16.95 | Toggle Switch (TS) |
| B | 19.44 | Toggle Switch (TS) |
| A | 2.57 | Toggle Switch with Self-Activation (TSSA) |
| B | 2.73 | Toggle Switch with Self-Activation (TSSA) |
| A | 4.95 | Toggle Triad (TT) |
| B | 5.19 | Toggle Triad (TT) |
| C | 5.03 | Toggle Triad (TT) |
| A | 0.70 | Toggle Triad with Self-Activation (TTSA) |
| B | 0.72 | Toggle Triad with Self-Activation (TTSA) |
| C | 0.75 | Toggle Triad with Self-Activation (TTSA) |
| A | 1.19 | Toggle Tetrahedron (TT4) |
| B | 1.32 | Toggle Tetrahedron (TT4) |
| C | 1.30 | Toggle Tetrahedron (TT4) |
| D | 1.28 | Toggle Tetrahedron (TT4) |
| A | 0.16 | Toggle Tetrahedron with Self-Activation (TT4SA) |
| B | 0.16 | Toggle Tetrahedron with Self-Activation (TT4SA) |
| C | 0.16 | Toggle Tetrahedron with Self-Activation (TT4SA) |
| D | 0.17 | Toggle Tetrahedron with Self-Activation (TT4SA) |
| A | 3.14 | Coupled Toggle Switches (TS4) |
| B | 3.08 | Coupled Toggle Switches (TS4) |
| C | 3.10 | Coupled Toggle Switches (TS4) |
| D | 3.11 | Coupled Toggle Switches (TS4) |

**Table S2:** Attractor frequencies across all network motifs. Each sub-panel lists attractors (monostable or multistable, written as state-tuples joined by “_”) and their frequencies in RACIPE.

| **(a) Toggle Switch (TS)** |  | **(b) TS Self-Activation (TSSA)** |  |
| --- | --- | --- | --- |
| Attractor | Freq. | Attractor | Freq. |
| (0,1) | 0.3744 | (0,1)_(1,0) | 0.4865 |
| (1,0) | 0.3318 | (0,1) | 0.1357 |
| (0,1)_(1,0) | 0.1858 | (1,0) | 0.1243 |
| (1,1) | 0.0573 | (0,0)_(0,1)_(1,0) | 0.0682 |
|  |  | (0,1)_(1,0)_(1,1) | 0.0369 |
|  |  | (1,0)_(1,1) | 0.0284 |
|  |  | (0,1)_(1,1) | 0.0254 |
|  |  | (0,0)_(0,1) | 0.0187 |
|  |  | (0,0)_(1,0) | 0.0184 |

**(g) Toggle Triad (TT)**
| Attractor | Freq. |
| --- | --- |
| (0,0,1) | 0.1312 |
| (0,1,0) | 0.1241 |
| (1,0,0) | 0.1184 |
| (0,0,1)_(0,1,0) | 0.0694 |
| (0,1,0)_(1,0,0) | 0.0638 |
| (0,0,1)_(1,0,0) | 0.0632 |
| (1,0,1) | 0.0567 |
| (1,1,0) | 0.0559 |
| (0,1,1) | 0.0535 |
| (0,0,1)_(0,1,0)_(1,0,0) | 0.0206 |
| (0,1,0)_(1,0,1) | 0.0187 |
| (1,0,0)_(1,0,1) | 0.0181 |
| (0,1,0)_(0,1,1) | 0.0171 |
| (1,0,0)_(1,1,0) | 0.0169 |
| (0,1,1)_(1,0,0) | 0.0166 |
| (0,0,1)_(1,1,0) | 0.0161 |
| (0,1,0)_(1,1,0) | 0.0146 |
| (0,0,1)_(1,0,1) | 0.0138 |
| (0,0,1)_(0,1,1) | 0.0136 |
| (0,1,1)_(1,1,0) | 0.0078 |
| (1,0,1)_(1,1,0) | 0.0078 |
| (0,1,1)_(1,0,1) | 0.0076 |
| (1,0,1)_(1,0,1) | 0.0055 |
| (1,1,0)_(1,1,0) | 0.0042 |
| (0,1,0)_(1,0,0)_(1,0,1) | 0.0039 |
| (0,0,1)_(1,0,0)_(1,1,0) | 0.0037 |
| (0,1,1)_(0,1,1) | 0.0034 |
| (0,0,1)_(0,1,0)_(1,1,0) | 0.0028 |

**(j) Coupled Toggle Switches (TS4)**
| Attractor | Freq. |
| --- | --- |
| (0,1,0,1)_(1,0,1,0) | 0.5244 |
| (1,0,1,0) | 0.1313 |
| (0,1,0,1) | 0.1251 |
| (0,1,0,1)_(1,0,1,1) | 0.0112 |
| (1,0,1,0)_(1,1,0,1) | 0.0108 |
| (0,1,0,1)_(1,1,1,0) | 0.0096 |
| (0,1,1,1)_(1,0,1,0) | 0.0096 |
| (0,1,0,1)_(1,0,1,0)_(1,1,0,0) | 0.0093 |
| (0,0,1,1)_(0,1,0,1)_(1,0,1,0) | 0.0092 |
| (0,1,0,1)_(1,0,0,1)_(1,0,1,0) | 0.0092 |
| (0,1,0,1)_(0,1,1,0)_(1,0,1,0) | 0.0088 |
| (0,1,0,1)_(0,1,1,1)_(1,0,1,0) | 0.0079 |
| (0,1,0,1)_(1,0,1,0)_(1,0,1,1) | 0.0065 |
| (0,1,0,1)_(1,0,1,0)_(1,1,1,0) | 0.0063 |
| (0,1,0,1)_(1,0,1,0)_(1,1,0,1) | 0.0059 |
| (0,1,0,1)_(1,0,0,0) | 0.0049 |
| (0,1,0,0)_(1,0,1,0) | 0.0042 |
| (0,0,1,0)_(0,1,0,1) | 0.0041 |
| (0,1,0,1)_(0,1,1,0) | 0.0040 |
| (0,1,0,1)_(1,0,0,1) | 0.0040 |
| (0,1,0,1)_(1,1,0,0) | 0.0040 |
| (1,0,0,1)_(1,0,1,0) | 0.0040 |
| (0,1,0,1)_(1,0,0,0)_(1,0,1,0) | 0.0038 |
| (0,0,0,1)_(1,0,1,0) | 0.0037 |
| (0,1,1,0)_(1,0,1,0) | 0.0037 |
| (1,0,1,0)_(1,1,0,0) | 0.0037 |
| (0,0,1,1)_(1,0,1,0) | 0.0036 |
| (0,1,0,1)_(0,1,0,1)_(1,0,1,0) | 0.0035 |
| (0,0,1,1)_(0,1,0,1) | 0.0034 |
| (0,0,0,1)_(0,1,0,1)_(1,0,1,0) | 0.0033 |
| (0,1,0,0)_(0,1,0,1)_(1,0,1,0) | 0.0032 |
| (1,0,1,0)_(1,1,1,0) | 0.0032 |

| Attractor | Freq. |
| --- | --- |
| (0,0,1)_(0,1,0)_(1,0,0) | 0.1449 |
| (0,0,1)_(0,1,0) | 0.0459 |
| (0,0,1)_(1,0,0) | 0.0444 |
| (0,1,0)_(1,0,0) | 0.0444 |
| (1,0,0) | 0.0362 |
| (0,0,1) | 0.0359 |
| (0,1,0) | 0.0330 |
| (0,0,1)_(0,1,0)_(1,0,0)_(1,0,1) | 0.0228 |
| (1,0,0)_(1,0,1) | 0.0227 |
| (0,0,1)_(0,1,0)_(0,1,1)_(1,0,0) | 0.0220 |
| (0,0,1)_(0,1,1) | 0.0215 |
| (0,1,0)_(0,1,1) | 0.0213 |
| (0,1,0)_(1,1,0) | 0.0203 |
| (0,0,1)_(0,1,0)_(1,0,0)_(1,1,0) | 0.0201 |
| (0,0,1)_(1,0,1) | 0.0198 |
| (1,0,0)_(1,1,0) | 0.0195 |
| (0,1,1) | 0.0188 |
| (1,0,1) | 0.0183 |
| (0,0,1)_(0,1,0)_(0,1,1) | 0.0166 |
| (1,1,0) | 0.0161 |
| (0,0,1)_(1,0,0)_(1,0,1) | 0.0140 |
| (0,1,0)_(1,0,0)_(1,1,0) | 0.0138 |
| (1,0,0)_(1,0,1)_(1,1,0) | 0.0135 |
| (0,1,0)_(0,1,1)_(1,1,0) | 0.0132 |
| (0,0,1)_(0,1,1)_(1,0,1) | 0.0117 |
| (1,0,1)_(1,1,0) | 0.0116 |
| (0,0,1)_(0,1,0)_(1,0,1) | 0.0111 |
| (0,0,1)_(0,1,1)_(1,0,0) | 0.0110 |
| (0,1,1)_(1,1,0) | 0.0102 |
| (0,0,1)_(0,1,0)_(1,1,0) | 0.0101 |
| (0,1,1)_(1,0,1) | 0.0100 |
| (0,1,0)_(0,1,1)_(1,0,0) | 0.0098 |
| (0,1,0)_(1,0,0)_(1,0,1) | 0.0097 |
| (0,0,1)_(1,0,0)_(1,1,0) | 0.0096 |
| (0,0,1)_(0,1,0)_(0,1,1)_(1,0,0)_(1,0,1) | 0.0073 |
| (0,1,1)_(1,0,0) | 0.0071 |
| (0,0,1)_(0,1,0)_(1,0,0)_(1,0,1)_(1,1,0) | 0.0070 |
| (0,0,1)_(0,1,0)_(0,1,1)_(1,0,0)_(1,1,0) | 0.0064 |
| (0,0,1)_(1,1,0) | 0.0062 |
| (0,1,1)_(1,0,1)_(1,1,0) | 0.0057 |
| (0,1,0)_(1,0,1) | 0.0055 |
| (0,0,1)_(0,1,1)_(1,1,0) | 0.0045 |
| (0,1,1)_(1,0,0)_(1,0,1) | 0.0044 |
| (0,0,1)_(0,1,1)_(1,0,0)_(1,1,0) | 0.0043 |
| (0,0,1)_(1,0,1)_(1,1,0) | 0.0043 |
| (0,1,0)_(0,1,1)_(1,0,0)_(1,0,1) | 0.0043 |
| (0,0,1)_(0,1,0)_(0,1,1)_(1,0,1) | 0.0041 |
| (0,0,1)_(0,1,1)_(1,0,0)_(1,0,1) | 0.0040 |
| (0,0,1)_(0,1,0)_(1,0,1)_(1,1,0) | 0.0035 |
| (0,1,0)_(1,0,0)_(1,0,1)_(1,1,0) | 0.0032 |
| (0,1,0)_(1,0,1)_(1,1,0) | 0.0031 |
| (0,0,1)_(0,1,0)_(0,1,1)_(1,0,0)_(1,0,1)_(1,1,0) | 0.0030 |
| (0,1,0)_(0,1,1)_(1,0,1) | 0.0030 |
| (0,1,1)_(1,0,0)_(1,1,0) | 0.0030 |
| (0,0,1)_(0,1,1)_(1,0,1)_(1,1,0) | 0.0029 |
| (0,0,1)_(1,0,0)_(1,0,1)_(1,1,0) | 0.0029 |
| (0,1,0)_(0,1,1)_(1,0,1)_(1,1,0) | 0.0028 |
| (0,0,1)_(0,1,0)_(0,1,1)_(1,1,0) | 0.0027 |
| (0,1,0)_(0,1,1)_(1,0,0)_(1,1,0) | 0.0026 |
| (0,1,1)_(1,0,0)_(1,0,1)_(1,1,0) | 0.0026 |
| (1,1,0)_(1,1,0) | 0.0021 |
| (1,0,1)_(1,1,1) | 0.0018 |
| (0,1,0)_(0,1,1)_(0,1,1) | 0.0017 |
| (0,1,0)_(0,1,1)_(1,0,0)_(1,0,1)_(1,1,0) | 0.0017 |
| (0,1,1)_(0,1,1) | 0.0016 |
| (1,0,1)_(1,0,1) | 0.0015 |
| (0,0,1)_(1,0,1)_(1,0,1) | 0.0014 |
| (0,1,1)_(1,1,1) | 0.0013 |
| (1,0,0)_(1,0,1)_(1,0,1) | 0.0013 |
| (0,0,1)_(0,1,1)_(0,1,1) | 0.0012 |
| (0,0,1)_(0,1,1)_(1,0,0)_(1,0,1)_(1,1,0) | 0.0012 |
| (0,1,0)_(0,1,1)_(1,1,0)_(1,1,0) | 0.0012 |
| (0,1,0)_(1,1,0)_(1,1,0) | 0.0012 |
| (0,0,1)_(0,1,0)_(0,1,0)_(1,0,0) | 0.0011 |
| (0,0,1)_(0,1,0)_(1,0,0)_(1,0,0) | 0.0011 |
| (0,0,1)_(0,1,0)_(1,1,0)_(1,1,0) | 0.0011 |
| (1,0,0)_(1,1,0)_(1,1,0) | 0.0011 |
| (1,1,0)_(1,1,1) | 0.0011 |
| (0,0,0)_(0,0,1)_(0,1,0)_(1,0,0) | 0.0010 |
| (0,1,1)_(0,1,1)_(1,0,1) | 0.0010 |
| (0,1,1)_(0,1,1)_(1,1,0) | 0.0010 |
| (0,1,1)_(1,0,1)_(1,1,1) | 0.0010 |
| (1,0,0)_(1,0,1)_(1,1,0)_(1,1,1) | 0.0010 |
| (1,0,1)_(1,1,0)_(1,1,0) | 0.0010 |
| (0,0,1)_(0,0,1) | 0.0009 |
| (0,0,1)_(0,1,0)_(1,0,1)_(1,0,1) | 0.0009 |
| (0,1,1)_(0,1,1)_(1,0,0) | 0.0009 |
| (0,1,1)_(1,1,0)_(1,1,1) | 0.0009 |
| (1,0,1)_(1,0,1)_(1,1,0) | 0.0009 |

## Notes

https://github.com/askhari139/Regulatory-noise-model

### Competing Interest Statement

The authors have declared no competing interest.

